# Conserved lifestyle-associated chromosome architectures across mammalian symbionts

**DOI:** 10.64898/2026.07.20.739713

**Authors:** Tobias Viehboeck, Nicole Krause, Philipp M. Weber, Éanna O. Shea, Valentin Noetzel, Nika Pende, Hanna Gölles-Kirth, Nelle Varoquaux, Frédéric Boccard, Ivan Junier, Virginia S. Lioy, Silvia Bulgheresi

## Abstract

Bacterial chromosome biology has largely focused on bacteria that are either free-living or facultatively associated with eukaryotes. Therefore, it is not known how obligate animal symbionts organize their chromosomes. Here, we studied the chromosome organization of three species of multicellular *Neisseriaceae* that colonize the oral cavity of mammals, *Alysiella filiformis, Simonsiella muelleri* and *Conchiformibius steedae*. DNA fluorescence in situ hybridization showed that – irrespective of their ploidy – their chromosomes are longitudinally configured with the origin of DNA replication consistently localized at their host-attached poles throughout the cell cycle. Immunolocalization, ChIP-seq and EMSA implicated ParBS complexes in maintaining this stable chromosome orientation. Moreover, chromosome conformation capture across two species and three growth conditions further revealed conserved lifestyle-associated chromosome architectures, including planktonic-specific ParB-associated chromatin loops and condition-specific chromatin frontiers.

Together, our findings show that obligate mammalian symbionts maintain stable longitudinal chromosome orientations irrespective of ploidy while remodeling higher-order chromosome architecture according to physiological state. Distinct yet conserved chromosome architectures characterize exponential, stationary and surface-associated growth, indicating that bacterial genome folding reflects lifestyle and environmental context.

## INTRODUCTION

Despite the importance of chromosome orientation and conformation for cell functioning, we do not know whether they mediate or are affected by the colonization of solid surfaces, be them abiotic or biotic (Abbondanzieri et al., 2025). We therefore set out to study the chromosome organization of three multicellular longitudinally septating *Neisseriaceae* obligatorily thriving on the oral mucosa of mammals: a caprine strain of *Alysiella filiformis* (Simons, 1922; Tønjum, 2015; Nyongesa et al., 2022), a human strain of *Simonsiella muelleri* (Hedlund and Tønjum, 2015; Kaiser and Starzyk, 1973) and a canine strain of *Conchiformibius steedae* (Kuhn and Gregory, 1978; Xie and Yokota, 2005; Nyongesa et al., 2022; Fig. 1). Given that in all three species the pili are specifically localized at the bacterium-host interface (Kuhn and Gregory, 1978; Tønjum, 2015; Hedlund and Tønjum, 2015; Nyongesa et al., 2022), we will henceforth refer to the fimbriae-rich pole of *A. filiformis* rods as the proximal (P) pole and to the free pole as the distal pole. As for crescent-shaped *S. muelleri* and *C. steedae*, they adhere to the oral mucosa by both poles, and it is only their concave, ventral (proximal) side which faces the host to be fimbriated (Nyongesa et al., 2022).

**Figure 1.**
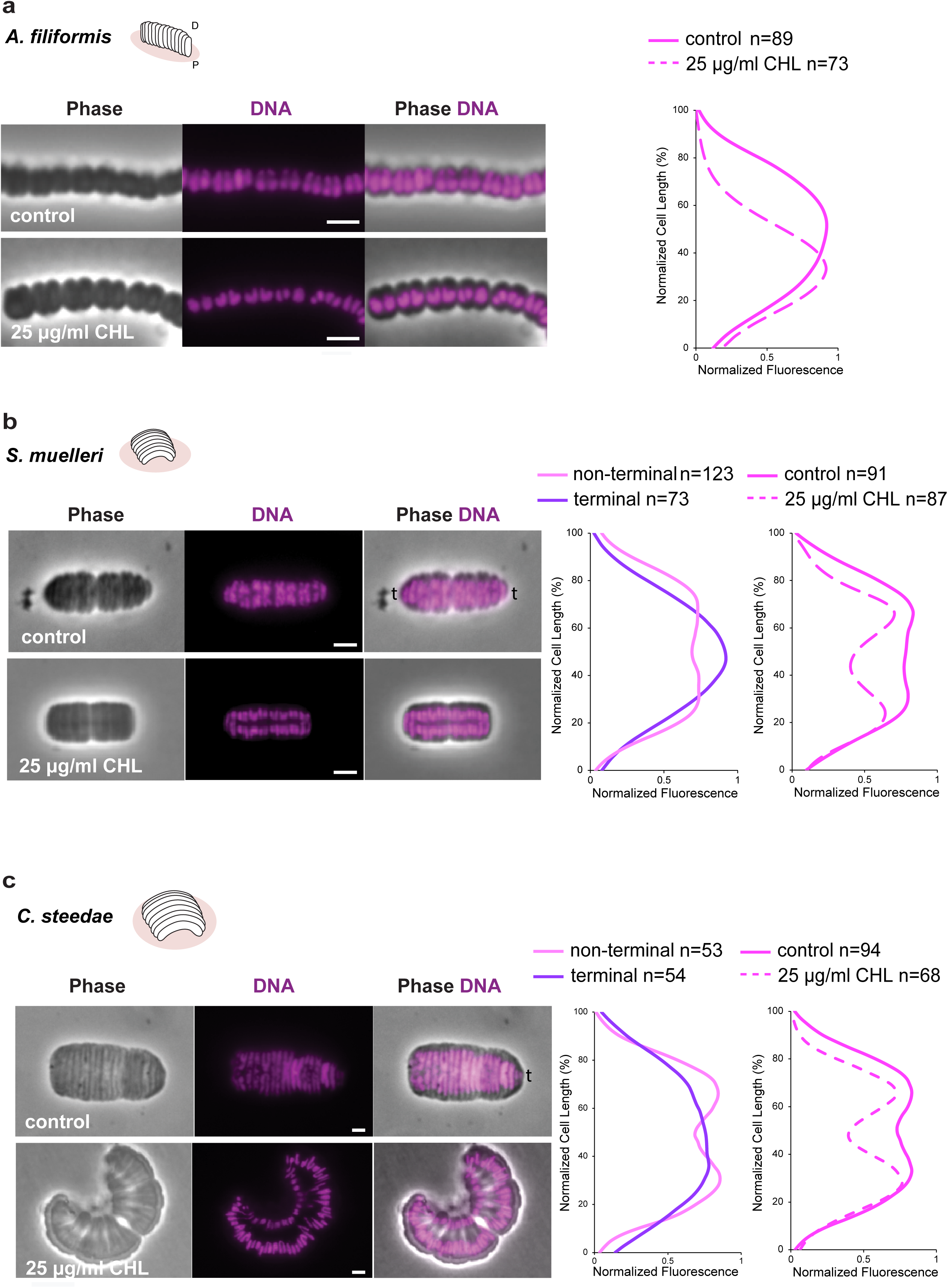
Nucleoid localization in translating or translationally halted *A. filiformis*, *S. muelleri* and *C. steedae* cells. (a) Phase contrast image of an *A. filiformis* filament (lateral view), corresponding DNA (Hoechst 3342, magenta) staining and overlay are displayed from left to right in untreated (control) or chloramphenicol-treated cells (25 µg/ml CHL) and normalized DNA fluorescence emitted by untreated (control, continuous line) or chloramphenicol-treated cells (dashed line) plotted against the long axis (%). (b) Phase contrast image of a *S. muelleri* filament (top view), corresponding epifluorescence microscope image of DNA staining (Hoechst 3342, magenta) and overlay are displayed from left to right in untreated (control) or chloramphenicol-treated cells (25 µg/ml CHL). Normalized DNA fluorescence emitted by non-terminal cells (magenta) or by the first and last cells of a filament (terminal cells, purple) plotted against the cell long axis (%). Normalized DNA fluorescence emitted by untreated (control, continuous line) or chloramphenicol-treated cells (dashed line) plotted against the cell long axis (%). (c) Phase contrast image of a *C. steedae* filament (top view), corresponding DNA staining (Hoechst 3342, magenta) and overlay are displayed from left to right in untreated (control) or chloramphenicol-treated cells (25 µg/ml CHL). Normalized DNA fluorescence emitted by non-terminal cells (magenta) or by the first and last cells of a filament (terminal cells, purple) plotted against the cell long axis (%). Normalized DNA fluorescence emitted by untreated (control, continuous line) or chloramphenicol-treated cells (dashed line) plotted against the cell long axis. D: distal. P: proximal side. t: terminal cell. All scale bars are 2 μm.

Although the chromosome biology of multicellular *Neisseriaceae* is still *terra incognita*, studies of model rods showed that free-living or facultatively host-associated bacteria may adopt two main spatial configurations (also referred to as orientations): left-*ori*-right (transverse) or *ori-ter* (longitudinal) (Toro and Shapiro, 2010; Wang and Lopis, 2013; Surovtsev and Jacobs-Wagner, 2018). In the transverse configuration, the origin of DNA replication (*ori*) and the terminus of DNA replication (a genomic region characterized by the presence of *dif* sites henceforth referred to as *ter*; Kleckner et al., 2014, Krogh et al., 2018) are at midcell, so that the left and right chromosome arms reside in opposite cell halves. In the longitudinal configuration, *ori* is located at one cell pole and *ter* at the opposite pole so that both arms (left and right) are parallel to the long axis of the cell. While model rods such as *Escherichia coli* and *Bacillus subtilis* can switch between transverse and longitudinal configuration depending, respectively, on the growth conditions or the cell cycle stage (Niki et al., 2000; Bates and Kleckner, 2005; Wang et al., 2014; Cass et al., 2016; Marbouty et al., 2015) the chromosome of vertically polarized bacteria^1^ (e.g., *Caulobacter crescentus*, *Pseudomonas aeruginosa* and *Vibrio cholerae*) are longitudinal irrespective of growth phase or growing condition (Viollier et al., 2004; Vallet-Gely and Boccard, 2013: David et al., 2014).

Longitudinal chromosome configuration in polarized bacteria is maintained by the ParABS system, which tethers one sister *ori* to one pole while pulling the second one to the opposite pole (Surovtsev and Jacobs-Wagner 2018; Jalal and Le, 2020). A variation of longitudinal configuration is the so-called *ori-ter-ter-ori* configuration reported for the diploid Actinomycetales *Corynebacterium glutamicum* (Böhm et al., 2017). In this configuration, the two *ori* are tethered to the poles in non-dividing cells, but in dividing cells two out of four *ori* are segregated to midcell, where they remain until fission is completed. Only then, they get tethered to the newly formed cell poles. As for the obligate symbiont *Candidatus* Thiosymbion oneisti, its *ori* is maintained in the central third of the cell throughout the cell cycle (Weber et al., 2019), albeit its sister *ter* move from midcell to the opposite poles and then back to midcell during segregation (Weber et al., 2019). Taken together, in all bacteria studied so far, irrespective of how the chromosome is configured, the subcellular localization of most genetic loci changes dramatically during chromosome segregation.

Besides differing in their spatial orientation, bacterial chromosomes may differ in their 3D organization (also referred to as chromosome organization or architecture), as revealed by chromosome conformation capture techniques coupled to deep sequencing (e.g., Hi-C, Micro-C) (Marbouty et al., 2015; Wang et al., 2015; Le et al., 2013; Lioy et al., 2018, 2021; Szafran et al., 2021; Ren et al., 2022; Ponndara et al., 2024; Gavrilov et al., 2025). Based on the available genome contact maps, chromosomes are segmented in sub-Megabase chromosomal interaction domains (CIDs) whose boundaries often contain long and highly expressed genes (HEGs) (or gene clusters) that form transcription-induced domains (TIDs; Le et al., 2013; Le and Laub, 2016; Yáñez-Cuna and Koszul, 2023; Gavrilov et al., 2025). Although CIDs are reminiscent of the eukaryotic topologically associated domains (TADs), the mechanisms generating them differ. In mammalian cells, TADs are often delimited by chromosomal loops (Hansen et al., 2018; Szalay et al., 2024). These loops are observed as corner peaks at TAD boundaries and result from the structural maintenance of chromosomes (SMC) cohesin complexes and the CCCTC-binding factor (CTCF) binding the DNA at specific sites (Busslinger et al., 2017; Wutz et al., 2017; Sanborn et al., 2015). In bacteria, chromosomal loops have been described in *B. subtilis.* Initially, they were proposed to be involved in the regulation of *ori* firing (Marbouty et al., 2015; Matthey-Doret et al., 2020) and, later, in the establishment of CIDs during stationary phase via the action of the transcriptional factor Rok (Dugar et al., 2022). Specifically, Rok was shown to form large nucleoprotein complexes that interact with each other over large distances, eventually leading to anchored chromosomal loops. More recently, the nucleoid associated protein (NAP) H-NS was shown to be especially abundant at the base of DNA loops in exponential *E. coli* and to mediate genome-wide long-range DNA looping in deep stationary phase (Way et al., 2025, biorxiv; Gavrilov et al., 2025). Moreover, high gene expression correlated with H-NS-free regions indicating that H-NS represses gene expression in the looped nucleoid (Way et al., 2025, biorxiv). Finally, chromosomal loops have also been detected in euryarchaeota (Cockram et al., 2021) and involved in the aggregation of highly expressed insulated domains in the crenarchaeon *Aeropyrum pernix* (Badel and Bell, 2023). In the former, loops were SMC-dependent but transcription insensitive, whereas in the latter organism, which is SMC-less, loops were transcription sensitive (Cockram et al., 2021; Badel and Bell, 2023).

Here, by applying DNA fluorescence in situ hybridization (FISH) and immunostaining with a specific antibody against the chromosome partitioning protein ParB, we determined the configuration of the chromosomes of three multicellular, longitudinally dividing *Neisseriaceae* that exclusively occur in the oral cavity of mammals. We found that *A. filiformis* is monoploid, whereas *S. muelleri* and *C. steedae* are diploid and that, irrespective of the ploidy their origins of DNA replication are maintained at the host-attached pole throughout the cell cycle. Immunolocalization and in vitro and in vivo binding assays suggest that chromosome longitudinal configuration is likely mediated by the centromeric protein ParB. In agreement with the longitudinal configuration of multicellular oral symbionts, chromosome conformation capture coupled with high-throughput sequencing (3C-seq), revealed significant inter-arm contacts (secondary diagonals) in the chromosomes of exponentially growing *A. filiformis* and *C. steedae*. Moreover, the predicted *parS* sites coincided with discontinuous contact properties (hereafter referred to as chromatin frontiers or frontier) and/or engaged in a chromosomal loop. Besides confirming an involvement of ParBS complexes in chromosome configuration, 3C-seq revealed condition-specific genomic interactions that transcend classical exponential/stationary distinctions. *A. filiformis* and *C. steedae* frontier-forming loci fell into three major classes. Class I comprised operons that matched chromatin frontiers exclusively in exponentially growing liquid cultures (e.g., *atp*, *nuo*, *sdh*, and *pilY1*). These loci encode for growth-coupled envelope systems active during exponential planktonic growth^2^. Class II consisted of loci forming frontiers in both exponential and stationary-phase liquid cultures (e.g., *ppi*, *comEA*, *bam*, and *ubi*). These loci encode factors required for envelope homeostasis, periplasmic folding, DNA handling, and redox maintenance. Class III included loci that formed frontiers only in agar-grown colonies and were strongly enriched for genes involved in peptidoglycan remodeling, adhesins, lipoprotein trafficking, nutrient scavenging, and surface-associated interactions (e.g., *mrdBA*, PBP7, M23, *mlt*, FHA, *lol*, *opp*). Of note, high-flux polar amino acid transporters formed frontiers in both exponentially growing and surface-attached *A. filiformis* and *C. steedae*, acting as cross-conditional loci whose function appears important for both planktonic rapid growth and colony-associated nutrient acquisition.

Altogether, we showed that three obligate oral symbionts maintained longitudinal chromosome configurations irrespective of their ploidy or chromosome segregation. Moreover, chromosome conformation correlated with gene expression dynamics and reflected three distinct physiological states: planktonic exponential growth, planktonic stationary phase, and surface-associated (colony) growth on agar. Each state is characterized by a unique set of frontiers possibly anchored at specific membrane- and envelope-associated loci, revealing that genome folding may convey information about lifestyle and environmental context. These results provide the first 3C-seq contact maps of bacterial colonies and demonstrate that chromosome architecture is a sensitive reporter of bacterial physiology, uncovering a previously unknown structural signature of surface colonization.

## RESULTS

### Chloramphenicol-induced translation blockage suggests that transertion is a major expanding force in multicellular *Neisseriaceae* nucleoids

Nucleoid structure is determined by a balance of compaction forces and one major expansion force (Krogh et al., 2018). The latter is mediated by transertion, a coupling of transcription, translation, and translocation of nascent membrane proteins and/or exported proteins (Cabrera et al., 2009; Bakshi et al., 2014; Roggiani and Gulian, 2015; Spahn et al., 2025).

To determine the localization of the nucleoid and how it reacted to translation blockage, we stained the DNA of exponentially growing multicellular *Neisseriaceae* untreated or treated with chloramphenicol (CHL). In *A. filiformis*, CHL not only led to DNA signal compaction but also to a shift of its localization toward the proximal pole (Fig.1a, Supplementary Fig.1, top panels). As for untreated *S. muelleri* and *C. steedae*, we noticed a decrease of DNA fluorescence at midcell in non-terminal cells (i.e., all the cells belonging to a given filament except the two cells located at its extremities; Fig.1b-c; Supplementary Fig.1, middle and bottom panels) suggesting the presence of two nucleoids. This decrease in DNA signal at midcell was more pronounced in CHL-treated cells suggesting translation blockage-induced compaction of two nucleoids per cell (Fig.1b-c).

In short, based on DNA staining, *A. filiformis* was monoploid and, upon translation blockage, its nucleoid compacted and localized proximally. As for *S. muelleri* and *C. steedae*, they appeared diploid, and the nucleoids of non-terminal cells were more compacted than those of terminal ones. Moreover, the treatment with the translation inhibitor chloramphenicol also resulted in nucleoid condensation in these two oral symbionts suggesting that coupling of transcription and translation at the membrane expands their nucleoids.

### Stable longitudinal chromosome configuration in the monoploid oral cavity symbiont *A. filiformis*

To determine the ploidy and chromosome configuration of *A. filiformis*, we fixed exponentially growing cells and subjected them to DNA fluorescence in situ hybridization (FISH) with probes specifically targeting their predicted *ori* and *ter* (Fig.2, Supplementary Fig.2a, Table S1). We found that 90% of the cells (n=748) had one *ori* and 78% (n=447) had one *ter*, indicating that this bacterium is monoploid (Table S1). Next, we observed that, independently of the cell cycle stage, *ori* localized to the proximal cell third (Fig.2b and Supplementary Fig.2a), whereas *ter* was excluded from it (Fig.2c and Supplementary Fig.2b). Moreover, lateral spreading of *ori* and *ter* foci in replicating *A. filiformis* cells indicated lateral chromosome segregation (Supplementary Fig.2a and b, respectively). Monoploidy and lateral segregation were also confirmed by performing DNA FISH with a probe against a central region of the left arm (Fig.1a, Supplementary Fig.2c, Table S1) and a probe against a central region of the right arm (Fig.1a, Supplementary Fig.2d, Table S1).

**Figure 2.**
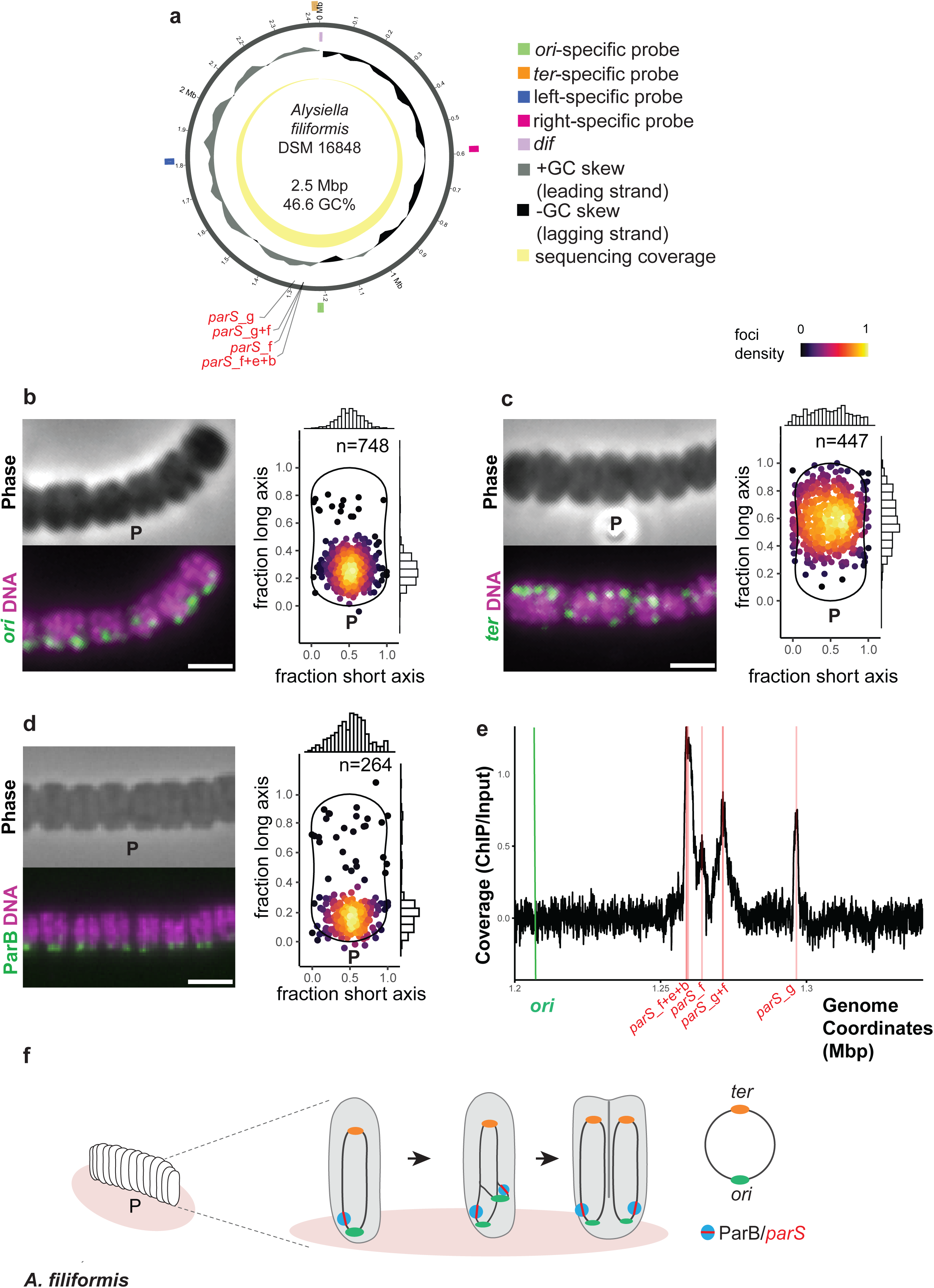
Stable *ori-ter* configuration in *A. filiformis.* (a) Circular map of *A. filiformis* DSM16848 chromosome Outer circle: genomic position (black); middle circle: GC skew (grey for positive, black for negative); inner circle: sequencing coverage (yellow). Colored highlights represent the region targeted by the *ori*-specific probe (green), left arm-specific probe (blue), *ter*-specific probe (orange), right arm-specific probe (pink). *dif* sites are violet. *parS* sites are indicated in red, see Table S3 for their exact genomic positions. (b) Epifluorescence microscope images of representative filaments of *A. filiformis* subjected to *ori* DNA FISH (b) or *ter* DNA FISH (c); phase contrast images are displayed in upper panels and corresponding FISH (green) and Hoechst 3342 (magenta) fluorescence images in bottom panels. Heat maps displaying the subcellular localization of 748 *ori* foci and of 447 *ter* foci in *A. filiformis* flank panel (b) or panel (c), respectively. (d) Epifluorescence microscope image (left) of an *A. filiformis* filament immunostained with anti-ParB antibody (green) and Hoechst 3342 (magenta) and heat map displaying subcellular localization of 234 ParB foci (right). (e) ParB ChIP-seq. The position of the ori is indicated by a green line and peaks of sequence coverage corresponding to the seven predicted *parS* sites are indicated by red lines. (f) Model of *ori-ter* chromosome configuration and lateral chromosome segregation in the monoploid *A. filiformis*. See Table S1 for distribution of *ori, ter,* left, right and ParB foci counts across the population. P: proximal pole. Scale bars are 2 µm.

**Figure 3.**
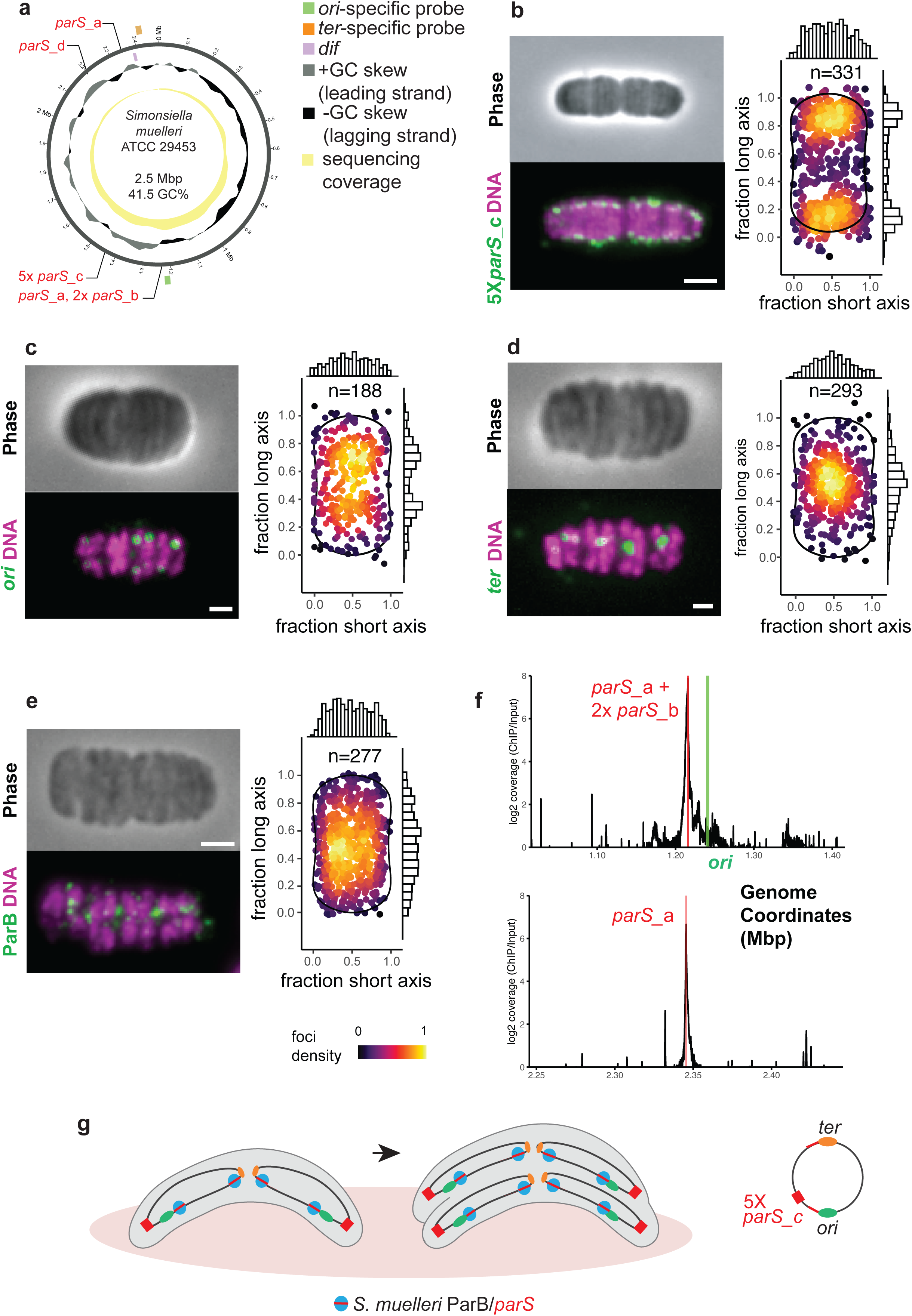
Stable 5X*parS_c*-*ori*-*ter*-*ter*-*ori*-5X*parS_c* chromosome configuration in *S. muelleri*. (a) Circular chromosome maps of *Simonsiella muelleri* DSM 29453. Outer circle: genomic position (black); middle circle: GC skew (grey for positive, black for negative); inner circle: sequencing coverage (yellow). Colored highlights represent the region targeted by the *ori*-specific probe (green), *ter*-specific probe (orange). *dif* site are violet. *parS* sites are indicated in red, see TableS3 for their exact genomic positions. (b-d) Epifluorescence microscope images of representative filaments of *S. muelleri* subjected to (b) 5X*parS*_c DNA FISH, (c *ori* DNA FISH (b) or *ter* DNA FISH (d); phase contrast images are displayed in upper panels and corresponding FISH (green) and Hoechst 3342 (magenta) fluorescence images in bottom panels. Heat maps displaying the subcellular localization of 331 5X*parS*_c foci, 188 *ori* foci and of 293 *ter* foci flank panel (b), panel (c) or panel (d), respectively. (e) Epifluorescence microscope image (left) of an *S. muelleri* filament immunostained with anti-ParB antibody (green) and Hoechst 3342 (magenta), and heat map displaying the subcellular localization of 277 ParB foci (right). (f) ParB ChIP-seq. The position of the *ori* is indicated by a green line and peaks of sequence coverage corresponding to three predicted *parS* sites are indicated by red lines. (g) Model of 5X*parS_c*-*ori*-*ter*-*ter*-*ori*-5X*parS* chromosome configuration and lateral chromosome segregation in the diploid *S. muelleri*. See Table S1 for distribution of *ori, ter* and ParB foci counts across the population. Mbp: Megabase; P: proximal pole. Scale bars are 2 µm.

Further, protein sequence alignment and phylogenetic analysis suggested that *A. filiformis* ParB might be involved in chromosome partitioning (Supplementary Fig.3a-b) and probing protein extracts with a specific anti-ParB antibody showed it was expressed (Supplementary Fig.3c, left). Immunostaining of *A. filiformis* with the anti-ParB antibody revealed that this centromeric protein localized at the proximal pole (Fig.2d). This was consistent with the presence of seven *ori*-adjoining *parS* sites (Fig.2a), chromosome immunoprecipitation (ChIP-seq) assays with anti-ParB antibody (Fig.2e), and with recombinant ParB retarding *parS* sites in electrophoretic mobility shift assays (EMSAs; Supplementary Fig.3d). Based on both in vivo and in vitro data, it appeared that ParB mostly bound one cluster of three *parS* sites (*parS*_f, *parS*_e, *parS*_b), followed by one cluster of two *parS* sites (*parS*_g and *parS*_f), a single *parS*_g site and, finally, a single *parS*_f site.

All in all, *A. filiformis* ParB bound to *ori*-adjoining *parS* sites may mediate *ori* positioning at the proximal pole and longitudinal chromosome configuration. As for the *dif* site-containing *ter* region, it occupies the distal cell half throughout the cell cycle. Finally, loci located either in the center of the left arm or in the center of the right arm occupied the central third of the cell throughout the cell cycle (Fig.2f). Therefore, in *A. filiformis*, not only *ori* but also, *ter,* and the centers of the left or right arms maintain their gross intracellular localization transgenerationally. We will henceforth refer to the chromosome configuration as “stable”.

### Stable longitudinal chromosome configuration in the diploid oral cavity symbiont *S. muelleri*

To determine the ploidy and chromosome configuration of *S. muelleri*, we consecutively applied three DNA FISH probes: (1) a *ori*-specific probe targeting the *ori* and three adjoining *parS* sites (*parS1* dialect a, *parS*2 dialect b and parS3 dialect b; Table S5, Fig.3a, Supplementary Movie 1), (2) a probe targeting a cluster of five, dialect c *parS* sites (*parS_4-8*; so-called 5X*parS*_c site; Fig.3a) situated ca. 250 Kbp away from the *ori* (Table S5), and (3) a *ter-*specific probe targeting a region containing the *dif* sites (Fig.3a). Surprisingly, 5X*parS*_c foci were polar in most of the cells (Fig.3b, Table S1), whereas most *ori* foci were subpolar (Fig.3c, Table S1, Supplementary Movie 1). Moreover, most cells contained two ori foci per cell (57%, n=331; Fig.3b) and most cells contained two 5X*parS*_c foci per cell (43%, n=188; Fig.3b) suggesting that *S. muelleri* is diploid. Of note, marker frequency analysis (MFA) indicated an *ori-ter* ratio between 1 and 2 irrespective of the growth phase, suggesting that the high foci count was not due to multifork replication (Supplementary Fig.5b). As for the *ter* probe, which targeted *S. muelleri dif* sites, most cells had two foci (32%, n=293), which localized to the central third of the cell (Fig.3d, Table S1), consistent with a tail-to-tail arrangement of two longitudinally configured chromosomes (5X*parS*_c-*ori-ter-ter-ori*-5X*parS*_c configuration).

Protein sequence alignment and phylogenetic analysis suggested involvement of *S. muelleri* ParB in chromosome configuration (Supplementary Fig.3a-b) and immunostaining with a specific anti-ParB antibody revealed that this protein was excluded from the poles (Fig.3e). Polar exclusion and ChIP-seq was consistent with ParB not binding the 5X*parS*_c cluster but rather three *ori*-adjoining *parS* sites (*parS1* dialect a, *parS2* dialect b and *parS3* dialect b; Fig.3f top, Table S5) and one *dif*-adjoining *parS* site (*parS10* dialect a; Fig.3f bottom, Table S5). Notably, the a and b dialects of *parS* were also the ones most retarded by recombinant *S. muelleri* ParB in EMSAs (Supplementary Fig.3e).

Collectively, in diploid *S. muelleri*, we observed longitudinal configuration of two chromosomes likely involving the DNA-binding protein ParB tethering *ori*-and *dif*-adjoining *parS* sites to, respectively, subpolar and midcell regions of the inner membrane (Fig.4g). As observed for *A. filiformis*, longitudinal chromosome configuration of both chromosomes was maintained throughout the *S. muelleri* cell cycle.

### Stable longitudinal chromosome configuration in the diploid oral cavity symbiont *C. steedae*

Finally, we sought to determine the chromosome configuration of another crescent-shaped oral symbiont, *C. steedae*. Thus, we applied a DNA FISH probe targeting its *ori* and two adjoining *parS* sites (*parS*_e and *parS*_l; Fig.4a, Supplementary Movie 2) and observed two foci in most cells (43%, n=215; Table S1). Of note, irrespective of whether they were non-replicating or replicating, most *C. steedae* displayed one *ori* focus at one pole and the second one at the opposite pole (Fig. 4b-c, Supplementary Movie 2, Table S1). Based on MFA, the *ori-ter* ratio was between 1 and 2 irrespective of the growth phase suggesting that the high abundance of cells displaying 2 *ori* foci was not due to multifork replication (Supplementary Fig.5c). Not only did the predicted *ori* of *C. steedae* localize at the poles, but due to the larger size of this symbiont, their lateral segregation was clearly discernible when pooling the fluorescence emitted by cells containing 3 *ori* foci (left and right heat map in Fig.4c, respectively).

**Figure 4.**
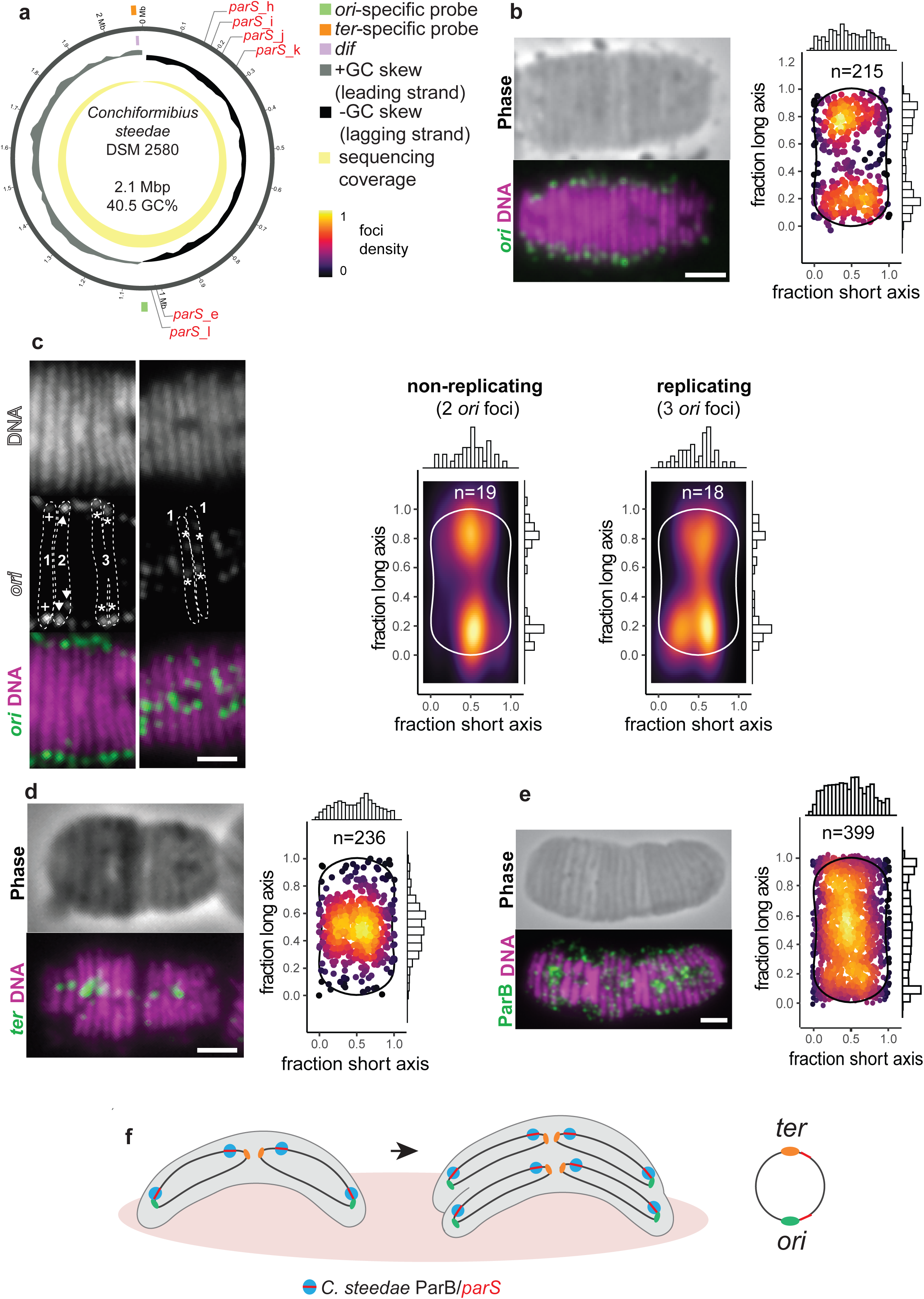
Stable *ori*-*ter*-*ter*-*ori* chromosome configuration in *C. steedae*. (a) Circular chromosome maps of *C. steedae* DSM 2580. Outer circle: genomic position (black); middle circle: GC skew (grey for positive, black for negative); inner circle: sequencing coverage (yellow). Colored highlights represent the region targeted by the *ori*-specific probe (green), *ter*-specific probe (orange). *dif* site are violet. *parS* sites are indicated in red, see TableS3 for their exact genomic positions. (b-d) Epifluorescence microscope images of a representative filament of *C. steedae* subjected to *ori* DNA FISH (b-c) or *ter* DNA FISH (d); phase contrast images are displayed in upper panels and corresponding FISH (green or white) and Hoechst 3342 (magenta) fluorescence images below. In (c) cells with two foci, three foci or four foci were arbitrarily referred to as stage 1 (non-replicating), 2 (early replicating) and 3 (late replicating), respectively. Pluses, arrowheads or asterisks indicate foci in stage 1, 2 or 3 cells, respectively. White dotted cell outlines are cell shapes deduced from phase contrast images (not shown). Heat maps displaying the subcellular localization of 215 *ori* foci, 19 *ori* foci from non-replicating (stage 1) cells, 3 *ori* foci from late replicating (stage 3) cells, or 236 *ter* foci flank representative images in (b-c) or (d), respectively. (e) Epifluorescence microscope image of an *C. steedae* filament immunostained with anti-ParB antibody (green) and Hoechst 3342 (magenta), and heat map displaying subcellular localization of 399 ParB foci (right). (f) Model of *ori*-*ter*-*ter*-*ori* chromosome configuration and lateral chromosome segregation in the diploid *C. steedae.* See Table S1 for distribution of *ori, ter* and ParB foci counts across the cell population. Scale bars are 2 µm.

As for the *ter* probe, which targeted the *dif* sites, we detected at least two foci in 75% of the cells (n=236; Fig.4d, Table S1) and, similar to *S. muelleri*, most *ter* foci localized to the central third of *C. steedae*. As for ParB, it was not only polar, but it appeared throughout the cell (Fig.4e). Given that we predicted both *ori*-adjoining and *dif*-adjoining *parS* sites and given that two terminal *parS* sites (*parS*_h and *parS*_j) matched *C. steedae* chromatin frontiers in all tested conditions (Fig.6 and see below), *C. steedae* chromosome configuration may be mediated by *ori*-proximal and *dif-*proximal regions of the chromatin.

Collectively, the two *C. steedae* chromosomes were juxtaposed resulting in a so-called *ori-ter-ter-ori* longitudinal configuration maintained throughout the cell cycle stage (Fig.4f). Based on its localization pattern, we hypothesize that ParB bound to both proximal and terminal *parS* sites may orient and segregate *C. steedae* chromosomes. Therefore, in all three studied multicellular *Neisseriaceae*, irrespective of their ploidy, *ori* and *ter* occupy the same cellular positions throughout the cell cycle. Of note, DNA FISH and ParB immunostaining also revealed stable chromosome configuration in an obligate symbiont of marine nematodes (Supplementary Fig.4), suggesting that this cellular feature might have convergently evolved in invertebrate and vertebrate symbionts.

### Spatial organization of *A. filiformis* and *C. steedae* chromosomes

We hypothesized that the stable orientation of the chromosomes would enhance population coherence in chromosome conformation capture data and that this, in turn, would facilitate the discovery of physiological state–specific features. To find out, we applied 3C-seq to *A. filiformis* and *C. steedae* and combined it with RNA-seq. Firstly, consistent with the longitudinal configuration revealed by DNA FISH (see above), genome-wide 5 Kbp-bin resolution contact maps revealed low frequency but significant inter-arm DNA contacts (see secondary diagonals in Fig.5, Fig.6 and Supplementary Fig.6). These were more frequent in *C. steedae* than in *A. filiformis* and more frequent in exponentially growing planktonic cells than in stationary or substrate-attached cells (Fig.5, Fig.6 and Supplementary Fig.6).

**Figure 5.**
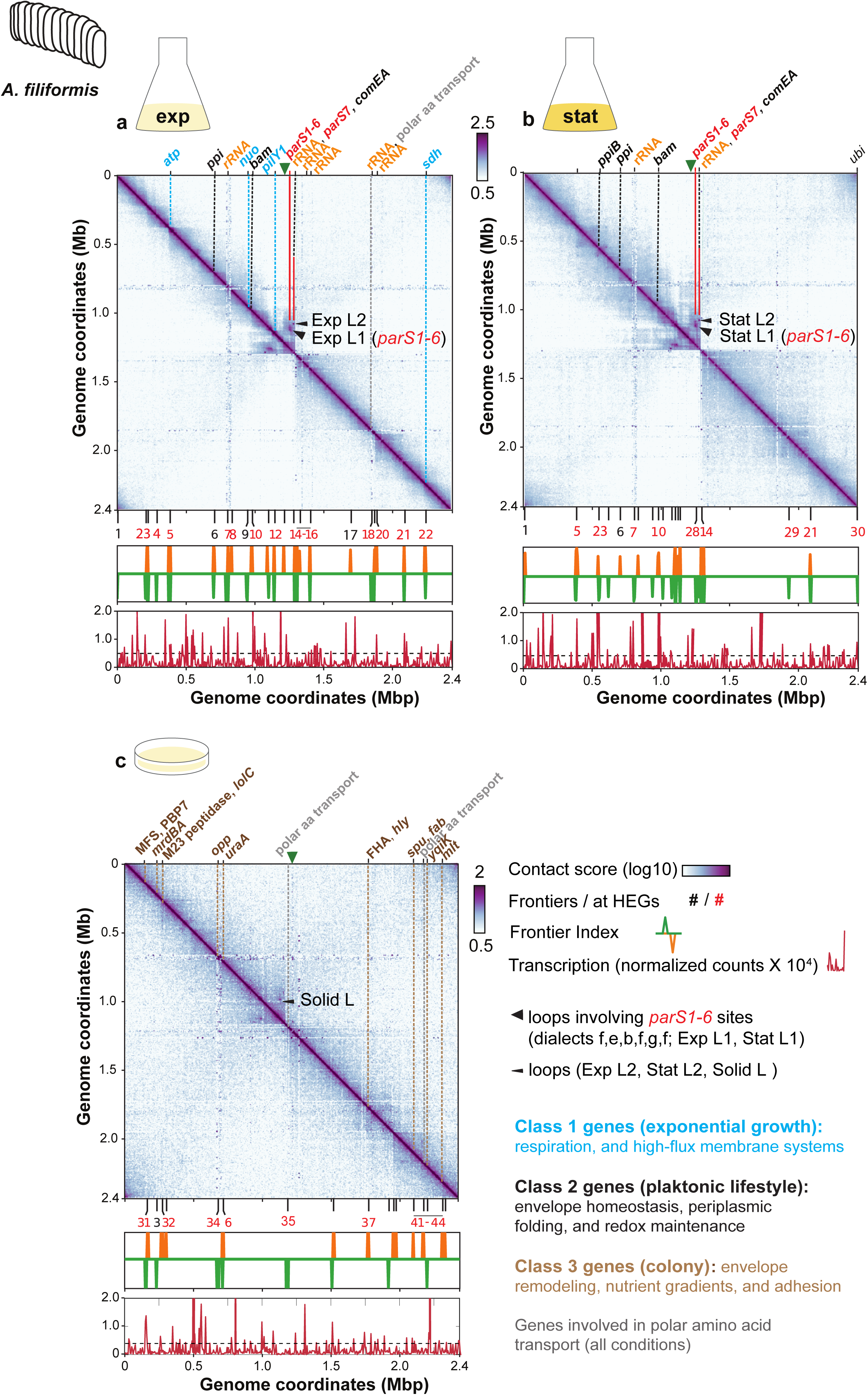
Normalized chromosome contact maps of *A. filiformis.* (a-c) *ori*-centered contact maps of liquid medium exponential (exp) phase cells, stationary (stat) phase cells or cells grown on agar. On top of the map, the green arrowhead indicates the *ori,* genes matching frontier in both *A. filiformis and C. steedae* (same condition) are indicated with their respective abbreviations. In blue are abbreviations of Class 1 genes (growth, respiration, high-flux membrane systems): *atp* (f5), *nuo* (F9), *pilY1* (f12), *sdh* (f22). In black are abbreviations of Class 2 genes (envelope homeostasis, periplasmic folding, redox maintenance): *ppiB* (f23), *ppi* (f6), *bam* (f10), *comEA* (f14) *ubi* (f30). In brown are abbreviations of Class 3 genes (envelope remodeling, nutrient gradients, adhesion): MFS and PBP7 (f31), *mrdBA* (f3), M23 peptidase and *lolC* (f32), *opp* (f34), *uraA* (f6), FHA and *hly* (f37), *spu* and *fab* (f41), *yqiK* (f43), *mlt* (f44). Genes involved in polar amino acid transport (grey) matched f18 in planktonic exponential cells and f35 and f42 in colonies. rRNA operons (orange) matched f7, f14, f15, f16, f18, f20 in planktonic exponential cells and f7 and f14 in planktonic stationary cells. *parS7*_g (red) matched f14 in planktonic cells, and *parS1-4*_*fbef* matched f28 in stationary planktonic cells. See Table S2 for complete list of predicted genes at frontiers and Supplementary Data Set 1,3 for additional information on genes. See Table S3 for genomic positions of rRNA operon and *parS* sites. Black arrowheads indicate chromosomal loops (Exp L1-2, Stat L1-2 and Solid L). See Supplementary Fig.8 and Table S6-7 for more information on loop anchors. Frontier numbers are red if they match HEGs and they are black if they do not. Below each map, the Frontier Index (FI) displays frontiers with upstream (orange) or downstream (green) bias of interaction. Below the FI, the normalized RNA-Seq counts in 5 kb bins show corresponding RNAseq data. The dashed line represents the threshold use to identify highly expressed loci. Contact map in (a) is a merge of 2 replicates. Mb: Megabase. f: chromatin frontier. HEG: highly expressed gene or rRNA operon.

Next, we applied the Frontier Index (FI), a tool specifically developed to analyze domain organization at multiple scales (Lioy et a., 2021; Varoquaux et al., 2022; Kortebi et al., 2026). Specifically, the FI detects bins associated with either a significant upwards or a significant downwards interaction bias at any scale (up- or downstream peaks, respectively, in orange or green in Fig.5, Fig.6, Supplementary Fig.7, Tables S2-S5, Supplementary Data Set 1, Supplementary Data Set 2). We will henceforth refer to loci matching an upstream peak or a downstream peak or both concomitantly (±1 bin) as chromatin frontiers or frontiers. For both *A. filiformis* and *C. steedae*, we detected the highest number of frontiers in planktonic exponential cells (22 and 18 frontiers, respectively; Fig.5a, Fig.6a, Tables S2-S5) than in planktonic stationary, or in colony-forming cells. Moreover, 77% and 100% of the frontiers matched at least one long and/or highly expressed gene (HEG; refers to a gene belonging to the highest expression level, CAT4, or to an rRNA operon; see Supplementary Fig.S8) in exponentially growing *A. filiformis* or *C. steedae*, respectively, suggesting a major role of transcription in shaping bacterial chromatin (Fig.5, Fig.6, Supplementary Fig.7, Tables S2-S5). Of note, none of the frontiers detected in *A. filiformis* colonies persisted when these were treated with RIF or CHL suggesting that not only transcription but also translation shapes the chromatin of multicellular oral symbionts.

**Figure 6.**
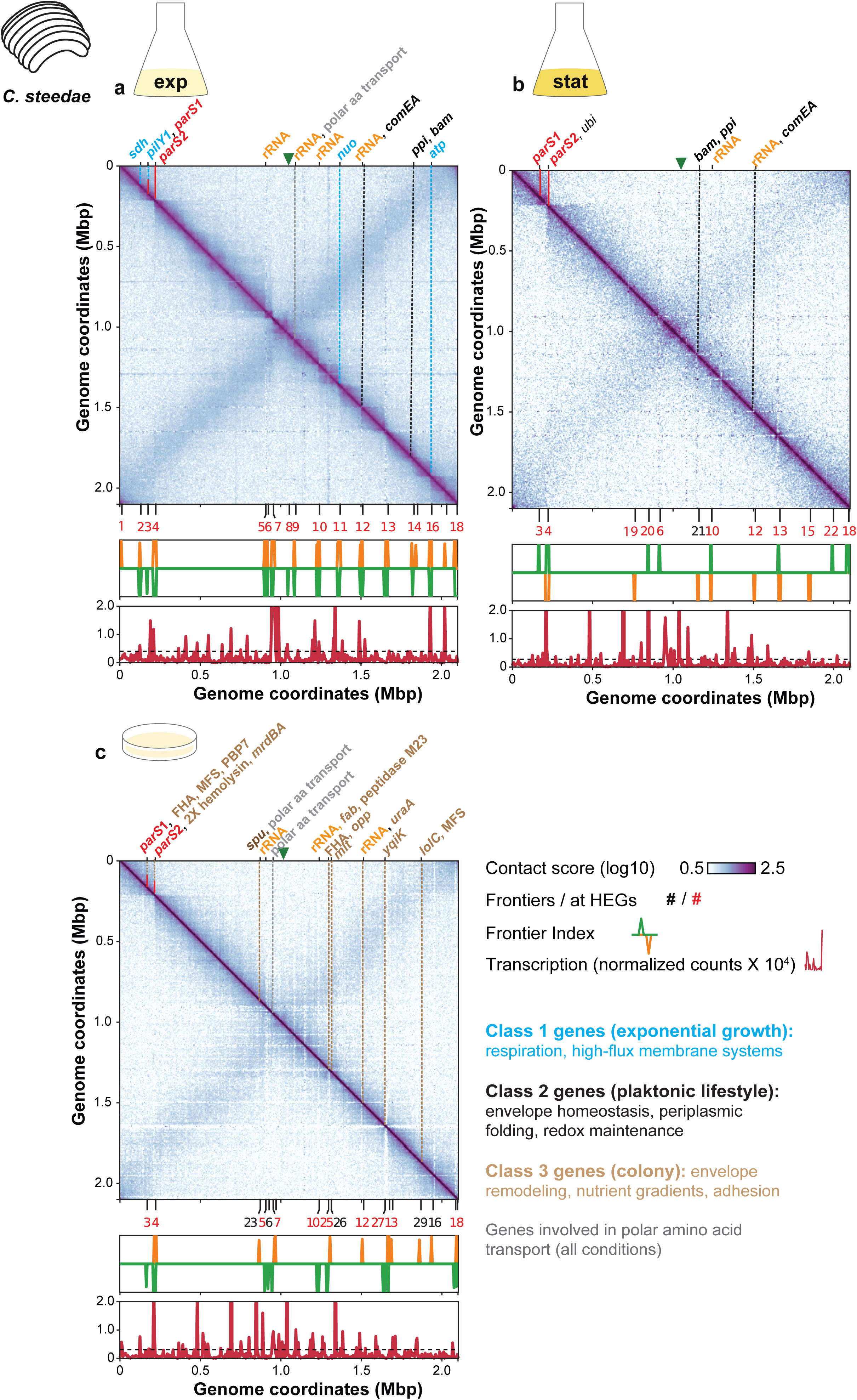
Normalized chromosome contact maps of *C. steedae*. (a-c) *ori*-centered contact maps of liquid medium exponential (exp) phase cells, stationary (stat) phase cells or cells grown on agar. On top of the map, the green arrowhead indicates the *ori,* orthologous genes matching a frontier in both *A. filiformis* and *C. steedae* in the same condition are indicated with their respective abbreviations. In blue are abbreviations of Class 1 genes (growth, respiration, high-flux membrane systems): *sdh* (f2), *pilY1* (f3), *nuo* (F11), *atp* (f16). In black are abbreviations of Class 2 genes (envelope homeostasis, periplasmic folding, redox maintenance): *ppiB* (f23), *bam* and *ppi* (f21), *comEA* (f12), *ubi* (f4). In brown are abbreviations of Class 3 genes (envelope remodeling, nutrient gradients, adhesion): MFS, PBP7 and FHA (f3), *spu* (f23), 2X *hly* and *mrdBA* (f4), M23 peptidase and *fab* (f10), *opp* and FHA (f25), *mlt* (f26). *uraA* (f12), *yqiK* (f27), *lolC* and MFS (f29). Genes involved in polar amino acid transport (grey) are at f7 in exponential phase and at f23 and f7 on solid medium. rRNA operons (orange) are at f5, f9, f10, f12 in exponential phase, at f10 and 12 in stationary phase and at f5, f10, and f12 on solid medium. *parS1*_h (red) matched f3 and *parS2*_j (red) matched f4 in all three conditions. See Table S4 for complete list of predicted genes at frontiers and Supplementary Data Set 2,3 for additional information on genes. See Table S5 for genomic positions of rRNA operon and *parS* sites. Frontier numbers are red if they match HEGs and they are black if they do not. Below each map, the Frontier Index (FI) displays frontiers with upstream (orange) or downstream (green) bias of interaction. Below the FI, the normalized RNA-Seq counts in 5 Kbp bins show corresponding RNAseq data. The dashed line represents the threshold use to identify highly expressed loci. Mbp: Megabase. f: chromatin frontier. HEG: highly expressed gene or rRNA operon.

Strikingly, in both *A. filiformis* and *C. steedae*, the *parS* sites were involved in frontiers and/or loops, linking chromosome configuration with chromosome conformation. In both planktonic and surface-attached *C. steedae*, two terminal *parS* sites, *parS1* (dialect h) and *parS2* (dialect j) matched frontier 3 and 4, respectively (Fig.6, Table S4–S5). In planktonic *A. filiformis*, *parS1-6* (bins 251-254) matched frontier 28 whereas *parS7* matched frontier 14 (Fig.5a-b, Table S2–S3). Moreover, in planktonic *A. filiformis*, *parS1-*6 but not *parS7* engaged in *ori*-containing chromosomal loops (Exp L1 and Stat L1; Fig.5a-b, Table S6, Supplementary Fig.8). As for *A. filiformis* colonies, the only identified loop did not involve any *parS* site, but genes potentially needed for host colonization such as the ones encoding for a cobalt transporter (*cbmI*), a D-amino acid dehydrogenase involved in cell wall remodeling (*dadA*), a transcription repressor involved in efflux-mediated antimicrobial resistance (*tetR*), and a tRNA-dihydrouridine synthase (*dusA*) potentially serving as insertion site for genomic islands or prophages carrying virulence genes (SolidL; Fig.5c, Supplementary Fig.8, Tables S6-S7). Conversely, loop anchors in planktonic cells comprised genes involved in translation, envelope biogenesis, central metabolism, DNA repair and CRISPR-mediated defense (Exp L1-2 and Stat L1-2; Fig.5c, Supplementary Fig.8, Tables S6-S7). Notably, a signal recognition particle (SRP)-containing 15-Kbp adhesin locus was found within *A. filiformis* loop anchors in all tested conditions indicating persistent involvement of this genomic region in high-order chromosome organization (Fig.5c, Supplementary Fig.8, Tables S6-S7).

Collectively, we unveiled the large-scale organization of the chromosomes of two bacteria obligatorily associated with mammals, *A. filiformis* and *C. steedae*. In agreement with what observed in other bacteria (Le and Laub 2016; Lioy et al., 2018, 2021; Ponndara et al., 2024; Abbondanzieri et al., 2025), ≥72% of the frontiers matched at least one HEG irrespective of the growing condition. Moreover, in both symbionts, we observed significant inter-arm chromosomal contacts and the engagement of *parS* sites in chromosomal frontiers. Moreover, in planktonic but not surface-attached *A. filiformis*, most *parS* sites engaged in an *ori-*containing loop further underscoring the link between chromosome configuration and spatial organization. Finally, analysis of chromatin interaction patterns in *A. filiformis* and *C. steedae* across three culture conditions revealed that loci engaged in frontiers fell into three major classes: planktonic exponential phase, planktonic stationary phase, and surface-associated (colony) growth. As presented in the following sections, each state is characterized by a unique set of frontiers containing membrane- and/or envelope-associated transcriptional units.

### Membrane-associated respiratory complexes match frontiers in the chromosomes of exponentially growing *A. filiformis* and *C. steedae*

Three loci (*sdh, nuo, pilY1,*) matched frontiers in exponentially growing *A. filiformis* and *C. steedae* (*A. filiformis* frontiers 22, 9, 12 and *C. steedae* frontiers 1, 11, 3; blue dashed lines in Fig.5a and Fig.6a, respectively; Table S2-5, Supplementary Data Sets 1-2). As for *atp*, it matched frontiers of planktonic *A. filiformis* and exponentially growing *C. steedae* (f5 and f16, respectively). Consistent with rapid growth, *sdh, nuo, atp* encode major respiratory and energy-conserving complexes. More precisely, the *sdh* operon encodes a succinate dehydrogenase (Complex II) which, during rapid metabolism, provides high-flux electron input to the respiratory chain by transferring electrons from succinate to the quinone pool. As for the Complex I NADH:quinone oxidoreductase (Nuo), it oxidizes NADH to NAD⁺ and uses the released energy to pump protons across the inner membrane, thereby providing the primary proton-pumping step in aerobic respiration. Finally, the F₀F₁ ATP synthase Atp converts proton motive force into ATP, and it is therefore the main route for generating bulk ATP during active metabolism and fast biomass production. Of note, *sdh, nuo* and *atp* all encode for large (multi-subunit) inner membrane complexes whose assembly and transertion relies on high translation and membrane insertion flux. As for PilY1, given its role in pilus-mediated adhesion to surfaces and mechanical sensing, it may be required for surface sensing and early adhesion events during active growth. Given that, in *A. filiformis*, *sdh, nuo*, *atp* and *pilY1* were not HEGs (Table S2) it is possible that it is their transertion, rather than their mere transcription, to generate frontiers.

Another hallmark of rapid growth are rRNA gene-matching frontiers. Consistently, six and four rRNA operons matched frontiers in exponential *A. filiformis* (Fig.5a) and *C. steedae* (Fig.6a), respectively, but only two rRNA operons matched frontiers in stationary cells. Finally, gene encoding polar amino acid transporters were detected in exponential planktonic and substrate-attached *A. filiformis* and *C. steedae* (Fig.5a,c, Fig.6a,c, Tables S2,S4). Polar amino acid transporters are high-flux permease and their presence in exponential and agar conditions may reflect their importance for amino acid acquisition during rapid growth and in nutrient-limited colony microenvironments, respectively.

### Genes involved in envelope homeostasis match frontiers in the chromosomes of planktonic *A. filiformis* and *C. steedae* irrespective of their growth phase

Three loci (*ppi, bam, comEA*) matched frontiers in the chromosomes of planktonic *A. filiformis* and *C. steedae*, irrespective of their transcript abundance (black dashed lines in Fig.5a-b and Fig.6a-b, Table S2,4, Supplementary Data Sets 1-2). Their products are envelope-associated factors whose periplasmic or outer-membrane functions may be essential across both exponential and stationary planktonic states. More precisely, peptidyl-prolyl isomerase (PPIase or Ppi) mediates proper folding, quality control, and stress adaptation of periplasmic and envelope-associated proteins, whereas the β-Barrel Assembly Machinery (Bam) catalyzes the insertion and folding of outer membrane β-barrel proteins, a fundamental requirement for maintaining envelope integrity and function. The existence of *ppi* and *bam* frontiers in both planktonic *A. filiformis* and *C. steedae*, irrespective of their growth phase, may reflect persistent envelope assembly demands. As for *comEA*, it encodes a periplasmic, double-stranded DNA-binding competence protein that captures extracellular DNA and delivers it to the inner-membrane translocation machinery (ComEC). Based on its planktonic-specific frontier behavior, ComEA’s role in periplasmic DNA handling and envelope-associated processes seems active in both growing and nutrient-limited cells.

Finally, both stationary *A. filiformis* and *C. steedae* 3C-seq contact maps were characterized by one frontier matching a non-HEG, the ubiquinone (Coenzyme Q) biosynthesis gene *ubi* (frontiers 30 and 4, respectively; Table S2, S4). Beyond its role in respiration, ubiquinone may maintain redox balance within the envelope thereby serving as a major periplasmic antioxidant.

Taken together, planktonic growth was associated with frontiers matching genes encoding periplasmic or outer-membrane proteins whose functions may be essential in both exponential and stationary phase.

### Colony-specific chromatin frontiers and loops reflect surface-associated physiological programs

Colony-specific frontiers were the most abundant type shared by *A. filiformis* and *C. steedae* (Fig. 5c, Fig.6c, Table S2, S4, Supplementary Data Sets 1-2). Loci engaged in frontiers detected exclusively in *A. filiformis* and *C. steedae* colonies clustered into three functional sub-groups: (1) peptidoglycan and envelope remodeling enzymes, (2) nutrient scavenging and transport systems, and (3) surface adhesion and intercellular interaction factors. Genes encoding for PBP7, MrdBA, M23 peptidase, LolC, Fab, YqiK and Mlt, belonged to the first sub-group. All of them may reorganize the cell wall and outer membrane to accommodate colony packing, surface contact, and localized stress. Of note, only three were HEGs, PBP7, the M23 peptidase gene and *mlt*. These cleave PG glycan strands or peptide bridges, facilitating wall expansion, and possibly stress-responsive remodeling at the colony interface.

The sub-group of genes encoding for nutrient-scavenging and import systems comprised one encoding for a major facilitator superfamily permease (MFS; two loci in *C. steedae*), a permease component of the oligopeptide ABC transporter (Opp), a periplasmic binding protein for spermidine/putrescine uptake (Spu), and uracil permease Ura.

Finally, the sub-group of genes encoding for putative host interaction factors comprised filamentous hemagglutinin (FHA) and hemolysin (*hly*) loci, with *C. steedae* frontiers matching two copies of each gene. Filamentous hemagglutinin–like adhesins mediate surface attachment and cell–cell cohesion, whereas hemolysins are toxin-like protein potentially involved in interbacterial or host interaction. Notably, FHA was a HEG only in *A. filiformis* and both *hly* were HEGs only in *C. steedae*, suggesting that transertion, rather than mere transcription, may be prominent in generating frontiers. Accordingly, all agar-specific frontiers disappeared in either transcriptionally or translationally halted *A. filiformis*.

To sum up, agar-specific frontiers in both *A. filiformis* and *C. steedae* matched genes encoding proteins involved in peptidoglycan remodeling, outer-membrane adaptation, nutrient scavenging, and surface adhesion. Although colony growth is heterogenous and likely comprise both exponential and stationary phase cells, colony-specific chromatin signatures differ from those identified in planktonic cells and may be a consequence of the expression of host colonization factors.

## DISCUSSION

During the past decade, studies on symbiont cell biology revealed that obligate bacterial symbionts orient their cytokinetic machineries non-randomly with respect to the animal surface (Leisch et al., 2012, 2016; Pende et al., 2018; Weber et al., 2021, Nyongesa et al., 2022) and they may specifically localize their fimbriae to their host-attached poles (Leisch et al., 2016; Weber et al., 2021; Nyongesa et al., 2022). More recently, we found that they can maintain chromosome orientation toward their host throughout generations (Weber et al., 2019). In this nematode symbiont, the *ori* was maintained at midcell, but *ter* position was so dynamic as to span the whole cell length. Here, we first asked whether obligate symbionts of mammals can stably orient their chromosomes, rather than dynamically flipping them as they replicate and segregate.

Both DNA staining and DNA FISH indicated that the rod-shaped *A. filiformis* is monoploid whereas the crescent-shaped *S. muelleri* and *C. steedae* are diploid, i.e., non-replicating cells have two *ori* and two *ter* and therefore two copies of the same chromosome. Multifork replication explains the dramatic changes of *ori-ter* ratio observed in fast-growing bacteria (e.g., *E. coli*) when they transition from exponential to stationary phase (from 2.5 to 0.97; Ivanova et al., 2015; Reyes-Lamothe & Sherratt, 2019). However, MFA-based *ori-ter* ratios of oral symbionts did not significantly change between exponentially growing (1.9) and stationary phase cells (1.3) suggesting that the relatively high *ori* count observed in *C. steedae* and *S. muelleri* is not merely ascribable to multifork replication. As for the relatively high percentage of *S. muelleri* or *C. steedae* displaying only one *ori* focus instead of two (28% and 27%, respectively) and one *ter* focus instead of two (25%) we ascribe it to foci being closer than the microscope resolution limit. Diploidy has also been reported in pathogenic *Neisseriaceae* (Tobiason and Seifert, 2010), in the Actinomycetes *C. glutamicum* (Böhm et al., 2017) and in the Euryarchaeon *Methanothermobacter thermautotrophicus* (Hildenbrand et al., 2011). Bearing two (or more) chromosome copies increases the chance of correctly repairing the DNA by homologous recombination and may enable cells to withstand genomic instability caused by transposable elements or bacteriophages (Soppa et al., 2014; Zborowsky & Lindell, 2019).

Based on immunostaining and ParB binding assays, we propose that chromosomes are oriented by ParB-mediated membrane anchoring and lateral chromosome segregation. In *A. filiformis* and *Candidatus* T. hypermnestrae, *ori*-adjoining *parS* sites may be involved, consistent with what was found in other polarized rods (i.e., bacteria with longitudinally configured chromosomes and polar asymmetry; Surovtsev and Jacobs-Wagner 2018; Jalal and Le, 2020). In *S. muelleri* and *C. steedae*, in addition to *ori*-adjoining *parS* sites, *dif*-adjoining *parS* sites may be involved in chromosome configuration. This agrees with non-polar ParB signal in immunostaining, the capacity of recombinant ParB to retard both proximal and distal *parS* sites in vitro and *S. muelleri* ParB ChIP-seq data.

Although it is unclear whether stable longitudinal configuration is an adaptation to the symbiotic lifestyle, it was confirmed by the presence of inter-arm contacts in 3C-seq-based contact maps of *A. filiformis* and *C. steedae*. These were most frequent in exponential planktonic cells, decreased in colonies and were the least frequent in stationary planktonic cells. It is unclear, however, why inter-arm contacts were more frequent in *C. steedae* than in *A. filiformis*. In *C. steedae*, this is compatible with a canonical, Smc-ScpAB-based, *ori-ter* oriented loop extrusion mechanism (Wang et al., 2017). How this activity is accommodated in a chromosome with proximal and distal *parS* sites remains to be investigated.

Crucially, the fact that *parS* sites matched frontiers and even engaged in chromosomal loops suggests that ParBS complexes provide architectural scaffolds, linking the chromosome segregation machinery to chromosome conformation. Given that a link between 3D chromosome organization and ParABS-mediates chromosome configuration and segregation has already been observed in archaea (Samson et al., 2025), this phenomenon may represent a conserved principle across prokaryotes. Indeed, in *Sulfolobus*, the SegAB locus, which encodes the segregation machinery, undergoes dynamic folding during DNA replication (Samson et al., 2025).

Prior to this study, intensively transcribed loci were found in at least 60% of CID boundaries (Ponndara et al., 2024, Abbondanzieri et al., 2025). Accordingly, in *A. filiformis* and *C. steedae*, at least 77% of the frontiers matched HEGs with planktonic exponential phase cells having the highest number of frontiers, whereas transcriptionally or translationally halted cells had the lowest. In *A. filiformis*, only frontier 6 was detected in all conditions. It involved loci encoding core envelope-maintenance and respiratory-chain functions (e.g., *ppi, petCBA*), suggesting that certain envelope-associated and redox-stabilizing processes form a condition-invariant architectural scaffold of the chromosome, onto which state-specific frontiers are layered. In *C. steedae*, we found 7 condition- and growth phase-insensitive frontiers. These involved *parS* sites (f3 and f4), rRNA genes (f10 and f12), two genes involved in the biosynthesis and export of exopolysaccharides (f13; Sandkvist et al., 1999; Cuthbertson et al., 2009) and a gene predicted to encode a trimeric autotransporter adhesin (the Yersinia virulence factor YadA; Mühlenkamp et al., 2015; f6). Intriguingly, in *A. filiformis*, a 15Kb-long adhesin gene engaged in chromosomal loops (Exp L1, ExpL2, Stat L1, Solid L) in all tested conditions. Localized transcription and translation of this adhesin, facilitated by stable chromosome configuration, might allow efficient transertion of membrane-bound molecular complexes required for host colonization.

Beyond condition-independent architectural elements, we also identified sets of conserved condition-specific frontiers that matched genes associated with a specific physiological state in both *A. filiformis* and *C. steedae*. Namely, in exponential cells, frontiers coincided with growth-coupled, high-load envelope systems^3^ (Class 1 genes), in stationary cells with envelope and redox maintenance (Class 2 genes), and, finally, during colony growth with peptidoglycan remodeling, adhesion, and nutrient scavenging (Class 3 genes). Thus, by revealing the dominant envelope and metabolic processes active in each lifestyle, chromosome conformation may inform us about the physiological state of a cell.

More specifically, in exponentially growing *A. filiformis* and *C. steedae*, frontiers matched operons encoding major inner membrane bioenergetic complexes (*sdh, nuo, atp*) and the type IV pilus adhesin *pilY1*. These loci specify succinate dehydrogenase (Complex II), NADH dehydrogenase I (Complex I), ATP synthase, and a surface-sensing pilus component. Their identification suggests that active translation, membrane insertion, and assembly of large respiratory and envelope structures impose strong, coordinated organizational constraints specifically during rapid growth. On the other hand, chromatin signatures of stationary *A. filiformis* and *C. steedae* related to envelope maintenance and redox buffering, including *ppi, bam, comEA*, and *ubi*. These encode, respectively, a periplasmic folding catalyst, β-barrel assembly factors, a DNA-binding periplasmic competence protein, and a ubiquinone biosynthetic enzyme. Their persistence across exponential and stationary states indicates a phase-stable requirement for envelope homeostasis, while the stationary-specific prominence of *ubi* highlights the increasing importance of redox control as growth slows. Ubiquinone pools must indeed be replenished or repaired during oxidative and envelope stress, particularly in slow-growing or stressed cells.

Most strikingly, a third regime appeared only in *A. filiformis* and *C. steedae* colonies. Here, frontiers matched genes for cell-wall remodeling, membrane adaptation, nutrient uptake, and adhesion. These included PBP7, M23, *mlt, mrdBA, fabI, lolC, yqiK*, transport systems (MFS, OppB, SpuE, UraA), and major adhesins (FHA, hemolysin). Colony-specific chromatin signatures suggest active restructuring of the envelope and diffusion-limited nutrient acquisition. Although we do not know in which growth phase *A. filiformis* and *C. steedae* colonies were when fixed for 3C-seq, their chromatin features differed from those of exponential or stationary planktonic cells. Agar-specific frontiers may arise because surface-grown colonies impose distinct mechanical, nutritional, and envelope stresses that synchronize peptidoglycan remodeling, adhesion, lipoprotein trafficking, and nutrient scavenging activities across the population. Of note, genes involve in chromosomal loops differed between planktonic and surface-associated cells suggesting that, just like frontiers, chromosome looping is associated with distinct physiological programs. Namely, in planktonic *A. filiformis*, genes in loop anchors mostly belonged to the eggNOG functional categories *Translation, ribosomal structure and biogenesis* (J) and *Cell wall/membrane/envelope biogenesis* (L). Additional recurrent categories included *Amino acid metabolism* (E) and *Replication, recombination and repair* (L). In contrast to the planktonic loops, the colony-specific loop anchors contained a comparatively small set of genes. These were predominantly associated with functions linked to surface-associated physiology, including adhesion, transcriptional regulation, amino acid metabolism, cofactor biosynthesis and cobalt transport, rather than to translation- and envelope-biogenesis-related functions characteristic of planktonic loop anchors. The involvement of a TetR regulator suggests that loop formation in surface-grown cells centers around regulatory control nodes that coordinate adhesion, envelope-associated physiology, and translational remodeling. *dusA* encodes a tRNA dihydrouridine synthase often induced during envelope stress and environmental adaptation (Fruchard et al., 2025). Of note, given that *dusA* may act as a target site for transposition or genomic islands and given that a transposase co-occurred in the same bin, this gene might also have a structural function rather than gene upregulation. Indeed, transcripts of genes in solid medium loop anchors were less abundant than transcripts of genes outside loop anchors (Supplementary Fig.9c).

Transertion has been observed for membrane proteins and was also shown to trigger chromosomal locus repositioning (Libby et al., 2012; Kaval et al., 2023; Kortebi et al., 2026). More recently, single molecule localization microscopy and live-cell imaging indicated that transcription and translation, possibly in the context of transertion, act as a principal organizer of the bacterial nucleoid (Spahn et al., 2025). The nucleoid compaction observed in CHL-treated oral symbionts and the CHL-sensitivity of all the frontiers detected in *A. filiformis* colonies are consistent with transertion being prominent in the multicellular oral symbionts. However, localization pattern analysis of putatively transerted mRNAs is needed to find out whether chromosomes serve as templates for the localization of mRNAs involved, for example, in host colonization.

In conclusion, the application of chromosome conformation capture to bacteria displaying stable chromosome configurations revealed not only growth phase-specific but also environmental state-specific (planktonic versus colony lifestyle) local chromosome conformations. In particular, colony-specific chromosome conformations might reflect transcription and translation of host colonization factors (e.g., pili, hemolysins, adhesins) in naturally occurring multicellular *Neisseriaceae.* Future work should test alternative solid supports to separate surface growth from agar chemistry, and examine mutants or perturbations that disrupt peptidoglycan remodeling, adhesion, or lipoprotein trafficking.

## Supporting information

Supplementary Data Set 1

Supplementary Data Set 2

Supplementary Data Set 3

Supplementary Data Set 4

## AUTHOR CONTRIBUTIONS

T.V. did 3C-seq, marker frequency analysis, RNA sequencing, genome sequencing and assembly, bioinformatic and formal analyses, visualization, revised the manuscript. N.K. did DNA staining, DNA FISH, EMSA, ChIP-seq, immunostaining, formal analyses and visualization, revised the manuscript. P.M.W. did DNA staining, DNA FISH, formal analyses and visualization; E.O.S. contributed to DNA FISH; V.N. contributed to ParB immunostaining and EMSAs; N.P. did ParB protein purification; H.K. contributed to EMSAs; I.J. performed computational analyses of 3C-seq data, developed the loop detection algorithm. N.V. performed preliminary FI analysis. B.D. did DNA FISH. F.B. acquired funding and edited the manuscript. V.S.L. performed 3C-seq analysis, conceptualized and supervised 3C-seq, provided resources, wrote, and revised the manuscript. S.B. conceptualized and supervised the work, acquired funding, provided resources, wrote, and revised the manuscript.

## ACKNOWLEDGEMENTS

This work was supported by the Austrian Science Fund (FWF) project P28593 (T.V. P.M.W, N.K.), FWF project P28743 (T.V., P.M.W), FWF doc.funds MAINTAIN (T.V.), a DOC-fellowship from the Austrian Academy of Science (P.M.W.), a PhD completion grant from the University of Vienna (P.M.W.), by FWF project V 931-B (N.P), by the French National Research Agency: grant number ANR-20-CE35-005 (V.S.L), by the 80 Prime CNRS project MIMIC (F.B. and I.J.) and SIRIG (N.V and V.S.L.) through the MITI interdisciplinary exploratory research program. We thank Bocar Diallo for performing some DNA FISH experiments, Frédéric Veyrier (INRS, Quebec, CA) for introducing *Conchiformibius steedae* to us and for sharing the RNAseq protocol, the members of the V.S.L. and S.B. laboratories for fruitful discussions and S. Bury-Moné for careful reading and feedback on the article. We acknowledge the sequencing and bioinformatics expertise of the I2BC High-throughput sequencing facility, supported by France Génomique (ANR-10-INBS-09). We are also indebted to the staff of the VBCF NGS Unit (Laura-Maria Bayer) for assistance with Oxford Nanopore MinION sequencing. The computational results of this work have been achieved using the Life Science Compute Cluster (LiSC) of the University of Vienna.

## Footnotes

1 Vertically polarized refers to bacterial species whose internal organization is polarized along the long axis of the cell, with distinct functional asymmetry between the two poles.

2 We will henceforth refer to these systems as “high-flux” systems as systems requiring high throughput protein synthesis, envelope assembly or metabolite transport during specific physiological states, e.g., exponential growth (rather than systems involved in respiratory electron flux).

3 i.e., systems that require many proteins being translated and inserted into membranes or exported.

## LEAD CONTACT AND MATERIALS AVAILABILITY

Further information and requests for resources and reagents should be directed to and will be fulfilled by the lead contact, Silvia Bulgheresi. This study did not generate new strains or unique reagents.

## DATA AND CODE AVAILABILITY

Data for genome sequencing has been deposited in Bioproject PRJNA687213 (*A. filiformis*) and PRJNA779076 (*C. steedae*), and genomes have been deposited in GenBank as GCF_020162295.1 (*A. filiformis*) and GCF_023547005.1 (*C. steedae*). Data for marker frequency analysis have been deposited in Bioproject PRJNA795449 (*A. filiformis*) and PRJNA795450 (*C. steedae*), and corresponding non-normalized, 1kbp-binned read count files in Gene Expression Omnibus (GEO) GSE194131 (*A. filiformis*) and GSE205139 (*C. steedae*). Data for RNA-Seq have been deposited in Bioproject PRJNA795348 (*A. filiformis*) and PRJNA795349 (*C. steedae*), and corresponding raw read count files in GEO GSE194132 (*A. filiformis*), and GSE250250 (*C. steedae*). 3C-seq data have been deposited in Bioproject PRJNA837061 (*A. filiformis*) and PRJNA1052629 (*C. steedae),* and corresponding matrices in GEO GSE250400 (*A. filiformis*) and GSE250401 (*C. steedae*). *A. filiformis* ChIP-seq data are available in GEO GSE322725, SRA SRX32354318 and SRX32354319. *S. muelleri* ChIP-seq data are available in GEO GSE322727, SRA SRX32354384 and SRX32354385. SRA accession numbers for sequencing reads and GEO accession numbers for data files are listed in Table S4. The documentation for the ImageJ plugin Fil-Tracer can be accessed here: https://sils.fnwi.uva.nl/bcb/objectj/examples/Fil-Tracer/MD/Fil-Tracer.html. Loop anchor analyses and scripts are accessible at https://github.com/VickyTche/A.-filiformis-loop-anchors-analysis.

## EXPERIMENTAL MODEL AND SUBJECT DETAILS

The bacterial strains *Alysiella filiformis* (DSM 16848), *Simonsiella muelleri* (DSM 2579) and *Conchiformibius steedae* (DSM 2580) have been received from the German Collection of Microorganisms and Cell Cultures GmbH (DSMZ). All *Neisseriaceae* strains were prepared for experimentation as described below unless otherwise noted. *A. filiformis* cultures were pre-cultured by inoculating liquid peptone-yeast (PY) medium with glycerol stocks stored at - 70°C and grown overnight at 37°C at 120 rpm, transferred to fresh PY medium the next day and grown until the specified OD_600_ (see each section). *S. muelleri* and *C. steedae* cultures were streaked out from glycerol stocks onto BSTSY agar plates, grown overnight at 37°C, and colonies transferred into liquid BSTSY medium, and grown at 37°C at 120 rpm until the specified OD_600_ (see each section).

For sampling symbiotic nematodes, sediment samples were collected on multiple field trips (2015-2019) in 1 m depth from a sand bar off Carrie Bow Cay, Belize (16°48’11.01’’N, 88°4’54.42’’W). Specimens of *Robbea hypermnestrae* were extracted from the sediment by stirring the sand in seawater and pouring the supernatant through a 63 µm mesh sieve. The retained material was transferred into a Petri dish, and single nematodes were handpicked using pipettes under a dissecting microscope. For DNA FISH, whole symbiotic nematodes were fixed in 3.7% PFA for 12-14 h at 4°C, washed with 70% ethanol and stored in 70% ethanol at −20°C. Fixed symbiotic nematodes were transported from Carrie Bow Cay to the University of Vienna deep-frozen.

## METHOD DETAILS

### DNA staining of untreated or chloramphenicol-treated symbionts

Methanol-fixed nematodes or oral-cavity symbionts fixed in 3.7% PFA for 12-14 h at 4°C (or for 1 h at room temperature) were rehydrated and washed in PBS containing 0.1% Tween 20 (PBT), followed by incubation in 5 µg/ml Hoechst 3342 PBT for 15 min. After two washing steps to remove unbound DNA stain, worms were sonicated for 45 s to dissociate *Ca*. T. hypermnestrae from its host prior mounting. 1.5 ml of the bacterial suspension was mixed with 0.75 ml of Vectashield mounting medium (Vector Labs) and applied to a 1% agarose covered microscopy slide.

For chloramphenicol treatment, oral cavity symbionts were grown until OD_600_ reached ∼0.5. Chloramphenicol (f.c. 25 μg/mL) was added to the cultures and incubated for 30 min at room temperature shaking at 120 rpm. After incubation the cultures were fixed with formaldehyde (f.c. 3%) for either 12-14 h at 4°C or for 1 h at room temperature. Cells were collected via centrifugation (10,000 x g for 10 min at RT), washed twice with PBS and resuspended in PBS. The DNA was stained as described above.

### Marker frequency analysis (MFA)

Cells were grown until an OD_600_ 0.6 (late exponential phase) and 1 (stationary phase) and collected by centrifugation at 16,000 g for 5 min at room temperature. Cells were lysed using lysozyme and Proteinase K, DNA extracted using Phenol-Chloroform and precipitated using ammonium acetate (2.5M final concentration), two volumes of 100% ice-cold Ethanol, and 5 µg/ml glycogen. RNA was degraded by incubation with 10 mg/ml RNase A for 30 min at 37°C, and DNA cleaned up using DNA Clean & Concentrator-5 (Zymo).

Sequencing libraries were prepared at the VBC NGS facility, and 150 bp paired-end sequenced on an Illumina NovaSeq 6000 machine. Reads were adapter-trimmed and filtered using trimmomatic v0.39 (ILLUMINACLIP:adapters.fa:2:30:10 LEADING:3 TRAILING:3 MINLEN:80 SLIDINGWINDOW:4:15 (Bolger et al., 2014) and prinseq-lite v0.20.4 (-min_qual_mean 30 -min_len 25) (Schmieder and Edwards, 2011). PhiX contaminant sequences were removed using bbduk.sh of bbmap v38.90 (k=31 hdist=1) (Bushnell, 2014), as well as further contaminants (human, animal, plant, fungi) removed against repeat-masked references (minid=0.95, maxindel=3, bwr=0.16, bw=12, quickmatch fast minhits=2, qtrim=rl, trimq=10 untrim). Reads were mapped against the reference chromosomes generated in this study using bowtie2 v2.4.4 (Langmead, B. & Salzberg, 2012), and index-sorted using SAMtools v1.9 (Danecek et al., 2021).

Reads were counted using the bamCoverage program of (standalone) deepTools v3.5.1 (Ramírez et al., 2016) in non-overlapping 1 Kbp windows and the resulting bigwig file converted to human-readable format using the bigWigToWig of UCSC v391. The count data was smoothed using the lowess function (smoother span=0.1) and plotted using ggplot2 (Valero-Mora, 2010) in R v.3.6.1 (R Core Team 2021), and the ratio of the GC-skew predicted *ori* to *ter* coverage (number of non-normalized mapped reads) calculated.

### Genome sequencing and assembly

For whole-genome sequencing, cells were lysed using lysozyme and Proteinase K, DNA extracted using phenol-chloroform and precipitated in two volumes of 100% ice-cold ethanol, 2.5 M ammonium acetate and 5 µg/ml glycogen. RNA was degraded by incubation with 10 mg/ml RNase A for 30 min at 37°C, and DNA cleaned up using DNA Clean & Concentrator-5 (Zymo). The library for Oxford Nanopore Technologies (ONT) sequencing was prepared using the ONT 1D ligation sequencing kit (SQK-LSK110) and sequenced on an R9.4 flow cell (FLO-MIN106) on a MinION for 48 h. Base calling was performed locally with ONT’s Guppy Basecalling Software guppy 5.0.11+2b6dbffa5 (dna_r9.4.1_450bps_sup.cfg), and resulting fastq-files were trimmed using qcat (https://github.com/nanoporetech/qcat). For short read sequencing, sequencing libraries were prepared using the DNA NEB Ultra II FS (NEB) and sequenced on an Illumina NovaSeq SP with 100 bp single-end reads at the VBC NGS facility. The ONT data was assembled using canu v2.1.1 with genomeSize=2.5m (Koren et al., 2017), and polished with the long reads using 5 rounds of racon v1.4.3 (-m 8 -x −6 -g −8 -w 500) (Vaser et al., 2017), reads mapped with minimap2 v2.7 -ax map-ont (Li, 2018) and SAMtools v1.9 (Danecek et al., 2021), and one round of medaka v1.74.0 using the ‘consensus’ (--model r941_min_sup_g507) and ‘stitch’ modules (https://github.com/nanoporetech/medaka). The resulting assemblies were polished with the short reads using 5 rounds of pilon v1.22 (Walker et al., 2014) with bwa v0.7.16a mem (Li and Durbin, 2010) mapped reads. The genome completeness was assessed using CheckM version 1.0.18 (Matsen et al., 2010; Mistry et al., 2013; Parks et al., 2015) with the gammaproteobacterial marker gene set using the taxonomy workflow. Assembled contigs were annotated using Prokka v1.14.6 (Seemann, 2014). Cluster of orthologous genes (COGs) were annotated using eggnog mapper v2.1.10 (Huerta-Cepas et al., 2017) against eggNOG 5.0 (Huerta-Cepas et al., 2019). Additional annotations were obtained through the MicroScope platform (Vallenet et al., 2009).

### Prediction of the origin of DNA replication, terminus of DNA replication and *parS* sites

The *ori* and *ter* positions were identified based on GC skew, sequencing coverage, and *dif* sites prediction (Supplementary Data Set 3). For GC skew analysis we used Genskew (https://genskew.csb.univie.ac.at/) and smoothed using the lowess function (smoother span = 0.005) of gplots v3.1.1 (Warnes, 2016) in R 3.6.1 (R Core Team, 2021). Sequencing coverage was obtained by mapping the raw genomic reads obtained for genome sequencing against the reference genomes using Bowtie2 v2.4.4 (Langmead and Salzberg, 2012) and SAMtools v1.9 (Danecek et al., 2021). Mapped reads were counted and binned in 5 Kbp non-overlapping windows using bamCoverage of (standalone) deepTools v3.5.1 (Ramírez et al., 2016), and the resulting bigwig file converted to human-readable format using the bigWigToWig of UCSC v391. The coverage data was square transformed to better depict differences between *ori* and *ter* and smoothed using the lowess function (smoother span=0.05). *dif* sites were predicted using the betaproteobacterial consensus *dif* sequence (5’-HNNNBNNAYVAYNNDBVTTATGTHAANT-3’) (Carnoy and Roten, 2009). Finally, to identify the *parS* sites, we first computed a consensus *parS* site sequence (5’-DGTTYCAYGTGRAACV-3’) by selecting Gammaproteobacterial *parS* sites out of the 1,030 *parS* sites identified by Livny et al., 2007 and excluding nucleotides that were found 5 times or less at any given position. This led to the identification of all *parS* sites reported in Table S5 but *Sm_parS4-8* (dialect c), *Sm_parS9* (dialect d), *Cs_parS1* dialect h and *Cs_parS2* dialect i. The latter *parS* sites were identified by means of a more degenerate consensus (5’-NGTTNCANGTGNAACN-3’).

In circular plots and 3C-seq heatmaps, the assemblies were reoriented to have the *ori* in the middle of the genome sequence. Circular plots were generated using circos v0.69-8 (Krzywinski, 2009) with Perl 5.028003 (Wall et al., 2000).

### Generation of DNA fluorescence in situ hybridization (FISH) probes

For *Alysiella filiformis*, *Simonsiella muelleri*, and *Conchiformibius steedae* we used Gene-PROBER (Moraru, 2021) to design specific primers hybridizing with multicellular *Neisseriaceae* genetic loci (Table S8). For *Candidatus* T. hypermnestrae probes were designed manually (Weber et al., 2019). For genomic DNA (gDNA) extraction, cells were lysed using lysozyme and Proteinase K, DNA extracted using Phenol-Chloroform and precipitated using ammonium acetate (2.5 M final concentration), two volumes of 100% ice-cold Ethanol, and 5 µg/ml glycogen. RNA was degraded by incubation with 10 mg/ml RNase A for 30 min at 37°C, and DNA cleaned up using DNA Clean & Concentrator-5 (Zymo). For each probe, extracted gDNA was used in 25 µl PCR reactions to generate ca. 9 Kbp-long templates which were in turn used to PCR-amplify ca. 300 nt-long dsDNA polynucleotide probes (see Table S8 and Table S9 for PCR primers and conditions).

All polynucleotide probes were chemically labeled with the Alexa Fluor 594 using the Ulysis Nucleic Acid Labeling Kit (ThermoFisher) following the same modifications to the manufacturers’ protocol as in (Barrero-Canosa et al., 2016).

### DNA fluorescence in situ hybridization (FISH)

Single *Robbea hypermnestrae* nematodes were rehydrated in PBS and nematode symbionts *Ca.* T. hypermnestrae were detached by sonication. Exponentially growing oral cavity symbionts were fixed in 3% or 4% PFA for 12-14 h at 4°C or for 1 h at room temperature, washed with PBS and stored in 1:1 ethanol/PBS. For the hybridization procedure, we followed a slightly modified version of the direct-gene FISH protocol (Barrero-Canosa et al., 2016). After letting the cell suspension dry onto a well of a Poly-L-lysine coated Epoxy-slide, cells were dehydrated in a series of increasing ethanol concentrations and permeabilized with freshly prepared lysozyme solution (0.5 mg/mL) for 1 h on ice or at 37°C. Probes were diluted in hybridization buffer containing 35 or 45% formamide to final concentrations ranging 62-246 pg/ml and each probe was applied to the cells individually. Slides were transferred into a hybridization chamber and incubated for 40 min at 85°C and subsequently at 46°C for 2 h. Washing buffer was applied to the cells once briefly and once for 15 min at 48°C and, finally, cells were incubated in PBS and PBS supplemented with 5 µg/ml DNA stain Hoechst 3342 for 10 min at room temperature. Upon a quick wash in ddH_2_O and, subsequently, in 100% ethanol, cells were air-dried and mounted in 4.5 µl Vectashield mounting medium (Vector Labs) per microscopic slide well.

### ParB protein alignment and phylogeny

ParB alignment was performed with mafft v7.427 (L-INS-I mode) (Katoh and Standley, 2013), and secondary structures predicted using ALi2D (Zimmermann et al., 2018; Gabler et al., 2020) and ESPript 3 (Robert and Gouet, 2014) using the crystal structure of *Caulobacter crescentus* ParB (PDB# 6T1F). The maximum likelihood phylogeny was reconstructed using IQ-TREE v2.1.2 (Minh et al., 2020) with the best-fit model automatically selected by ModelFinder Plus (Kalyaanamoorthy et al., 2013) and 1,000 ultrafast bootstraps (Hoang et al., 2018). The phylogeny was visualized in FigTree v1.4.4 (http://tree.bio.ed.ac.uk/software/figtree/).

### Cloning of *S. muelleri parB* for recombinant protein production in *E. coli*

Genomic DNA of *S. muelleri* and *A. filiformis* was extracted as described for FISH probe generation. The *S. muelleri* genome was used as template for 50 μl PCR reactions to amplify the gene encoding for *S. muelleri* parB (CP083930.1) (Fwd: 5’-CGA ACA GAT TGG TGG CAT GGC AAT CAA ATC AAA AGG-3’, Rev: GTT AGC AGC CGG ATC TTT ATT TAT CAA TGG TTA CAC). The *A. filiformis* genome (CP059564.1) was used as template for 50 μl PCR reactions to amplify the gene encoding for *A. filiformis parB* (Fwd: 5’-CGA ACA GAT TGG TGG CAT GGC GAT TAA AAA AGG CGG-3’, Rev: 5’-GTT AGC AGC CGG ATC TTC AAT TAT CCC ATT TCA CGC-3’). PCR products of the expected size were cloned into the pT7 vector containing an N-terminal 6xHis-SUMO tag by Gibson assembly. Constructs were transformed into chemically competent DH5α *E. coli* cells. Transformation was confirmed by a colony PCR and by Sanger sequencing (Eurofins Genomics, Austria).

### Expression of recombinant *S. muelleri* and *A. filiformis* ParB

Protein expression and purification was carried out as previously described (Pende and Sogues 2021; Studier, 2005). In brief, N-terminal 6xHis-SUMO-tagged ParB from *S. muelleri* and from *A. filiformis* were expressed independently *in E. coli* BL21 (DE3). After 4 h at 37 °C cells were grown for 24 h at 37 °C in 2YT complemented auto-induction medium containing 50 μg/ml kanamycin. Cell pellets were resuspended in 50 ml lysis buffer (50 mM Hepes pH8, 300 mM NaCl, 5% (v/v) glycerol, benzonase, lysozyme, 0.25 mM DTT, EDTA-free protease inhibitor cocktails (ROCHE)) at 4 °C and lysed by sonication. The lysate was centrifuged for 30 min at 30,000x g at 4 °C and loaded onto a Ni-NTA affinity chromatography column (HisTrap FF crude, GE Healthcare). His-tagged proteins were eluted with a linear gradient of buffer B (50 mM Hepes at pH 8, 300 mM NaCl, 5 % (v/v) glycerol, 1M imidazole). The eluted protein of interest was dialyzed at 4 °C overnight in SEC buffer (50 mM Hepes at pH 8, 150 mM NaCl, 5 % (v/v) glycerol) in the presence of the SUMO protease Ulp1 (ratio used, 1:100). The cleaved protein was concentrated and loaded onto a Superdex 75 16/600 size exclusion (SEC) column (GE Healthcare) pre-equilibrated at 4°C in 50 mM Hepes at pH 8, 150 mM NaCl, 5 % (v/v) glycerol. The purified protein was concentrated, aliquoted, flash frozen in liquid nitrogen and stored at −70 °C.

### Western blotting

ParB Western blots were performed as described (Weber et al., 2019). The primary antibody used was either peptide anti-Thiosymbion ParB antibody (Weber et al., 2019) or a guinea pig polyclonal antibody against recombinant *S. muelleri* ParB (1:1000 dilution). The secondary antibody used was a horseradish peroxidase-conjugated anti-rabbit or anti-guinea pig secondary antibody (Amersham Biosciences) with a 1:10,000 dilution.

### Immunostaining

For the oral cavity symbionts, exponentially growing cells were fixed in 3% PFA for 12-14 h at 4°C or 1 h at room temperature, washed with PBS and stored in PBS. Fixed cells were applied on Poly-L-lysine coated cover slips and dried for 15 min at 37°C. The cells were permeabilized with 0.5 mg/ml lysozyme for 12 min at 37°C, briefly washed in water, and then washed in PBS for 1 min. Blocking was performed for 45 min at room temperature in PBS containing 0.1% Tween-20 (PBT) and 2% BSA. Coverslips were incubated overnight at 4 °C with primary antibody (guinea pig polyclonal antibody against *S. muelleri* ParB) diluted 1:1000 in blocking solution. The next day, coverslips were washed in PBT and incubated for 1 h at room temperature with an Alexa555 conjugated anti-guinea pig antibody (Thermo Fisher Scientific) diluted 1:1000 in PBT. After washing the coverslip in PBT and PBS for 1 min and 3 min, respectively, coverslips were washed in water for 1 min, air-dried, and mounted in 3 μl Vectashield mounting medium (Vector Labs) before placement on microscopy slides for imaging.

For the nematode symbionts, deep-frozen methanol-fixed nematodes were rehydrated and washed in PBS containing 0.1% Tween 20 (PBT), followed by permeabilization of the bacterial peptidoglycan by a 15 min incubation with 0.1% (wt/vol) lysozyme at room temperature. Blocking was carried out for 1 h in PBT containing 2% (wt/vol) bovine serum albumin (blocking solution) at room temperature. *Ca*. T. hypermnestrae was incubated with a 1:500 dilution of peptide rabbit polyclonal anti-*Ca*. T. oneisti ParB antibody (Eurogentec) in blocking solution overnight at 4°C. Upon incubation with primary antibody (or without, in the case of the negative control) samples were washed three times in PBT and incubated with a 1:500 dilution of secondary Alexa 488-conjugated anti-rabbit antibody (Jackson ImmunoResearch, USA) in blocking solution for 1 h at room temperature. Unbound secondary antibody was removed by three washing steps in PBT and thereupon incubated in 5 mg/ml Hoechst 3342 PBT for 15 min. After two washing steps to remove unbound DNA stain, worms were sonicated for 45 s to dissociate *Ca*. T. hypermnestrae from its host prior mounting. 1.5 µl of the bacterial suspension was mixed with 0.75 µl of Vectashield mounting medium (Vector Labs) and applied to a 1% agarose covered microscopy slide.

### Fluorescence microscopy

For Fig.1-3 and Fig.4e, cells subjected to DNA staining, DNA FISH or immunostaining were imaged using a Nikon Eclipse NI-U microscope equipped with a MFCool camera (Jenoptik). Images were acquired using the ProgRes Capture Pro 2.8.8 software (Jenoptik). Microscopic images were processed using the public domain program ImageJ (Schneider et al., 2012) in combination with plugin ObjectJ (Vischer et al., 2015) and Fil-Tracer (Nyongesa et al., 2022) or MicrobeJ 5.13J (Ducret et al., 2016). Cell outlines were traced and morphometric measurements recorded. Positions of the fluorescence foci (i.e., points of maximal fluorescent emissions) within the cell boundaries were measured and plotted as fraction of the normalized cell width and length of the cell that contained them. Automatic cell recognition was double-checked manually. For representative images, the background subtraction function of ImageJ was used, and brightness and contrast were adjusted for better visibility. Data analysis was performed using Excel 2026 (Microsoft Corporation), density plots were created with the ggplot2 (Valero-Mora, 2010) in R v.3.6.1 (R Core Team, 2021) and figures were compiled using Illustrator 2026 (Adobe Systems).

### Confocal microscopy

For Fig.4b-d and Supplementary Fig.1, bacteria were visualized with a Leica TCS SP8 X confocal laser scanning microscope. Images were taken with a 63X Plan-Apochromat glycerin objective with a NA of 1.30 and a refraction index of 1.46 (glass slide, glycerin and antifade mounting medium). The Leica software LASX (3.7.2.22383) including the Lightning deconvolution software package (Leica) was used for image acquisition and post-processing.

### 3D Structured Illumination Microscopy (SIM)

SIM was performed on a Zeiss Elyra 7 Lattice SIM microscope (Carl Zeiss, Germany) using Plan-Apochromat 63x/1.4 Oil DIC M27 (WD 0.19 mm) objective with 1.518 immersion oil (Carl Zeiss, Germany). The samples were excited with a 488 nm laser and a 555 nm laser, and the emission was detected using a laser blocking filter LBF 405/488/561/642 and a beam splitter filter SBS BP490-560/LP640. The fluorescence signal was detected on a pco.edge4.2 CL HS sCMOS camera. Raw images are composed of fifteen images per plane per channel (15 phases) and acquired with a Z-distance of 0.11 μm. Acquisition parameters were adapted individually for each recording to optimize the signal to noise ratio. SIM images were processed with ZEN Black 3.0 SR software (Carl Zeiss, Germany) using SIM2. For further image analysis of SIM image z stacks, we used ImageJ Version 2.0.0-rc-68/1.52i. Namely, we assigned a color to the fluorescent channel, stacks were fused to a single image (z projection, maximum intensity), movies were created via 3D projection and images at different rotation angles were extracted. Regions of interest were cut out and, for uniformity, placed on a black rectangular background. Figures were compiled using Adobe Illustrator 2024 (Adobe Systems Inc. USA).

### Electrophoretic Mobility Shift Assay (EMSA)

Forward and reverse primers corresponding to the 16 bp-long *parS* sequences were ordered, along with one primer pair in which the 11 most conserved bases were mutated to serve as a negative control. The primer pairs were hybridized to each other using a 1:1 molar ratio of hybridization buffer (100 mM Tris, 500 mM NaCl, 10 mM EDTA) to primer pair that was heated to 70°C for 5 min then gradually cooled to room temperature overnight. The *parS* sequences were ligated into the pJET 1.2/blunt cloning vector for *E. coli* and then transformed into DH5α *E. coli*. The plasmids were isolated from *E. coli* grown in LB with 100 μg/ml ampicillin using the Monarch Plasmid Miniprep kit. Using specific primers (Fwd: 5’-ACT ACT CGA TGA GTT TTC GG-3’, Rev: 5’-TGA GGT GGT TAG CAT AGT TC-3’) a 338 bp long DNA sequence was amplified by PCR to use as the DNA in EMSAs. EMSAs were carried out as described (Weber et al., 2019) with minor modifications. Briefly, 750 nM, 500 nM, 250 nM, 125 nM affinity purified recombinant ParB was incubated with 200 ng DNA containing either wild-type or the mutated *parS* site in 20 μl total volume, at 30°C for 30 min. As an additional negative control, 750 nM ParB was heat-inactivated by incubating at 99°C for 30 min with 1 mM DTT before being mixed with the DNA fragment. 6x loading dye (10 mM Tris-HCl, 1.5 mg/ml Orange G, 60% (v/v) glycerol) was added to each sample (f.c. 1x) and 11 μl of each sample was loaded onto a 1% agarose gel. Electrophoresis was performed for 1 h at 100 V, subsequently the gel was stained in a SYBR safe gel bath for 20 min and visualized.

### Chromatin immunoprecipitation (ChIP-Seq)

ChIP-Seq was performed as described (Lioy et al., 2018), with minor modifications. Briefly, cells were fixed in exponential phase with formaldehyde (f.c. 1%), incubated for 30 min, after which glycine (f.c. 250 mM) was added. The cells were washed twice in 1X TBS (50 mM Tris-HCl pH 7.4, 150 mM NaCl) and the pellet was resuspended in 0.5 ml lysis buffer (50 mM Tris pH 7.4, 150 mM NaCl, 1 mM EDTA and cOmplete Mini Protease Inhibitor). 10 mg/ml lysozyme was added (f.c. 0.625 mg/ml) and incubated for 20 min at room temperature. Triton X-100 was added (0.5%) and incubated for 10 min. The volume was adjusted to 1 ml with lysis buffer and sonicated using the Covaris S220 Focused-Ultrasonicator for 10 min, intensity 4, peak incident 140W, duty cycle 5%, and cycle per burst 200. The lysate was centrifuged for 30 min at 4°C at 18,000 x g. A 200 μl aliquot was taken as the whole-cell extract and the rest was added to Dynabeads™ Protein A for Immunoprecipitation (Thermo Fischer) washed twice with lysis buffer before use. 10 μg of ParB antibody was added to the beads and incubated for 2.5 h at 4°C under rotation. Subsequently, the beads were washed once in lysis buffer, after which the cell extract was added and incubator for 2 h at 4°C with rotation. The beads were washed three times and resuspended in TE buffer (10mM Tris-HCl pH 7.4, 1mM EDTA). RNA was degraded by incubation with 10 mg/ml RNAse A (0.1 mg/ml) for 30 min at 37°C. Cells were lysed using proteinase K (0.5 mg/ml), SDS (1%) and EDTA (0.025M) and incubated at 65°C overnight. DNA was extracted from both the ChIP samples and whole cell extracts using phenol-chloroform and precipitated in two volumes of 100% ice cold ethanol, 2.5 M ammonium acetate. DNA was cleaned up using a miniElute kit (Qiagen). Raw sequencing reads were quality-processed by adapter and quality trimming using bbmap v38.90 (Bushnell, 2014). The cleaned reads were then aligned to the respective generated genome reference assemblies using Bowtie2 v2.4.4 (Langmead, B. & Salzberg, 2012) with default parameters. For comparative analysis between paired samples, BAM files were processed using bamCompare from deepTools v3.5.1 (Ramírez et al., 2016), with log2 ratio transformation, BPM normalization, bin size 10 bp, pseudocount of 1, and without scaling factor correction. Output was generated in bedGraph format. Downstream visualization and plotting of processed data were performed in R v.3.6.1 (R Core Team 2021).

### Chromosome conformation capture (3C-seq)

3C-seq libraries were prepared as previously described (Marbouty et al., 2015). For growth on solid medium, *A. filiformis* and *C. steedae* were plated from glycerol stock onto BSTSY agar plates and grown at 37°C for 24 h and 48 h, after which they were scraped off and transferred to BSTSY/FBS containing 6% formaldehyde, fully resuspended, and fixed at room temperature at 120 rpm, followed by 30 min more at 4°C. Glycine (250 mM f.c.) was added, and the cells incubated 30 min at 4°C at 120 rpm. After fixation, cells were collected by centrifugation for 10 min at 3,500 g at 4°C, resuspended in fresh medium, collected again by centrifugation, and resuspended in 1 ml fresh medium before final collection by centrifugation. Cell pellets were stored at −70°C. For liquid cultures, *A. filiformis* was grown until OD_600_ of 0.3 (exponential phase) or 1 (stationary phase). For blocking transcription or translation, cells grown overnight on plate were incubated with rifampicin (25 µg/ml) or chloramphenicol (25 µg/ml) for 30 min prior to cell fixation and collection.

For digestion, cell pellets were thawed on ice for 10 min, resuspended in 500 µl Tris 10 mM EDTA 0.5 mM (TE) (pH 8), and lysed using 4 µl Ready-lyse lysozyme (Epicentre) for 20 min at 37°C, 300 rpm. SDS (0.5% f.c.) was added and incubated for 10 min at room temperature. 500 µl lysed cells were then transferred to a tube containing 4.5 ml digestion buffer (1% Triton X-100, 1x CutSmart buffer (New England Biolabs)), and 100 µl cells (non-digested control) were transferred to a tube containing 0.9 ml of digestion buffer. 100U of HpaII (New England Biolabs) were added to the 5 ml digestion mix. Both tubes were incubated for 3 h at 37°C. After digestion, 1 ml of digestion mix was put on ice (digested control) together with the non-digested control. The remaining digestion mix was split into 4 parts, centrifuged for 20 min at 20,000 g, and pellets were carefully resuspended in 1 ml water each, and pooled into 7 ml ligation buffer (1X ligation buffer 3 (New England Biolabs; without ATP), 1 mM ATP, 1 mg/ml BSA, 500 U of T4 DNA ligase 5 U/µl (ThermoFisher)). Ligation was carried out at 16°C for 4 h, followed by incubation together with the two control tubes overnight at 65°C with proteinase K (0.25 mg/ml f.c.) and EDTA (6 mM f.c.). The next day, DNA of the ligated sample was precipitated with 1/10 volume of 3 M sodium acetate (pH 5.2) and two volumes of isopropanol, and after 1 h at −80°C, DNA was pelleted by centrifugation for 30 min at 10,000 g at 4°C and resuspended in 500 µl 1X TE buffer. DNA in all tubes (ligation and two controls) were extracted once with 400 µl phenol-chloroform pH 8.0 (vortexed for 1 min and centrifuged for 5 min 16,000 g at 4°C), precipitated (see above; 30 min at −80°C, followed by centrifugation for 20 min at 16,000 x g at 4°C), washed with 400 µl 80% cold ethanol, and diluted in 30 µl 1X TE buffer supplemented with RNase A (1 mg/ml f.c.; ThermoFisher). Tubes containing ligated DNA were pooled. A 1% agarose gel was run with aliquots of the ligated and two control DNA to check for efficiency of digest and ligation.

Ligated DNA (5 µg) was adjusted with water to 130 µl, and sheared using a Covaris S220 (Duty cycle 5, Intensity 5, cycles/burst 200, time 60 s for 4 cycles), and the DNA purified using the QIAquick PCR purification kit (40 µl elution buffer; Qiagen), and DNA end-repaired with 1X T4 DNA ligase buffer (New England Biolabs), 0.4 mM dNTP mix, 15 U T4 polynucleotide kinase (New England Biolabs), 5 U Klenow DNA polymerase (Roche), and incubated for 30 min at room temperature. DNA was purified using MinElute columns (Qiagen). A-tailing was done with 1X NEBuffer 2 (New England Biolabs), 0.2 mM dATP, and 15 U Klenow exo (New England Biolabs), for 30 min at 37°C before inactivation for 20 min at 65°C, and DNA again purified using MinElute columns (Qiagen). Sequencing adapters were ligated with 0.5 mM custom-made adapters, 2.10^6^ U T4 DNA ligase, and 1X T4 DNA ligase buffer, for 2 h at room temperature, followed by inactivation for 20 min at 65°C. DNA fragments were purified in a size from 400 to 900 bp using a PippinPrep apparatus (SAGE Science). Each library was amplified in 12 cycles with 1X Phusion Buffer (ThermoFisher), 0.2 mM dNTPs, 0.2 µM Illumina primers PE1.0 (5’-AATGATACGGCGACCACCGAGATCTACACTCTTTCCCTACACGA-3’) and PE2.0 (5’-CAAGCAGAAGACGGCATACGAGAT-3’), 3 µl of 3C library, and 1U of Taq Phusion polymerase (ThermoFisher; 98 °C 30 s, 12 cycles of 65°C 30 s, and 72°C 30 s, and 72°C for 7 min). The PCR reaction was purified using MinElute PCR purification kit (Qiagen), and libraries sequenced on an Illumina NextSeq500 with 75 bp paired-end reads.

### Generation of 3C-seq contact maps

Raw contact maps were built as described previously (Lioy et al., 2018). Briefly, each read was assigned to a restriction fragment. Non-informative events such as self-circularized restriction fragments, or uncut co-linear restriction fragments were discarded. The chromosome of each bacterium was divided into 5 Kbp bins and the frequencies of contacts between genomic loci for each bin were assigned. Contact frequencies were visualized as heatmaps. Raw contact maps used for comparative analyses were built using 1 million valid reads. Raw contact maps were normalized using Iterative correction and eigenvector decomposition (ICE) (Imakaev et al., 2012). To facilitate visualization, normalized contact matrices are visualized as log matrices. A Gaussian filter of H=0.5 is used when plotting the matrices. To quantify the frequency of interactions in the secondary diagonals (Supplementary Figure 6), we applied a method to our data based on the filtering of significant 3C contacts and developed as described (Wang et al., 2017).

### Identification of frontiers in contact maps

To identify frontiers, we applied the Frontier Index (FI) method as described (Varoquaux et al., 2022; Lioy et al., 2021; Kortebi et al., 2026). Briefly, the FI method allows to identify loci associated with demarcations, or frontiers, in the 3C contact maps that can occur at any scale, meaning that the method is useful to identify domain organization irrespective of the underlying scale. Specifically, at a given locus (let say *i*), two frontiers can be found at most, each one corresponding to a side of the locus – one frontier indicates that the neighboring loci on the same side of the locus *i* tend to make more contacts with each other with respect to neighboring loci separated by the same genomic distances but located on each side of the locus Identified frontiers appeared as upstream or downstream peaks in Fig.5-6 and SFig.7.

### Identification of *A. filiformis* chromatin loops

Significantly interacting loci within the loop anchors were identified using a Viewpoint-based 4C/Hi-C approach. A viewpoint is a genomic position chosen as an anchor from which interaction frequencies with the rest of the chromosome are extracted, generating a one-dimensional contact profile analogous to a virtual 4C experiment. Viewpoint 2 (bins 251–254) contains *parS1–6*, whereas Viewpoint 1 (bins 227–229) and Viewpoint 3 (bins 257–258) were identified by a loop detection algorithm as follows. Firstly, to facilitate analysis, normalized bins interacting with less than 10% of the total number of bins were considered empty, reflecting multi-mapping reads originating from repetitive regions such as multiple rRNA operons, and were removed from further analysis. In practice, the corresponding rows and columns were removed from the normalized 3C-seq contact matrices, and all subsequent analyses were performed on the resulting reduced matrices. Chromatin loop anchors were identified using a multi-step approach. First, the filtered 3C-seq matrices were slightly smoothed using a Gaussian filter (width = 0.5). A virtual 4C interaction profile was then computed for each genomic bin by treating each bin as a viewpoint (anchor). Peaks in these profiles were identified as candidate chromatin loops, with peak height defined as the difference between the interaction frequency at the peak and the maximum of the two flanking minima. The distribution of peak heights consisted of an exponential distribution for low values and a clear deviation for higher values. A conservative threshold separating these two distributions was used to identify significant peaks, which were defined as chromatin loops. Loop anchors were subsequently defined as maximal sets of contiguous bins in which any two consecutive bins were involved in at least one significant loop.

All bins contained in each of the three viewpoints were analyzed independently. Namely, for each Viewpoint, interaction frequencies were compared to a distance-dependent threshold, defined as the 95th percentile of Hi-C contact frequencies calculated within genomic distance intervals (1–5, 5–10, 10–20, 20–40, 40–80, 80–160, 160–320, and >320 bins). Genomic bins with contact frequencies exceeding the corresponding distance-dependent threshold in all viewpoint bins were considered as putative interacting loci. The viewpoint region and its adjacent bins (±1 bin) were excluded from the analysis to avoid calling local contacts as positive interactions. When two or more interacting regions were identified for a single viewpoint, they were merged if they belonged to the same anchor region (i.e., they belonged to the same 4C peak).

### Transcriptomics

All organisms were grown in the same conditions as those applied for the 3C-seq experiments in three biological replicates each as for the 3C-seq experiments. 500 µl liquid culture were mixed well with 1 ml RNAprotect Bacteria Reagent (Qiagen) and incubated for 5 min, and cells grown on solid medium scraped off and resuspended in 1 ml thereof. Cells were collected by centrifugation for 10 min at 5,000 g, and cell pellets stored at −70°C until the next day.

Total RNA was extracted using the RNeasy Mini Kit (Qiagen) for low cell numbers, including the on-column DNA digestion. Eluted RNA (30 µl) was treated with 2 U of DNase for 30 min using the TURBO DNA-free kit (Invitrogen). RNA was cleaned up using the RNA Clean & Concentrator-25 kit (Zymo) including the on-column DNA digestion. Absence of DNA was checked via PCR (35 cycles) against a ∼200 bp fragment of *gyrB*. RNA was quality checked at the VBC NGS facility (all samples had a RIN>9), rRNA depleted using an in-facility developed protocol, and reverse-stranded sequencing libraries were prepared using the NEBNext Ultra II Directional RNA Library Prep Kit (New England Biolabs). Libraries were sequenced on an Illumina NovaSeq 6000 machine, using 100 bp single-end reads (*A. filiformis*) and 150 bp paired-end reads (*C. steedae*). Reads were adapter-trimmed and filtered using trimmomatic v0.39 (ILLUMINACLIP:adapters.fa:2:30:10 LEADING:3 TRAILING:3 MINLEN:80 SLIDINGWINDOW:4:15; (Bolger et al., 2014)) and prinseq-lite v0.20.4 (-min_qual_mean 30 -min_len 25) (Bushnell, 2014), as well as further contaminants (human, animal, plant, fungi) removed against repeat-masked references (minid=0.95 maxindel=3, bwr=0.16, bw=12, quickmatch fast minhits=2, qtrim=rl, trimq=10 untrim). Reads mapping to rRNA were removed using SortMeRNA v4.1.0 (Kopylova et al., 2012) against the pre-built SortMeRNA databases and the organism-specific rRNA sequences. Reads were mapped against the reference genome generated in this study using Bowtie2 v2.4.4 (Langmead and Salzberg, 2012), Q>30 filtered and index-sorted using SAMtools v1.9 (Danecek et al., 2021). Reads were counted using featureCounts of the subread-package v2.0.0 (Liao et al., 2014) in single-end mode (-s 2) or in paired-end mode (-s 2 -p -B -C) against the “gene” features of the reference genomes’ gff-files.

To show transcription levels along the chromosome, trimmed reads were mapped against the reference chromosomes using Bowtie2 v2.4.4 (Langmead and Salzberg, 2012), and index-sorted using SAMtools v1.9 (Danecek et al., 2021). Per-basepair coverage was determined using SAMtools depth (-a -d 0) (Danecek et al., 2021) and binned in 5 Kbp bins using a custom script in R v3.6.1 (R Core Team, 2021). Counts were normalized to the sum of counts per sample and square-root transformed. All plots were made using matplotlib v3.5.1 (Hunter, 2007) in python v3.7.12 (Van Rossum and Drake, 2009).

For differential expression analysis, raw per-gene counts and metadata (i.e., conditions) were imported using DESeqDataSetFromMatrix in DESeq2 package v1.30.1 (love et al., 2014) in R v4.0.3 (R Core Team, 2021), and differential expression analysis carried out using the function ‘DESeq’, and results extracted using the function ‘results’ with padj.cutoff=0.05 and lfc.cutoff=log2(2). For each sample, raw per-gene counts were divided by gene length, and TPM values calculated by multiplying each normalized count value by 10^6^, divided by the sum of all counts per sample. Per-sample TPM values were averaged per condition. Genes were classified into expression level categories by considering the distribution parameters as indicated in Supplementary Fig.9.

## Supplementary Figure legends

**Supplementary Figure 1. DNA localization in *A. filiformis*, *S. muelleri* and *C. steedae* cells.** Schematic representations and confocal microscope images of (a) *A. filiformis,* (b) *S. muelleri* and (c) *C. steedae* labeled with the peptidoglycan precursor TADA for 60 min (false colored in green) and stained with Hoechst 3342 (magenta). (a) Frontal view of an *A. filiformis* filament reveals a V-shaped DNA localization pattern due to longitudinal septation starting at the distal (D) pole (Nyongesa et al., 2022). Rightmost panel displays a 90 degrees rotated view of the cells marked with an asterisk. (b) Viewed from the top*, S. muelleri* DNA is absent from the center of a non-terminal, likely proliferating cell (n) or present at the center of a terminal (t), likely non-replicating cell. Rightmost panel displays side views of a non-terminal (n) and a terminal (t) cell. Rightmost panel displays side views of a non-terminal (n) and a terminal cell (t). (c) Viewed from the top, *C. steedae* DNA is absent from the center of a non-terminal, likely DNA replicating cell (n) or present at the center of a prospective terminal, non-replicating cell (t). Rightmost panel displays side views of a non-terminal (n) and a terminal (t) cell. Scale bars are 1 µm.

**Supplementary Figure 2. Lateral segregation of *A. filiformis* chromosome.** (a) DNA FISH-based subcellular localization of *A. filiformis ori* in 709 non-replicating (1 *ori* focus; left) or 39 replicating (2 *ori* foci; right) cells. (b) DNA FISH-based subcellular localization of *A. filiformis ter* in 402 non-replicating (1 *ter* focus; left) cells or in 45 replicating (2 *ter* foci; right) cells. (c) Epifluorescence microscope image of a representative filament of *A. filiformis* subjected to DNA FISH with a probe targeting the left arm of its chromosome; phase contrast image is displayed in the upper panel and corresponding FISH (green) and Hoechst 3342 (magenta) fluorescence image in the bottom panel. Flanking heat maps display the subcellular localization of left probe in 355 non-replicating (1 focus; left heat map) or 17 replicating (2 foci; right heat map) cells. (d) Epifluorescence microscope image of a representative filament of *A. filiformis* subjected to DNA FISH with a probe targeting the right arm of its chromosome; phase contrast image is displayed in the upper panel and corresponding FISH (green) and Hoechst 3342 (magenta) fluorescence image in the bottom panel. Flanking heat maps display the subcellular localization of the right probe in 420 non-replicating (1 focus; left heat map) or 84 replicating (2 foci; right heat map) cells. P: proximal. Scale bars are 2 µm.

**Supplementary Figure 3. Structural alignment of ParB, phylogenetic placement, protein expression in animal symbionts and ParB/*parS* electrophoretic mobility shift assays (EMSAs).** (a) Secondary structure prediction of selected ParB proteins. Red indicates predicted alpha helices whereas blue indicates predicted beta strands. The experimental secondary structure elements are inferred from the *Caulobacter vibrioides* ParB and depicted above each sequence block. Prediction was performed using PSIPRED and Ali2D. *B. subtilis*: *Bacillus subtilis. C. steedae*: *Conchiformibius steedae*. *A. filiformis*: *Alysiella filiformis. S. muelleri*: *Simonsiella muelleri. N. elongata*: *Neisseria elongata. N. meningitidis*: *Neisseria meningitidis. Ca.* T. oneisti: *Candidatus* Thiosymbion oneisti. *Ca.* T. hypermnestrae: *Candidatus* Thiosymbion hypermnestrae. ParB box I and II, as well as HTH (*parS* binding) sites are indicated below the alignment. (b) Unrooted phylogenetic tree including ParB of nematode and oral cavity symbionts, as well as diverse other bacterial ParB. Sequences on mauve background are predicted to mediate chromosome segregation whereas sequences on rose background were predicted to mediate plasmid segregation based on Ghosh et al., 2006. The analysis is based on a MAFFT alignment of full-length proteins (data not shown) and estimated under the Q.pfam+G4 model using ML analysis (IQ-TREE) with node support calculated by UltraFast bootstraps. Scale bar represents 0.7% estimated sequence divergence. Closed circles represent values ≥95% based on 10,000 replicates. (c) Western blots of protein extracts of, from left to right, *A. filiformis*, *S. muelleri*, *C. steedae* and *Ca.* T. hypermnestrae probed with an anti-ParB antibody or with the secondary antibody only (-). Numbers indicate apparent MWs expressed in kDa. (d) Electrophoretic mobility shift assays (EMSAs) of recombinant *A. filiformis* ParB and four *parS* dialects or a mutated *parS* site (top). (e) Electrophoretic mobility shift assays (EMSAs) of recombinant *S. muelleri* ParB and four *parS* dialects or a mutated *parS* site (top).

**Supplementary Figure 4. DNA localization, chromosome configuration and ParB localization in *Ca.* T. hypermnestrae.** (a) Epifluorescence microscope images of Hoechst 3342 staining of fixed *Ca.* T. hypermnestrae cells. Cells are arranged from the youngest to the oldest from left to right. Upper panels display phase contrast images of cells and lower panels their corresponding DNA fluorescence (white dotted line represent cell outlines). (b) Averaged DNA fluorescence (a.u.) plotted against the cell length (right; n=447). (c and f) Histograms showing number of (c) *ori* foci (n=413) and (f) *ter* foci (n=576) per cell. (d and g) Confocal microscope images of *Ca.* T. hypermnestrae cells subjected to (d) *ori* DNA FISH or (g) *ter* DNA FISH; phase contrast images are displayed in upper panels and corresponding FISH (white) fluorescence images in bottom panels. White dotted lines represent cell outlines. (e) Subcellular localization of 413 *ori*, 320 *ftsQAZ* and 1,124 *parS2* foci (from left to right) and (h) subcellular localization of 576 *ter* foci. (i) Representative images showing five *Ca.* T. hypermnestrae cells arranged from the youngest to the oldest from left to right. Panels display phase contrast images, ParB (green) and DNA fluorescence (Hoechst 3342; red) and corresponding overlays (from top to bottom). White dotted lines represent cell outlines. (j) Subcellular localization of symbiont ParB foci of 576 cells immunostained with an anti-ParB antibody. (k) Averaged ParB and DNA fluorescence (a.u.) plotted against the cell length (n=82). (l) Model of *ori-ter* chromosome configuration and lateral chromosome segregation in the monoploid *Ca*. T. hypermnestrae. Scale bars are 1 µm.

**Supplementary Figure 5. Marker frequency analysis (MFA) of (a) *A. filiformis*, (b) *S. muelleri*, (c) *C. steedae* and (d) *E. coli*.** Cells were collected in exponential (exp) or stationary (stat) phase (left and right plots, respectively). The x axis represents the genomic coordinates of the reference genome, and the y axis the coverage expressed in number of mapped reads. The coverage along the chromosome (black dots) is smoothed (red line), and horizontal lines represent the coverage at the *ori* (highest coverage) and the coverage at the *ter* (lowest coverage) used to calculate the *ori*-*ter* ratio. For comparison, we performed MFA of *Escherichia coli* in (g) exponential phase and (h) stationary phase based on Ivanova et al., 2015.

**Supplementary Figure 6. Contact map analysis of the primary and secondary diagonal in *A. filiformis* and *C. steedae*.** (a-c) *ori*-centered contact maps of *A. filiformis* and (d-f) *C. steedae* were generated after setting the threshold for significant interactions at 0.5X (top) or 0.25X (bottom) of the standard deviation above the median to the normalized contact maps obtained in each condition. Contact frequencies above or below this threshold were assigned a value of 1 or 0, respectively, generating a binary contact map in which significant interactions are plotted in yellow and non-significant interactions in blue (left). Additionally, the signal in the binary map is connected by linking five elements considered significant using a diamond-shaped kernel of size 15 to fill empty points defined by the connected elements (right).

**Supplementary Figure 7. Normalized chromosome contact maps of *A. filiformis* grown on solid medium with rifampicin or chloramphenicol or without.** Contact map of *A. filiformis* grown on solid medium and incubated with (a) 25 µg/ml rifampicin (RIF), (b) 25 µg/ml chloramphenicol (CHL) for 30 min or without (c; identical to panel c of Fig.5). Below each map, the Frontier Index (FI) displays frontiers with upstream (orange) or downstream (green) bias of interaction. See Table S2 for frontier descriptions and Fig.5 for detailed description of panel c.

**Supplementary Figure 8. Hi-C contact maps and interaction frequency profiles across genomic loci.** (a-c) Magnification of *A. filiformis* contact maps (log10 scale) for liquid medium exponential phase (exp), liquid medium stationary phase (stat), and solid medium conditions centered at the origin of replication (*ori,* green). Viewpoints are black except for Viewpoint 2 which contains *parS1-6*. (d-f) Frequency of contacts (log10) across genome coordinates for Viewpoint 1 (bins 227-229; d), Viewpoint 2 (bins 251-254; e) and Viewpoint 3 (bins 257-258; f) in liquid medium exponential phase, liquid medium stationary phase, or on solid medium. Anchors interacting with viewpoints are grey in d or light blue in e-f. A green arrowhead indicates the origin of DNA replication. *parS1-6* are red and *parS7* are red. See Supplementary Data Set 4 for further details.

**Supplementary Figure 9. Transcriptomic data of oral cavity symbionts**. (a) *A. filiformis* or (b) *C. steedae* transcriptomes were binned at 5 kb resolution and represented as boxplots. The boxplots represent the first quartile (Q1), median and third quartile (Q3). The upper whisker extends from the hinge to the largest value no further than 1.5 * the inter-quartile range (IQR, i.e., distance between the first and third quartiles) from the hinge. The lower whisker extends from the hinge to the smallest value at most 1.5 * IQR of the hinge. Outliers were considered for the analyses, but they are not shown for simplicity. (b) Description of categories of expression level of each chromosomal bin, related to panel a. **(c)** Boxplots comparing gene expression levels (log2(TPM)) of *A. filiformis* genes located in loop anchors versus those not located in loop anchors in cells grown in liquid culture or on solid medium. Gene expression (TPM) of genes located in loop anchors was compared to genes not in loop anchors within each condition (planktonic exponential, planktonic stationary, and colony) using a two-sided Wilcoxon rank-sum test with Benjamini–Hochberg correction for multiple testing. In the solid condition, genes within loop anchors were significantly undertranscribed compared with genes outside loop anchors (median log₂(TPM + 1) = 5.48 vs. 6.58; Wilcoxon rank-sum test, W = 49081, Benjamini–Hochberg-adjusted p = 0.017).

**Supplementary Movie 1.** 3DSIM of an *S. muelleri* filament subjected to DNA FISH with an *ori*-specific probe (green) and stained with WGA (false colored in red). Scale bar 2µm

**Supplementary Movie 2.** 3DSIM of a C*. steedae* filament subjected to DNA FISH with an *ori*-specific probe (green) and stained with WGA (false colored in red). Scale bar 2µm.

