## Supplementary figures and images for "Conserved lifestyle-associated chromosome architectures across mammalian symbionts"

### Supplementary Fig.1

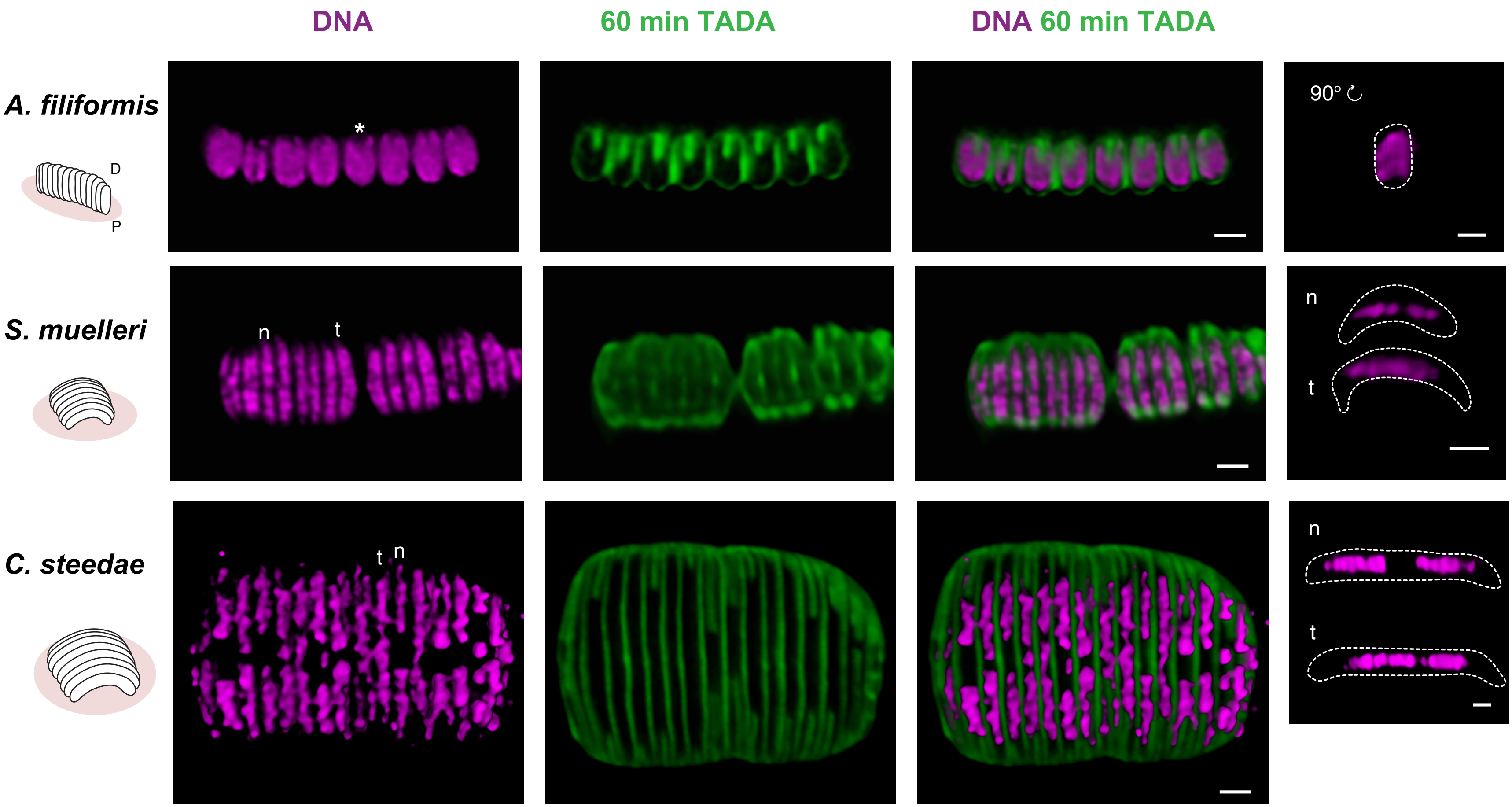

### Supplementary Fig.2

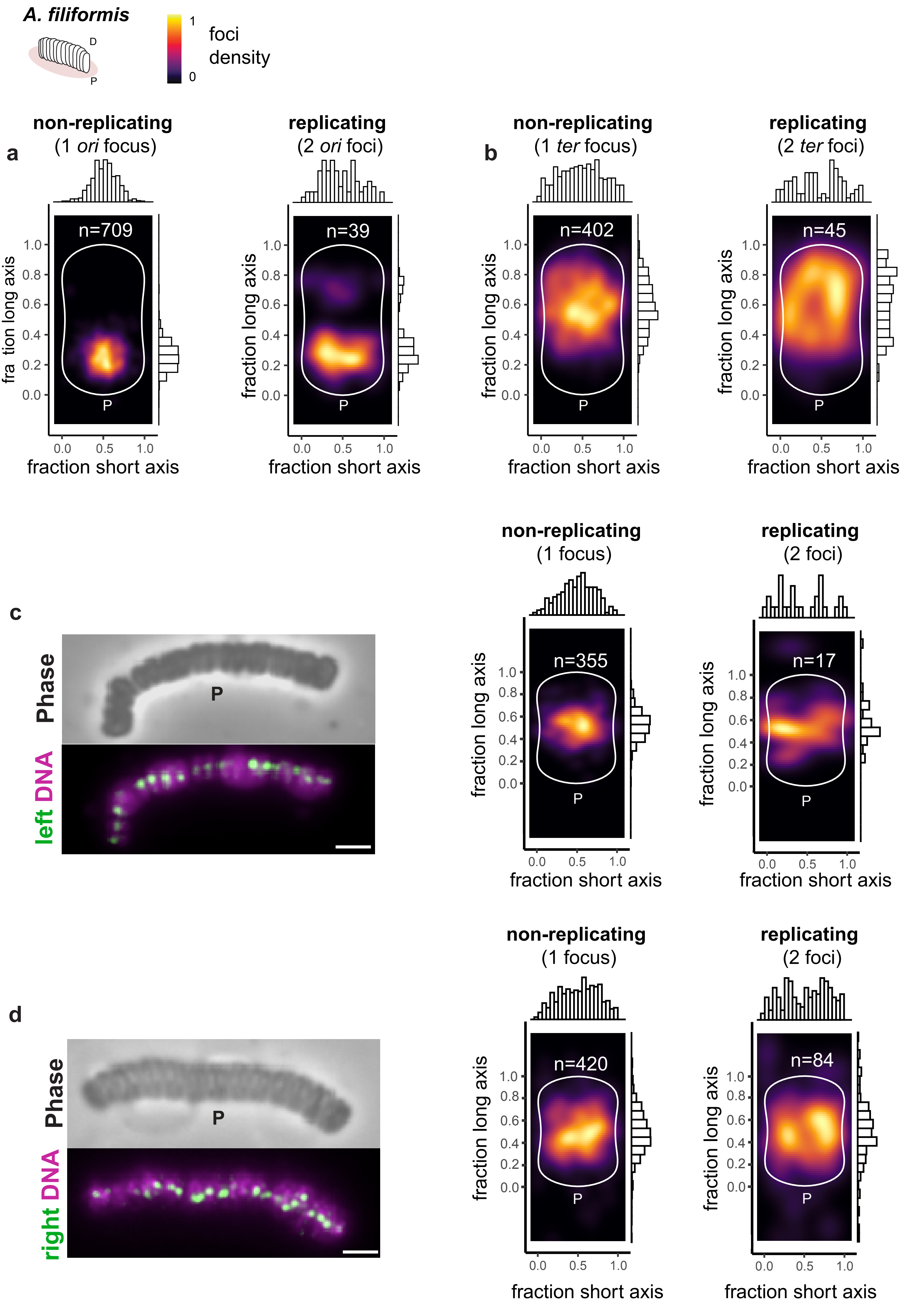

### Supplementary Fig.3

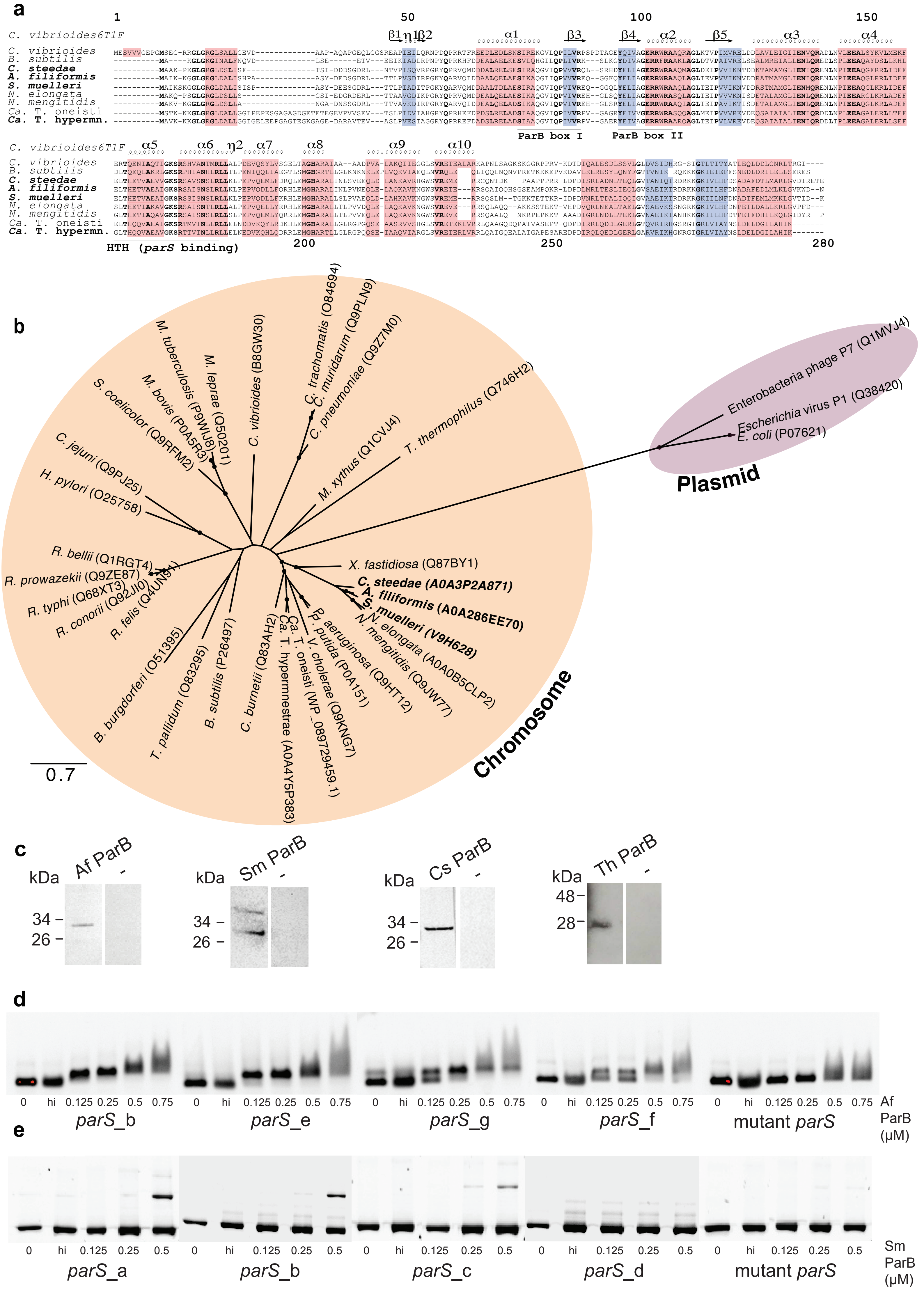

### Supplementary Fig.4

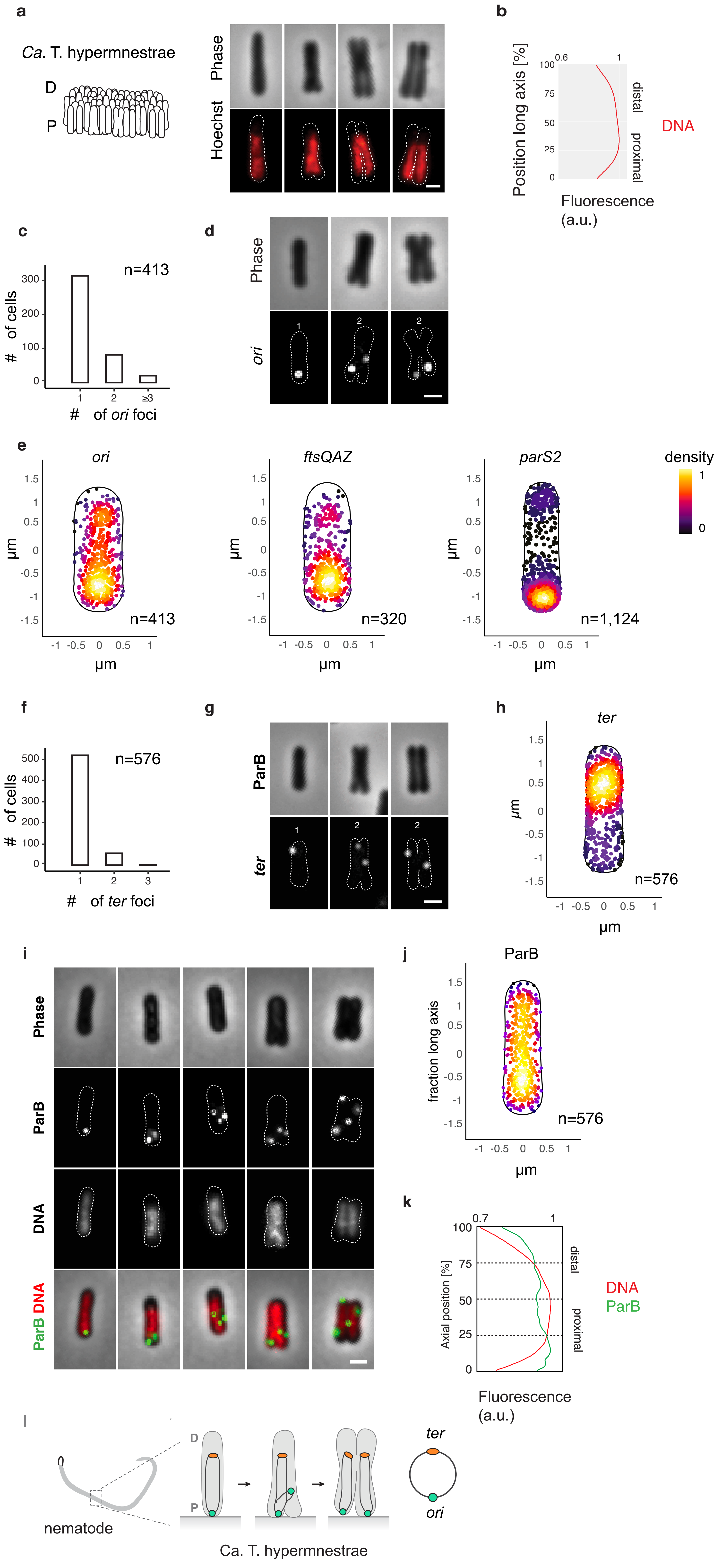

### Supplementary Fig.5

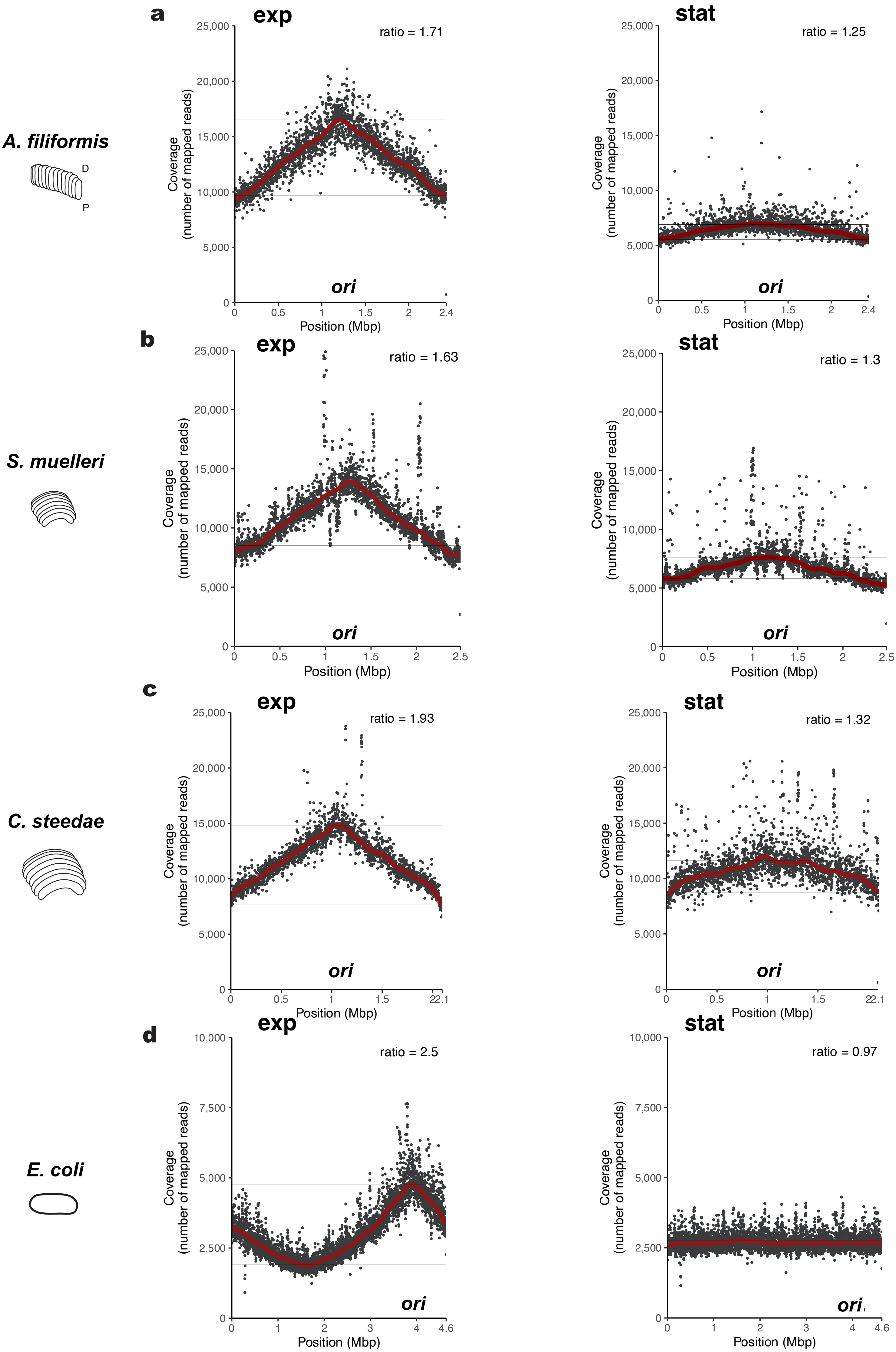

### Supplementary Fig.6

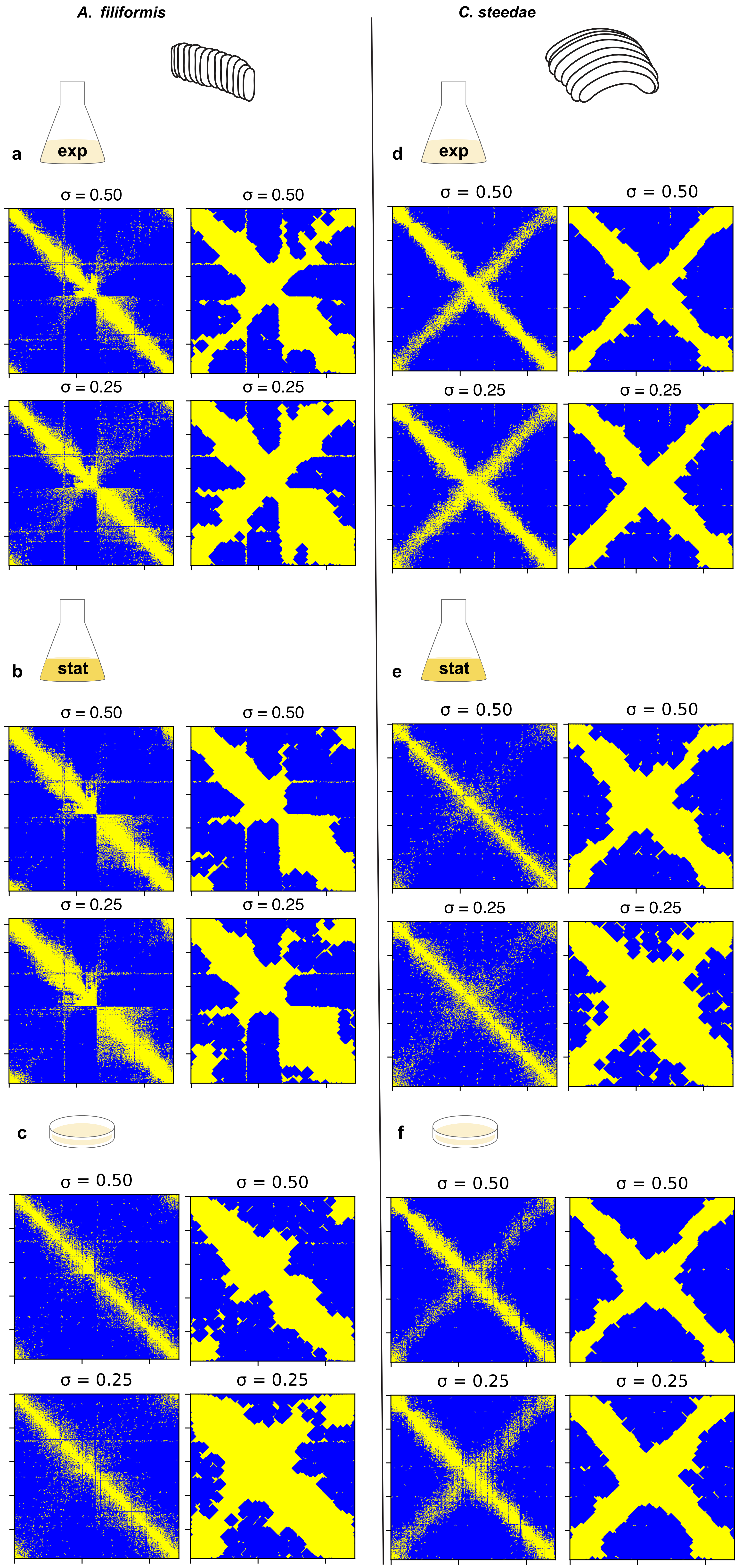

### Supplementary Fig.8

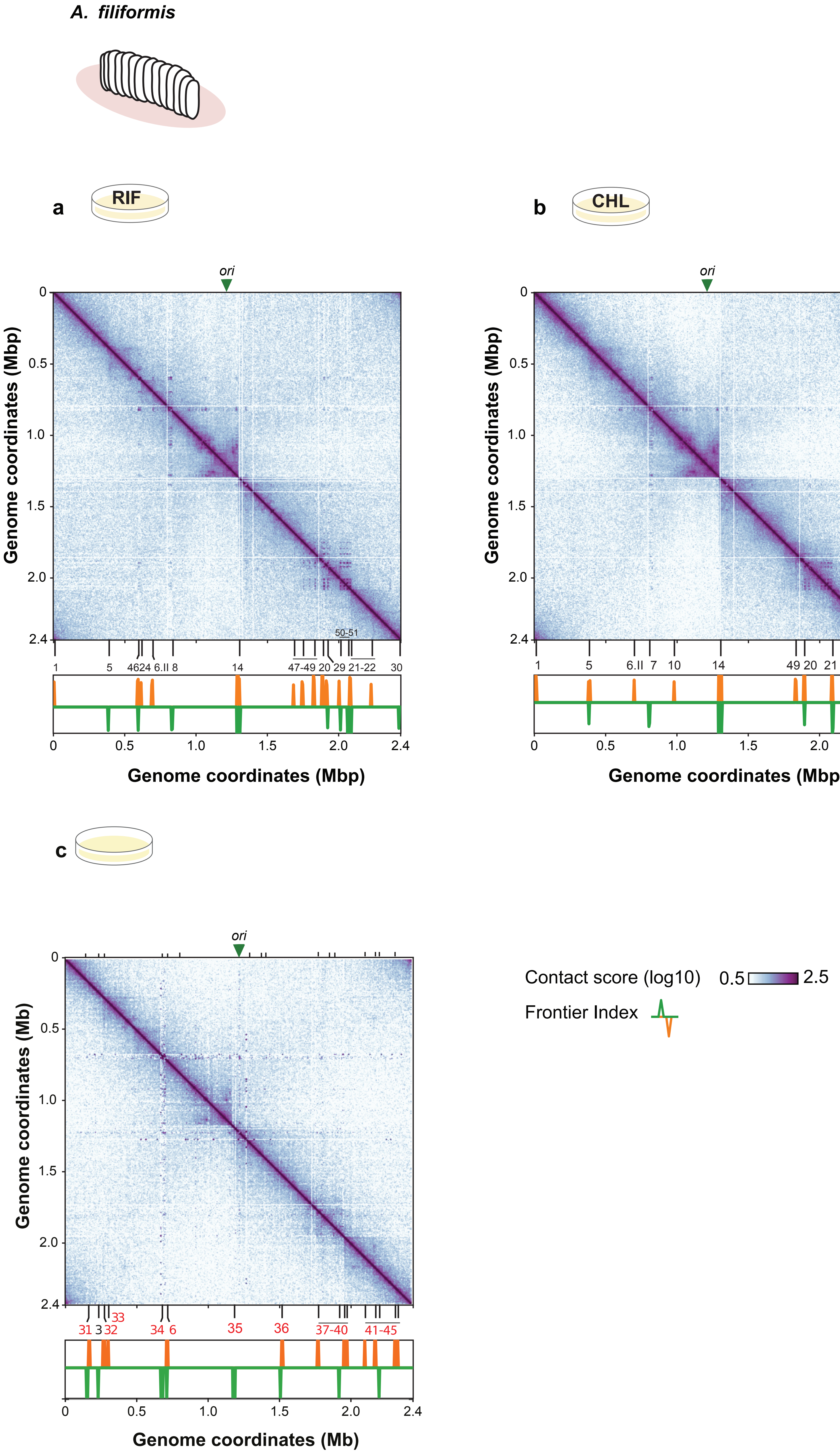
