## Supplementary Fig.9 for "Conserved lifestyle-associated chromosome architectures across mammalian symbionts"

**a**

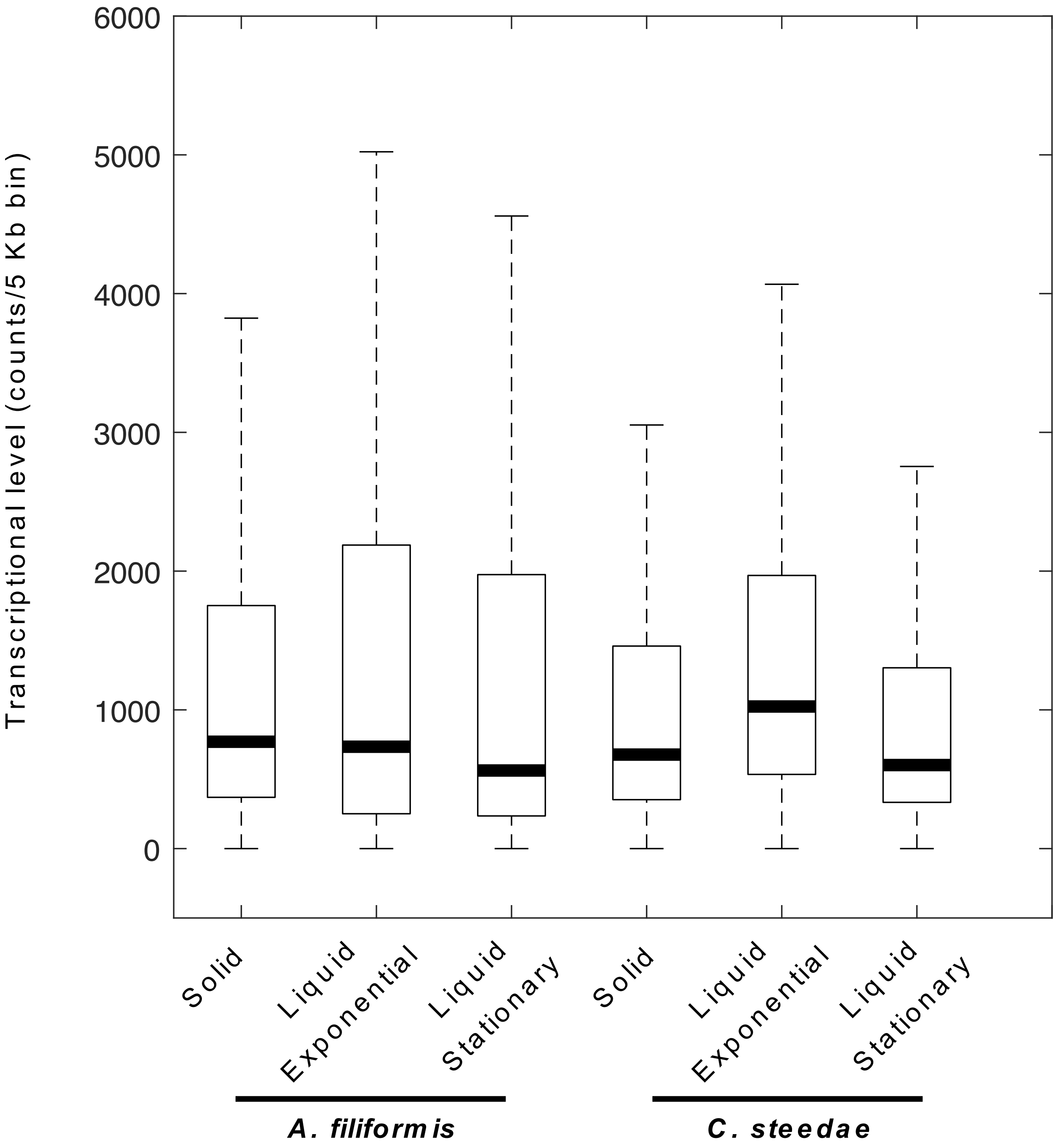


**b**

Categories of transcription level:

| | | |
|---|---|---|
| Cat_4 | Upper limit | Very high |
| Cat_3 | Between Q3 & upper limit | High |
| Cat_2 | Between median & Q3 | Medium |
| Cat_1 | Between Q1 & median | Low |
| Cat_0 | Between 0 and Q1 | No/poor |

**c**

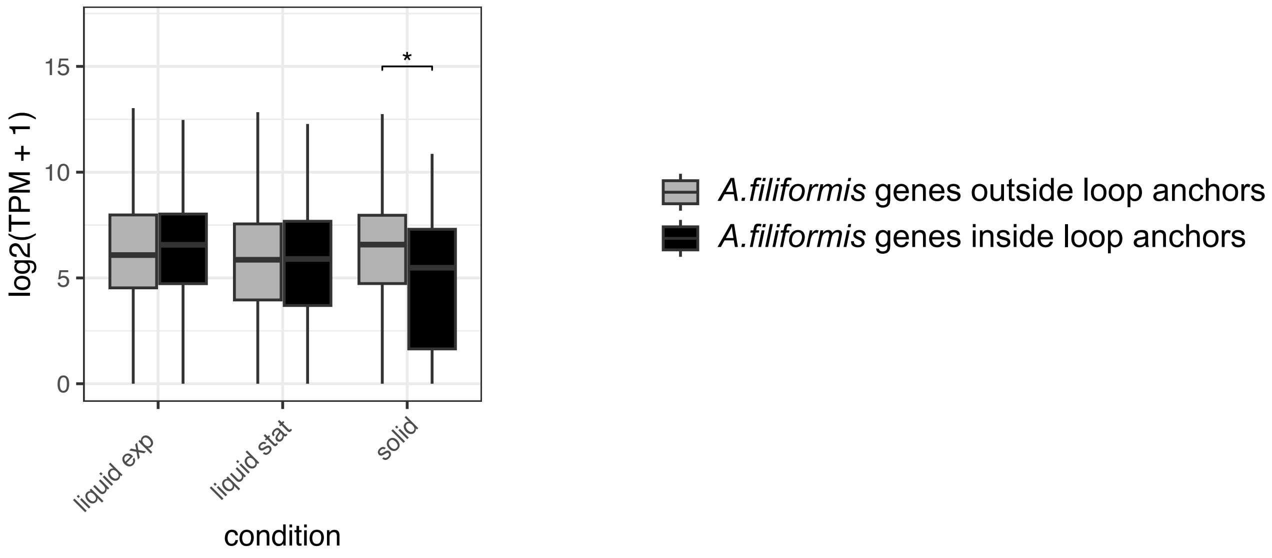
