## Supplementary Tables 1-10 for "Conserved lifestyle-associated chromosome architectures across mammalian symbionts"

**Supplementary Information**

**Table S1.** DNA FISH and immunostaining foci counts.

| **Species** | ***ori* foci per cell (%)** | | | | ***ter* foci per cell (%)** | | | |
|---|---|---|---|---|---|---|---|---|
| | **1** | **2** | **3** | **>4** | **1** | **2** | **3** | **>4** |
| ***Candidatus* T. hypermnestrae** ($n_{ori}$ =413; $n_{ter}$ =576) | 76 | 20 | 5 | - | 90 | 10 | - | - |
| ***A. filiformis*** ($n_{ori}$ =748; $n_{ter}$ =447) | 95 | 5 | - | - | 90 | 10 | - | - |
| ***S. muelleri*** ($n_{ori}$ =188; $n_{ter}$ =293) | 28 | 43 | 21 | 8 | 25 | 32 | 21 | 22 |
| ***C. steedae*** ($n_{ori}$ =215; $n_{ter}$ =236) | 27 | 43 | 22 | 8 | 25 | 32 | 22 | 21 |
| | **left arm foci per cell (%)** | | | | **right arm foci per cell (%)** | | | |
| | **1** | **2** | **3** | **>4** | **1** | **2** | **3** | **>4** |
| ***A. filiformis*** ($n_{left}$=373, $n_{right}$=519) | 95 | 5 | - | - | 81 | 16 | 3 | - |

| | **5X*parS*_c foci per cell (%)** | | | |
|---|---|---|---|---|
| | **1** | **2** | **3** | **>4** |
| ***S. muelleri*** (n=331) | 20 | 57 | 18 | 5 |
| | **ParB foci per cell (%)** | | | |
| | **1** | **2** | **3** | **>4** |
| ***A. filiformis*** (n=611) | 87 | 13 | - | - |
| ***S. muelleri*** (n=277) | 29 | 35 | 21 | 15 |
| ***C. steedae*** (n=399) | 5 | 15 | 23 | 57 |

**Table S2. Description of *A. filiformis* chromatin frontiers.**

| LIQUID MEDIUM, EXPONENTIAL PHASE | | | | | | |
|---|---|---|---|---|---|---|
| **#[1]** | **start** | **end** | **type** | **Growth condition specificity[2] (± 1 bin)** | **Proteins encoded by CAT4 genes[3], rRNA genes[4] and *parS* sites[5] (contained in ± 1 bin)** | **Membrane, periplasmic or extracellular proteins[6] (contained in ± 1 bin)** |
| **1** | 1 | 1 | down | exponential and stationary | ND | Ppk |
| **2** | 42 | 42 | down | exponential | Anr, unknown function | PurK, 2X unknown function, cytochrome c5, PlsY |
| **3** | 44 | 46 | up & down | exponential and solid | Unknown function | FtsN, colicin V production protein, RarD, Lgt, HisJ, 1x unknown function |
| **4** | 58 | 58 | down | exponential | Csd2 | ND |
| **5** | 76 | 78 | up & down | exponential and stationary | Unknown function, LipA, ferredoxin | Unknown function, aerotaxis receptor, **Atp**FEBA, FixG |
| **6** | 141 | 142 | up & down | all conditions | ND | Ppi, PetCBA, unknown function |
| **7** | 159 | 162 | up & down | exponential and stationary | Fructokinase, EngB, 2X unknown function, multidrug transporter, YahK, 5S/16S/23S rRNA | Amidase, PgsBA, cytochrome c oxidase cbb3-type, 2X unknown function, |
| **8** | 167 | 167 | up | exponential and stationary | LysR-family transcriptional regulator, Tmk, YciO, 2X unknown function, LysA | Psd, SEL1 repeat-containing protein |
| **9** | 189 | 190 | down | exponential and stationary | ND | **Nuo**NMLKJH, 3X unknown function |
| **10** | 196 | 197 | up | exponential and stationary | Unknown function | Tpx, 4X unknown function, **BamA** |
| **11** | 219 | 221 | up & down | exponential and stationary | Nicotinamidase/pyrazinamidase, ErfK/SrfK, HexR | Thiamine transport system permease protein, unknown function |
| **12** | 229 | 229 | up & down | exponential and stationary | Unknown function (membrane protein), Ser tRNA | Unknown function, ABC-type Fe3+/spermidine/putrescine transport system, **PilY1** |
| **13** | 243 | 243 | up | exponential | ND | 2X unknown function, SecE, CpdB, recombinase |
| **14** | 259 | 262 | up & down | exponential and stationary | Peptide/bleomycin uptake transporter, LTXXQ-motif family protein; 5S/16S/23S rRNA; *parS*_7 (dialect g) | LutA, **ComEA**, peptide/bleomycin uptake transporter, LTXXQ-motif family protein |
| **15** | 266 | 266 | up | exponential | TamA, unknown function, 5S/16S/23S rRNA | TamA, unknown function, AqpZ |
| **16** | 281 | 281 | up & down | exponential | YceF, unknown function, 5S/16S/23S rRNA | DltB, unknown function, RnhB |
| **17** | 340 | 340 | up | exponential | ND | NqrBD, RsxA, unknown function, LptD |
| **18** | 370 | 370 | down | exponential | TilS, phosphohistidine phosphatase; 5S/16S/23S rRNA | Unknown function, 2X amino acid ABC transporter, GltL, **polar amino acid transporter** |
| **19** | 373 | 375 | up & down | exponential | PyrL, putative ABC transport system substrate-binding protein, unknown function | Dual specificity phosphatase, unknown function |
| **20** | 377 | 378 | up | exponential | MurU, YdzA, unknown function, PykA, 5-formyltetrahydrofolate cyclo-ligase, LysR-family transcriptional regulator; 5S/16S/23S rRNA | MurU, YdzA, FtsK |
| **21** | 418 | 418 | up & down | exponential and stationary | Pcp, unknown function, HTH-type transcriptional regulator, haloacid dehalogenase-like hydrolase, phosphoglycolate phosphatase | TPR repeat-containing protein, SodA, Pcp, SpuE, autotransporter barrel domain-containing protein |
| 22 | 448 | 449 | up & down | exponential | Transposase, transglutaminase-like superfamily protein | 2X unknown function, phospholipase A1, **Sdh**ABCD, transglutaminase-like protein |

| LIQUID MEDIUM, STATIONARY PHASE | | | | | | |
|---|---|---|---|---|---|---|
| **#[1]** | **start** | **end** | **type** | **Growth condition specificity[2] (± 1 bin)** | **Proteins encoded by CAT4 genes[3], rRNA genes[4] and *parS* sites[5] (contained in ± 1 bin)** | **Membrane, periplasmic or extracellular proteins[6] (contained in ± 1 bin)** |
| **1** | 1 | 3 | up & down | exponential and stationary | ND | Ppk, DgkA, OmpR, O-antigen ligase, unknown function |
| **5** | 76 | 79 | up & down | exponential and stationary | 2X unknown function, LipA, ferredoxin | 2X Unknown function, aerotaxis receptor, AtpFEBA, FixG, FtsH |
| **23** | 109 | 111 | up & down | stationary | 5X unknown function, **Ppi**B | 3X unknown function, thiol:disulfide interchange protein |
| **24** | 124 | 124 | down | stationary | PrfA, 2X unknown function, YafQ, RelB | Transporter, unknown function, Era |
| 6 | 141 | 141 | up | all conditions | ND | **Ppi**, PetCBA |
| **7** | 161 | 162 | down | exponential and stationary | EngB, 2X unknown function, multidrug transporter, YahK; 5S/16S/23S rRNA | Cytochrome c oxidase cbb3-type, 2X unknown function |
| **8** | 167 | 167 | up | exponential and stationary | LysR, unknown function, LysA | Psd, SEL1 |
| **9** | 188 | 188 | down | exponential and stationary | ND | 3X unknown function, NuoNMLKJH |
| **10** | 196 | 197 | up | exponential and stationary | 2X unknown function | Tpx, 4X unknown function, **Bam**A |
| **25** | 203 | 203 | down | stationary | Unknown function, RNaseP, IspU | 3X unknown function, serine protease, SCO1/2, CdsA |
| **26** | 216 | 216 | down | stationary | Eda, RDD family protein, YrdA, MviN, Nicotinamidase/pyrazinamidase | RDD family protein, MviN, RTX-toxin related |
| **11** | 220 | 222 | up & down | exponential and stationary | ErfK/SrfK, HexR | Thiamine transport system permease, unknown function |
| **27** | 225 | 225 | up | stationary | TsaC | OmpA |
| **12** | 228 | 228 | up & down | exponential and stationary | ND | ND |
| **28** | 251 | 252 | down | stationary | RuvA; *parS*_1-4 (dialects f, e, b, f) | ABC transporter, 2X unknown function |
| **14** | 256 | 263 | up & down | exponential and stationary | Unknown function, LldF, peptide/bleomycin uptake transporter, LTXXQ motif family protein, 5S/16S/23S rRNA; *parS*_7 (dialect g) | LutA, **ComEA**, peptide/bleomycin uptake transporter, LTXXQ motif family protein, RhtA |
| **29** | 386 | 386 | down | stationary | TldD, serine aminopeptidase, alpha/beta hydrolase | Alpha/beta hydrolase, pertactin |
| **21** | 417 | 418 | up & down | exponential and stationary | haloacid dehalogenase-like hydrolase, phosphoglycolate phosphatase | TPR repeat-containing protein, Soda, pcp, SpuA, autotransporter barrel domain-containing protein |
| **30** | 486 | 486 | down | stationary | IscA, TatD, 3X unknown function, CysE | IscU, 2X unknown function, NhaD, **Ubi**A |

| SOLID MEDIUM | | | | | | | | |
|---|---|---|---|---|---|---|---|---|
| #[1] | start | end | type | Growth condition specificity[2] (± 1 bin) | RIF | CHL | Proteins encoded by CAT4 genes[3], rRNA genes[4] and *parS* sites[5] (contained in ± 1 bin) | Membrane, periplasmic or extracellular proteins[6] (contained in ± 1 bin) |
| **31** | 31 | 35 | up & down | solid | **N** | **N** | AccA, Rpe, AcpS, 2X unknown function, **MFS transporter**, **PBP7**, Apt, AroQ, Ung, Adk, UDP-glucose:protein N-beta-glucosyltransferase | **MFS transporter**, **PBP7**, ArnT, FkpA, unknown function |
| **3** | 47 | 47 | down | exponential and solid | **N** | **N** | ND | HisJ, unknown function, **MrdBA**, MreD |
| **32** | 54 | 56 | up | solid | **N** | **N** | Ddc, unknown function, **M23 peptidase**, IlvA, Csd2 | 3X unknown function, **LolC**D, DacA |
| **33** | 61 | 61 | up | solid | **N** | **N** | ND | 4X unknown function, Signal transduction histidine kinase involved in nitrogen fixation, septation protein A |
| **34** | 135 | 137 | down | solid | **N** | **N** | hldE, YggH, 3',5'-nucleoside bisphosphate phosphatase | **Opp**B, GltL, unknown function |
| **6** | 143 | 144 | up & down | all conditions | **N** | **N** | FAD/FMN-containing dehydrogenase | Ppi, PetCBA, 3X unknown function, EfeO/EfeM, EfeB, **UraA** |
| **35** | 236 | 238 | down | solid | **N** | **N** | ND | Unknown function, azaleucine resistance, CBS domain-containing protein, **polar amino acid transport system** substrate-binding protein |
| **36** | 302 | 305 | up & down | solid | **N** | **N** | 2X unknown function, LysM, CysI | 2X unknown function |
| **37** | 354 | 355 | up | solid | **N** | **N** | **FHA** | YrbEF, YidC, **hemolysin** activation/secretion protein, **FHA** |
| **38** | 384 | 384 | down | solid | **N** | **N** | Serine aminopeptidase, alpha/beta hydrolase | RmpM, MacB, Membrane fusion protein, macrolide-specific efflux system, alpha/beta hydrolase |
| **39** | 391 | 391 | up | solid | **N** | **N** | ND | Pertactin, 2X unknown function, YdgI, HemYX |
| **40** | 394 | 395 | up | solid | **N** | **N** | put. DNA primase/helicase, unknown function, tmRNA | OmpR-family protein |
| **41** | 420 | 420 | up | solid | **N** | **N** | HTH-type transcriptional regulator, haloacid dehalogenase-like hydrolase, phosphoglycolate phosphatase, YiaA | **Spu**E, autotransporter barrel domain-containing protein, **FabI**, Hemoglobin/transferrin/lactoferrin receptor protein, YiaA, 2X unknown function, YclN |
| **42** | 434 | 435 | up | solid | **N** | **N** | PseF, RsmG | 3X unknown function, MglBAC, **polar amino acid transporter**, GlnB |
| **43** | 440 | 440 | down | solid | **N** | **N** | PseB, membrane protein implicated in regulation of membrane protease activity, TrmH | 3X unknown function, membrane protein implicated in regulation of membrane protease activity, **YqiK** |
| **44** | 462 | 462 | up | solid | **N** | **N** | **Mlt**G, MetF, TrpB, unknown function | NlpD, YhjE, holin-like protein, 2X unknown function |
| **45** | 466 | 466 | up | solid | **N** | **N** | 3X unknown function, AroG, CtrD | 3X unknown function, CtrBC |

| SOLID MEDIUM + RIFAMPICIN (RIF) | | | | | |
|---|---|---|---|---|---|
| **#[1]** | **start** | **end** | **type** | **Present in solid medium?** | **Present in CHL?** |
| **1** | 2 | 2 | up | N | Y |
| **5** | 78 | 78 | down | N | Y |
| **46** | 119 | 120 | up & down | N | N |
| **24** | 124 | 124 | up | N | N |
| **6.II** | 139 | 140 | up | N | Y |
| **8** | 167 | 168 | down | N | N |
| **14** | 259 | 263 | up & down | N | Y |
| **47** | 338 | 338 | up | N | N |
| **48** | 350 | 351 | up | N | N |
| **49** | 366 | 367 | up | N | Y |
| **20** | 378 | 379 | up | N | Y |
| **29** | 384 | 386 | up & down | N | N |
| **50** | 402 | 404 | up & down | N | N |
| **51** | 414 | 414 | down | N | N |
| **21** | 417 | 419 | up & down | N | Y |
| **22** | 447 | 447 | up | N | Y |
| **30** | 486 | 486 | down | N | Y |

| SOLID MEDIUM + CHLORAMPHENICOL (CHL) | | | | | |
|---|---|---|---|---|---|
| **#[1]** | **start** | **end** | **type** | **Present in solid medium?** | **Present in RIF?** |
| **1** | 2 | 3 | up | N | Y |
| **5** | 76 | 78 | up & down | N | Y |
| **6.II** | 140 | 140 | up | N | Y |
| **7** | 161 | 162 | down | N | N |
| **10** | 196 | 196 | up | N | N |
| **14** | 259 | 263 | up & down | N | Y |
| **49** | 366 | 367 | up | N | Y |
| **20** | 378 | 379 | up & down | N | Y |
| **21** | 417 | 419 | up & down | N | Y |
| **52** | 444 | 444 | down | N | N |
| **22** | 447 | 449 | up & down | N | Y |
| **30** | 486 | 487 | down | N | Y |

[1]Frontiers are numbered sequentially (from liquid medium exponential phase to liquid medium stationary phase, to solid medium). See Supplementary Data Set 1 for genomic position of frontiers, gene names, intracellular localization, category of expression and Supplementary Data Set 3 for annotation.

[2]All conditions frontiers: frontiers that share at least 1 bin irrespective of the growth condition. Exponential and solid frontiers: frontiers that share at least 1 bin in liquid medium exponential phase and on solid medium. Exponential and stationary frontiers: frontiers that share at least 1 bin in liquid medium exponential phase and stationary phase.

[3]Proteins encoded by genes that are CAT4 at least in the respective condition. See Supplementary Data Set 1 for genomic position of frontier-generating bins, gene names, intracellular localization category of expression.

[4]See Table S3 for rRNA genes genomic position.

[5]See Table S3 for *parS* genomic position.

[6]Proteins are defined as membrane proteins based on PSORTb. See Supplementary Data Set 1 for genomic position of frontier-generating bins, gene names, intracellular localization, category of expression.

RIF: rifampicin (25 µg/ml, 30 min); CHL: chloramphenicol (25 µg/ml, 30 min). Y: frontier present. N: frontier absent.

**Table S3. Genomic positions and bins of *A. filiformis* rRNA genes and *parS* sites**

| rRNA operon # | rRNA gene | start | end | bin | |
|---|---|---|---|---|---|
| **1** | 5S | 793,150 | 793,260 | 159 | |
| | 23S | 793,397 | 796,282 | 159 \| 160 | |
| | 16S | 797,108 | 798,634 | 160 | |
| **2** | 16S | 1,300,206 | 1,301,732 | 261 | |
| | 23S | 1,302,558 | 1,305,443 | 261 \| 262 | |
| | 5S | 1,305,580 | 1,305,690 | 262 | |
| **3** | 16S | 1,318,439 | 1,319,965 | 264 | |
| | 23S | 1,320,791 | 1,323,676 | 265 | |
| | 5S | 1,323,813 | 1,323,923 | 265 | |
| **4** | 16S | 1,392,778 | 1,394,304 | 279 | |
| | 23S | 1,395,130 | 1,398,015 | 280 | |
| | 5S | 1,398,152 | 1,398,262 | 280 | |
| **5** | 16S | 1,854,102 | 1,855,628 | 371 \| 372 | |
| | 23S | 1,856,454 | 1,859,339 | 372 | |
| | 5S | 1,859,476 | 1,859,586 | 372 | |
| **6** | 16S | 1,877,147 | 1,878,673 | 376 | |
| | 23S | 1,879,576 | 1,882,461 | 376 \| 377 | |
| | 5S | 1,882,598 | 1,882,708 | 377 | |
| ***parS* site #** | **dialect** | **start** | **end** | **bin** | **sequence** |
| ***parS1*** | f | 1,258,860 | 1,258,875 | 251 | TGTTTCACGTGAAACA |
| ***parS2*** | e | 1,258,901 | 1,258,916 | 251 | TGTTTCACGTGAAACG |
| ***parS3*** | b | 1,259,463 | 1,259,478 | 251 | **TGTTTCACATGAAACA** |
| ***parS4*** | f | 1,264,185 | 1,264,200 | 252 | TGTTTCACATGAAACA |
| ***parS5*** | g | 1,271,279 | 1,271,294 | 254 | TGTTTCACATGAAACC |
| ***parS6*** | f | 1,271,393 | 1,271,408 | 254 | TGTTTCACATGAAACA |
| ***parS7*** | g | 1,296,548 | 1,296,563 | 259 | TGTTTCACATGAAACC |

**Table S4. Description of *C. steedae* chromatin frontiers.**

| LIQUID MEDIUM, EXPONENTIAL PHASE | | | | | | |
|---|---|---|---|---|---|---|
| **#[1]** | **start** | **end** | **type** | **Growth condition specificity[2] (± 1 bin)** | **Products of CAT4 genes[3], rRNA genes[4] and *parS* sites[5] (contained in ± 1 bin)** | **Membrane, periplasmic or extracellular proteins[6] (contained in ± 1 bin)** |
| **1** | 2 | 3 | up | exponential | phosphate acetyltransferase | DUF2788 domain-containing protein, LplX, NhaC, phage holin family protein |
| **2** | 26 | 26 | up & down | exponential | SucAB, **Sdh**AB, AspC | **Sdh**ABCD |
| **3** | 35 | 35 | down | all conditions | **PilY1**; *parS*_1 (dialect h) | Isoprenylcysteine carboxyl methyltransferase family protein; MFS transporter, **PilY1** |
| **4** | 43 | 46 | up & down | all conditions | LldD, 2X hemolysin, GatA; *parS*_2 (dialect j) | OmpA, MotA/TolQ/ExbB proton channel family protein, **Ubi**B, 2X hemolysin, MrdAB, MreCD |
| **5** | 181 | 181 | up & down | exponential and solid | MetE, **BamE**[7]; 5S rRNA,16S rRNA, 23S rRNA | DUF1275 domain-containing protein, conserved membrane protein of unknown function, DUF4184 family protein, **BamE**[7] |
| **6** | 184 | 184 | up & down | all conditions | BamE[7], YadA-like adhesin, methyltransferase regulatory domain-containing protein | BamE[7], YadA-like adhesin[8], energy-coupling factor ABC transporter permease, LspA |
| **7** | 190 | 191 | up & down | exponential and solid | 25 ribosomal proteins, InfA, SecY | nonpolar-amino-acid-transporting ATPase, SecY, **polar amino acid transporter** |
| **8** | 210 | 210 | down | exponential | HemL, 1 ribosomal protein, YceD, TetR/AcrR, NmoA | conserved membrane protein of unknown function |
| **9** | 218 | 218 | up & down | exponential | 5S rRNA, 16S rRNA, 23S rRNA | Zinc/Manganese transport system permease protein, Uncharacterized transporter HI_0223, conserved membrane protein of unknown function, Zinc/Manganese transport system ATP-binding protein, DUF2339 domain-containing protein |
| **10** | 246 | 249 | up & down | all conditions | 2 ribosomal proteins, TrmD, alpha-glucan phosphorylase, peptidase M23, roadblock/LC7 domain-containing protein; 5S rRNA, 16S rRNA, 23S rRNA | Azu, alpha-glucan phosphorylase, CcpA, FabF, polar amino acid transport system substrate-binding protein |
| **11** | 272 | 274 | up & down | exponential | **Nuo**AH, LolA, HsdM | **Nuo**HJKLN, Nqo, DUF2818 family protein, LolA |
| **12** | 300 | 302 | up & down | all conditions | 1 ribosomal protein; 5S rRNA, 16S rRNA, 23S rRNA | CstA, **ComEA**, UraA, conserved protein of unknown function, conserved membrane protein of unknown function, AMP-binding protein, conserved membrane protein of unknown function, 1-acyl-sn-glycerol-3-phosphate acyltransferase |
| **13** | 332 | 333 | up & down | all conditions | Tig | EpsL, polysaccharide biosynthesis protein, MrcA |
| **14** | 364 | 365 | up & down | exponential | Zn ribbon domain-containing protein, Mtn | **Ppi**, DUF485 domain-containing protein, acetate/glycolate:cation symporter membrane-associated protein, anhydro-N-acetylmuramic acid kinase, conserved exported protein of unknown function (extracellular) |
| **15** | 369 | 369 | up | exponential | L-lactate permease, Adh | L-lactate permease, tyrosine-protein kinase Etk/Wzc |
| **16** | 386 | 387 | up & down | exponential and solid | **Atp**EFHEGDC | ATP F0F1 synthase subunit I, **Atp**ACB, P-type Cu2+ transporter |
| **17** | 405 | 405 | up | exponential | 3 ribosomal proteins, PriB, HemY, CopA | ferredoxin/flavodoxin---NADP+ reductase, YgjV, CobA, Prepilin-type N-terminal cleavage/methylation domain-containing protein |
| **18** | 418 | 418 | up & down | all conditions | Lpd | Kdo2-lipid IVA lauroyltransferase/acyltransferase, choline dehydrogenase, BetT |

| LIQUID MEDIUM, STATIONARY PHASE | | | | | | |
|---|---|---|---|---|---|---|
| #[1] | start | end | Type | Growth condition specificity[2] | Products of CAT4 genes[3], rRNA genes[4] and *parS* sites[5] (contained in ± 1 bin) | Membrane, periplasmic or extracellular proteins[6] (contained in ± 1 bin) |
| **3** | 34 | 34 | down | all conditions | PilY1; *parS*_1 (dialect h) | LysO; PilY1 |
| **4** | 44 | 46 | up & down | all conditions | 2X hemolysins, GatA; *parS*_2 (dialect j) | **Ubi**B, 2X hemolysins, MrdBA, MreDC |
| **19** | 153 | 153 | up | stationary | Nitroreductase family protein, GrpE, Odhl | YedIH, TatCBA, putative outer membrane protein NMB0088 |
| **20** | 170 | 170 | down | stationary | Adhesin, NmoA | NarYG, RsfS, conserved membrane protein of unknown function |
| **6** | 184 | 184 | down | all conditions | BamE[7] | BamE[7], YadA-like adhesin[8], energy-coupling factor ABC transporter permease, LspA |
| **21** | 232 | 232 | up | stationary | ND | conserved protein of unknown function, hemolysin, Blc, PotIHG, **Bam**A, putative Zinc metalloprotease NMA0084, **Ppi** |
| **10** | 248 | 248 | up & down | all conditions | roadblock/LC7 domain-containing protein, 1 ribosomal protein, TrmD; 5S rRNA, 16S rRNA, 23S rRNA | CcpA, FabF |
| **12** | 302 | 302 | up | all conditions | Unknown function; 23S rRNA, 5S rRNA | **ComEA**, UraA, conserved protein of unknown function, conserved membrane protein of unknown function, AMP-binding protein, conserved membrane protein of unknown function, PlsC |
| **13** | 332 | 333 | up & down | all conditions | Tig | EpsL, polysaccharide biosynthesis protein, MrcA |
| **15** | 370 | 371 | up | stationary | Adh, prepilin-type N-terminal cleavage/methylation domain containing protein, unknown function, PilVW, FimT, DnaB | L-lactate permease, tyrosine-protein kinase Etk/Wzc, GfcE, LolDC, ParA, UshA |
| **22** | 399 | 399 | down | stationary | Mdh | Lnt, YegL, Rhomboid family intramembrane serine protease, phosphate:Na+ symporter |
| **18** | 417 | 419 | down | all conditions | Lpd, Pcm | choline dehydrogenase, BetT, multiple antibiotic resistance protein |

| SOLID MEDIUM | | | | | | |
|---|---|---|---|---|---|---|
| **#[1]** | **start** | **end** | **Type** | **Growth condition specificity[2]** | **Products of CAT4 genes[3], rRNA genes[4] and *parS* sites[5] (contained in ± 1 bin)** | **Membrane, periplasmic or extracellular proteins[6] (contained in ± 1 bin)** |
| **3** | 31 | 37 | down | all conditions | Unknown function, PilY1, **PBP7**, unknown function; *parS*_1 (dialect h) | NupX, **FHA**, LysO, Isoprenylcysteine carboxyl methyltransferase family protein, **MFS transporter**, S-adenosylmethionine uptake transporter, **PBP7** |
| **4** | 44 | 45 | up & down | all conditions | 2X **hemolysin**; *parS*_2 (dialect j) | UbiB, 2X **hemolysin**, **MrdBA** |
| **23** | 174 | 174 | up | solid | ND | RecC, **polar amino acid transporter**, KefC, **Spu**D |
| **5** | 181 | 181 | down | exponential and solid | BamE[7]; 5S rRNA, 16S rRNA, 23S rRNA | BamE[7], DUF1275 domain-containing protein, conserved membrane protein of unknown function, DUF4184 family protein |
| **6** | 185 | 185 | down | all conditions | ND | YadA-like adhesin[8], energy-coupling factor ABC transporter permease, LspA, Phage coat protein |
| **7** | 190 | 190 | down | exponential and solid | DUF4124, RpoA, SecY, 10 ribosomal proteins | nonpolar-amino-acid-transporting ATPase, SecY, **polar amino acid transporter** |
| **24** | 193 | 194 | up | solid | 10 ribosomal proteins, TufB | YocR, ZupT, FtsX, FtsE, Recombinase |
| **10** | 246 | 248 | down | all conditions | Unknown function, alphaglucanphosphorylase, ATP/GTP binding protein, **M23 peptidase**; 5S rRNA, 16S rRNA, 23S rRNA | metallophosphoesterase, Azu, Alpha-glucan phosphorylase, CcpA, **Fab**F |
| **25** | 258 | 259 | down | solid | Unknown function | 2X RDD family protein, **OppA**, CdiB, **FHA** |
| **26** | 262 | 262 | up | solid | ND | conserved membrane protein of unknown function, **MltB** |
| **12** | 302 | 302 | up | all conditions | Unknown function; 5S rRNA, 16S rRNA, 23S rRNA | ComEA, **UraA**, 3X unknown function, AMP-binding protein, PlsC |
| **27** | 328 | 329 | down | solid | PIN7 domain-containing protein | **YqiK**, YbjQ, 3X unknown function, WecA |
| **13** | 334 | 334 | up & down | all conditions | PilNOPQ | PilNOPQ, EpsL, polysaccharide biosynthesis protein, MrcA |
| **28** | 338 | 338 | up | solid | IgA peptidase, 2X unknown function | LepA, signal peptidase I, Iga |
| **29** | 373 | 373 | up | solid | ND | **LolC**, ParA, UshA, Mtr, **MFS transporter** |
| **16** | 388 | 388 | up | exponential and solid | ND | P-type Cu2+ transporter, Mg transporter, membrane protein of unknown function, cardiolipin synthase A/B |
| **18a[1]** | 416 | 416 | down | all conditions | PrpC | LysE, 2X membrane protein of unknown functionm, bacterial/archaeal transporter family-2 protein, Kdo2-lipid IVA lauroyltransferase/acyltransferase |
| **18b[1]** | 419 | 419 | up & down | all conditions | Lpd | choline dehydrogenase, BetT, multiple antibiotic resistance protein |

[1]Frontiers are numbered sequentially (from liquid medium exponential phase to liquid medium stationary phase, to solid medium). In liquid medium exponential phase, bin 418 is involved in one frontier (frontier #18); in liquid medium stationary phase, bins 417 through 419 are involved in one frontier (frontier #18); on solid medium, bin 416 is involved in frontier 18a and bin 419 is involved in frontier 18b. See Supplementary Data Set 2 for genomic position of frontiers, gene names, intracellular localization, category of expression and Supplementary Data Set 3 for annotation.

[2]All conditions frontiers: frontiers that share at least 1 bin irrespective of the growth condition. Exponential and solid frontiers: frontiers that share at least 1 bin condition in exponential phase and on solid medium. Exponential and stationary frontiers: frontiers that share at least 1 bin condition in exponential phase and stationary phase.

[3]Proteins encoded by genes that are CAT4 at least in the respective condition. See Supplementary Data Set 2 for genomic position of frontier-generating bins, gene names, intracellular localization, category of expression.

[4]See Table S3 for rRNA genes genomic position.

[5]See Table S3 for *parS* genomic position.

[6]Proteins are defined as membrane proteins based on PSORTb. See Supplementary Data Set 2 for genomic position of bins, gene names, intracellular localization, category of expression.

[7]*bamE* spans across bins 182 and 183. Therefore, it matches frontier #5 (181) and frontier #6 (184) in liquid medium exponential *C. steedae*, but only #6 in stationary phase. On solid medium, *bamE* lies between frontier #5 (181) and frontier #6 (185).

[8]The *yadA*-like adhesin gene spans across bins 183 and 185.

**Table S5. Genomic positions and bins of *C. steedae* rRNA genes and *parS* sites and genomic positions of *S. muelleri parS* sites.**

| ***C. steedae* rRNA operon #** | **rRNA gene** | **start** | **end** | **bin** | |
|---|---|---|---|---|---|
| **1** | 5S | 903740 | 903851 | 181 | |
| | 23S | 903928 | 906811 | 181 \| 182 | |
| | 16S | 907406 | 908934 | 182 | |
| **2** | 16S | 1079024 | 1080552 | 216 \| 217 | |
| | 23S | 1081147 | 1084031 | 217 | |
| | 5S | 1084108 | 1084219 | 217 | |
| **3** | 16S | 1231766 | 1233294 | 247 | |
| | 23S | 1233889 | 1236773 | 247 \| 248 | |
| | 5S | 1236850 | 1236961 | 248 | |
| **4** | 16S | 1498454 | 1499982 | 300 | |
| | 23S | 1500610 | 1503495 | 301 | |
| | 5S | 1503572 | 1503683 | 301 | |
| ***C. steedae parS* site #** | **dialect** | **start** | **end** | **bin** | **sequence** |
| ***parS1*** | h | 170,813 | 170,828 | 34 | **TGTTCCAAGTGGAACT** |
| ***parS2*** | i | 189,742 | 189,757 | 37 | **CGTTCCACCTGAAACG** |
| ***parS3*** | j | 226,624 | 226,639 | 45 | AGTTTCATGTGAAACG |
| ***parS4*** | k | 278,630 | 278,645 | 55 | CGTTCCACCTGCAACA |
| ***parS5*** | e | 1,023,430 | 1,023,445 | 204 | TGTTTCACGTGAAACG |
| ***parS6*** | l | 1,038,803 | 1,038,818 | 207 | AGTTTCACGTGAAACG |
| ***S. muelleri parS* site #** | **dialect** | **start** | **end** | | **sequence** |
| ***parS1*** | a | 1,215,754 | 1,215,769 | | **TGTTTCATGTGAAACA** |
| ***parS2*** | b | 1,215,793 | 1,215,808 | | TGTTTCACGTGAAACA |
| ***parS3*** | b | 1,215,824 | 1,215,839 | | TGTTTCACGTGAAACA |
| ***parS4*** | c | 1,447,496 | 1,447,511 | | CGTTCCAAGTGAAACA |
| ***parS5*** | c | 1,448,045 | 1,448,060 | | CGTTCCAAGTGAAACA |
| ***parS6*** | c | 1,448,594 | 1,448,609 | | CGTTCCAAGTGAAACA |
| ***parS7*** | c | 1,449,143 | 1,449,158 | | CGTTCCAAGTGAAACA |
| ***parS8*** | c | 1,449,692 | 1,449,707 | | CGTTCCAAGTGAAACA |
| ***parS9*** | d | 2,178,556 | 2,178,571 | | TGTTGCAAGTGTAACA |
| ***parS10*** | a | 2,345,466 | 2,345,481 | | **TGTTTCATGTGAAACA** |

**Table S6. Genomic position, gene content and *parS* site content of *A. filiformis* loop anchors.** Only genes with predicted known functions and assigned EggNOG COG class (in parenthesis) are shown. Genes in eggNOG-based Clusters of Orthologous Groups (COG) Category *Metabolism* are in green, *Information processing* are in orange and *Cellular processes and signaling* are in black. See Supplementary Data Set 4 for additional information on genes with predicted function contained in loop anchors.

| | | |
|---|---|---|
| **Exponential Loop 1 (Exp L1;** anchor 2 + anchor 2') | ***Anchor 2'*** | |
| | **Genomic position** (Mbp; bins) | 1.045–1.130; 210–226 |
| | **Genes** | *polar aa transport permease (P), ribF (H), ubiG (H), truA (J), edd (GE), normM (V), murB (M), rlmD (H), ytfH (K), RTX toxin-related (Q), eda (G), asd (E), prmC (J), bioH (I), pncA (Q), lcbA (M), rlpA (M), yibK (J), thiamin transport (P), hexR (K), transpeptidase (M), nudE (L), trxA (O), valS (J), tsaC (J), recC (L), ompA (JM), adhesin (W)* |
| | ***Anchor 2*** (Viewpoint 2; contains *parS1-6*) | |
| | **Genomic position** (Mbp; bins) | 1.245–1.270; 251–254 |
| | **Genes** | *ruvA (L), hemerythrin-like (P), heptosyltransferase (M), etfB (C), etfA (C), ltaE (E), purD (F), ldh (C)* |
| **Exponential Loop 2; Exp L2;** anchor 3 + anchor 3') | ***Anchor 3'*** | |
| | **Genomic position (**Mbp; bins) | 1.020–1.135; 204–227 |
| | **Genes** | *had (O), hisG (E), cdsA (M), dxr (I), proC (E), hisD (E), dtd (J), nagK (Q), truC (J), nnrD (H), gtrA (U), yfdH (M), transposase (L), polar aa transport permease (P), ribF (H), ubiG (H), truA (J), edd (GE), norM (V), murB (M), rlmD (H), ytfH (K), RTX toxin-related (Q), eda (G), asd (E), prmC (J), bioH (I), pncA (Q), lcbA (M), rlpA (M), yibK (J), thiamin transport (P), hexR (K), transpeptidase (M), nudE (L), trxA (O), valS (J), tsaC (J), recC (L), ompA (JM), adhesin (W)* |
| | ***Anchor 3*** (Viewpoint 3) | |
| | **Genomic position** (Mbp; bins) | 1.285–1.290; 257–258 |
| | **Genes** | *csy1 (L), cas3 (L), cas1 (L), lldF (C)* |
| **Stationary Loop 1 (Stat L1;** anchor 2 + anchor 2') | ***Anchor 2'*** | |
| | **Genomic position** (Mbp; bins) | 1.030–1.135; 207–227 |
| | **Genes** | *proC (E), hisD (E), dtd (J), nagK (Q), truC (J), nnrD (H), gtrA (U), yfdH (M), transposase (L), polar aa transport permease (P), ribF (H), ubiG (H), truA (J), edd (GE), norM (V), murB (M), rlmD (H), ytfH (K), RTX toxin-related (Q), eda (G), asd (E), prmC (J), bioH (I), pncA (Q), lcbA (M), rlpA (M), yibK (J), thiamin transport (P), hexR (K), transpeptidase (M), nudE (L), trxA (O), valS (J), tsaC (J), recC (L), ompA (JM), adhesin (W)* |
| | ***Anchor 2*** (Viewpoint 2; contains *parS1-6*) | |
| | **Genomic position** (Mbp; bins) | 1.245–1.270; 251–254 |
| | **Genes** | *ruvA (L), hemerythrin-like (P), heptosyltransferase (M), etfB (C), etfA (C), ltaE (E), purD (F), ldh (C)* |
| **Stationary Loop 2 (Stat L2;** anchor 3 + anchor 3') | ***Anchor 3'*** | |
| | **Genomic position** (Mbp; bins) | 1.065–1.090; 213–218 |
| | **Genes** | *ytfH (K), RTX toxin-related (Q), eda (G), asd (E), prmC (J), bioH (I), pncA (Q), lcbA (M), CDP-glycerol glycerophosphotransferase (M)* |
| | ***Anchor 3*** (Viewpoint 3) | |
| | **Genomic position** (Mbp; bins) | 1.285–1.290; 257–258 |
| | **Genes** | *csy1 (L), cas3 (L), cas1 (L), lldF (C)* |

| | | |
|---|---|---|
| **Solid Loop (Solid L;** anchor 1 + anchor 1’) | ***Anchor 1’*** | |
| | **Genomic position** (Mbp; bins) | 1–1.005; 200–201 |
| | **Genes** | *dadA (C)*, *serS (J)*, *transposase (L)*, *tetR (K)*, *dusA (H)* |
| | ***Anchor 1*** (Viewpoint 1) | |
| | **Genomic position** (Mbp; bins) | 1.135–1.145; 227–229 |
| | **Genes** | *adhesin (W)*, *argF (E), pdxA (H), ftsY (U)*, *cbiM (H)* |

**Table S7. Genes in loop anchors in liquid exponential, liquid stationary or solid medium *A. filiformis*.**

| Gene | Liquid Exponential | Liquid Stationary | Solid medium |
|---|---|---|---|
| Adhesin | ✓ | ✓ | ✓ |
| Transposase | ✓ | ✓ | ✓ |
| *recC/ruvA* | ✓ | ✓ | – |
| *cas* genes | ✓ | ✓ | – |
| *murB-lcbA* region | ✓ | ✓ | – |
| *tetR* | – | – | ✓ |
| *dusA* | – | – | ✓ |
| *ftsY* | – | – | ✓ |
| *cbiM* | – | – | ✓ |

**Table S8.** DNA FISH probe primers.

| Target | Probe No. | Primer | Sequence (5‘-3‘) | Product (nt) |
|---|---|---|---|---|
| ***A. filiformis ori*** (9,232 nt) | 1 | Af_Ori1F | GGTTTTCAATGCCGTATCT | 300 |
| | | Af_Ori1R | GACAATTTGTCCAAAGATGAGG | |
| | 2 | Af_Ori2F | CGCAGCCAATTCAATATGAC | 300 |
| | | Af_Ori2R | CAACAAGCAGGTTCAGAAG | |
| | 3 | Af_Ori3F | GTGGTGCAGTTTCAGATAGT | 300 |
| | | Af_Ori3R | AAACTCTACAACGGTAGCG | |
| | 4 | Af_Ori4F | TGTCAAACCAATCATGACG | 300 |
| | | Af_Ori4R | AAGACACGTCTTTGCGTAT | |
| | 5 | Af_Ori5F | GCAATCACATAGCGACCTT | 300 |
| | | Af_Ori5R | AACCAATCCCTTGTCTGAA | |
| | 6 | Af_Ori6F | TTTAAGCCAGCCTTTTGGAT | 300 |
| | | Af_Ori6R | GGATTTTGAGTTTGACCCG | |
| | 7 | Af_Ori7F | GCTTCAGCTTGAGATACTG | 300 |
| | | Af_Ori7R | AGACATTTTGGAAGCGGTAA | |
| | 8 | Af_Ori8F | TGGAATAGAAGCCAACTCAG | 300 |
| | | Af_Ori8R | TATCGCGGTATCAGTGTTG | |
| | 9 | Af_Ori9F | ACGCTAGATACATCTACGC | 300 |
| | | Af_Ori9R | GATTTTGGGTCCTCGTGG | |
| | 10 | Af_Ori10F | ACACATCTACTGATTCGTCA | 300 |
| | | Af_Ori10R | CAGTTCCAATTCTTTGACCAAC | |

| | | | | |
|---|---|---|---|---|
| ***A. filiformis ter*** (6,778 nt) | 1 | Af_Ter1F | TTGGTTTACCCCATTAGCG | 300 |
| | | Af_Ter1R | CATGTCAAGGCAGCTTTG | |
| | 2 | Af_Ter2F | ATGTGGCAGCAATTTCAG | 300 |
| | | Af_Ter2R | GAACACGAATTGCCCTTTTA | |
| | 3 | Af_Ter3F | ACGCAAATACGCCAAAAAC | 300 |
| | | Af_Ter3R | TACATTTGGGGCGACAAA | |
| | 4 | Af_Ter4F | AACTTTGCACCACATATTCG | 300 |
| | | Af_Ter4R | GCCGTTTTGTTCAAGTGT | |
| | 5 | Af_Ter5F | TTATAATGCGCCCAACAATG | 300 |
| | | Af_Ter5R | GCACTTCGCATTAAAAACAC | |
| | 6 | Af_Ter6F | TGACGACAAAGTGGAACG | 300 |
| | | Af_Ter6R | AACAACCACCAAGCCAAT | |
| | 7 | Af_Ter7F | CCTGAAACAAAATCCGCC | 300 |
| | | Af_Ter7R | ATTCATGCGGATTTTGACC | |
| | 8 | Af_Ter8F | TGGAAGAGTCCACCAGTT | 300 |
| | | Af_Ter8R | AAAACAACCACATTGGCG | |
| | 9 | Af_Ter9F | GCAACAACACGCCATTTT | 300 |
| | | Af_Ter9R | TTCCCCTGCCCCATTATTA | |
| | 10 | Af_Ter10F | CGCACAGCAATTCAAAATCT | 300 |
| | | Af_Ter10R | GTGTGAATGTGATACAAGCG | |

| | | | | |
|---|---|---|---|---|
| ***A. filiformis* left arm** (6,718 nt) | 1 | Af_leftarm1F | TGCAGGTTTAGGAACATCTAC | 300 |
| | | Af_leftarm1R | GCTGAAATTCCGGGTTTG | |
| | 2 | Af_leftarm2F | GATGGCAACCAAGTCTTTATC | 300 |
| | | Af_leftarm2R | ATGGTGTTGTGTATCTGCA | |
| | 3 | Af_leftarm3F | AACCTGTATTGGGAATGGAAC | 300 |
| | | Af_leftarm3R | AAGCCCTGATTATTGAGTCG | |
| | 4 | Af_leftarm4F | CACTCGGCTCACTTCCG | 303 |
| | | Af_leftarm4R | CTAGACATCAGCCCCAGC | |
| | 5 | Af_leftarm5F | TTTATTCTTCTGATGGCGTG | 282 |
| | | Af_leftarm5R | AGGCTTCATTGAATTTGCG | |
| | 6 | Af_leftarm6F | CACTTTGACCAAAGACCAAA | 300 |
| | | Af_leftarm6R | CGTTCGTAAATCAATTCGGC | |
| | 7 | Af_leftarm7F | GCGCAATTTTCCATCAAAAG | 300 |
| | | Af_leftarm7R | TGGGTAACATAAGGACGGT | |
| | 8 | Af_leftarm8F | GGCTACGATTTCCACAGTAA | 300 |
| | | Af_leftarm8R | GAAACCAAAACGTTGCACA | |
| | 9 | Af_leftarm9F | GCGCTGGAAAAATTGCAA | 314 |
| | | Af_leftarm9R | TTGCCCACATAATCTTGTG | |
| | 10 | Af_leftarm10F | TCAAATCCACATCGCGC | 316 |
| | | Af_leftarm10R | TTTTTCTTGCGCGTGATG | |

| | | | | |
|---|---|---|---|---|
| ***A. filiformis* right arm** (7,846 nt) | 1 | Af_rightarm1F | ACGATAAACCTTGATGTTGC | 300 |
| | | Af_rightarm1R | TTGATGATGCGCTTGAAAC | |
| | 2 | Af_rightarm2F | AAAATCGGTCAAAATAGCCC | 300 |
| | | Af_rightarm2R | CAAACGCATTATCCCAATCTC | |
| | 3 | Af_rightarm3F | GCAGCCTGAAAATTATTCG | 300 |
| | | Af_rightarm3R | TTGCAAAACGTGGACATC | |
| | 4 | Af_rightarm4F | CAAACGTGGCAAATTTCG | 317 |
| | | Af_rightarm4R | GGCACCAAATAGCGTACC | |
| | 5 | Af_rightarm5F | AATGTCGCTGATTGGTTTG | 300 |
| | | Af_rightarm5R | GCCAAACTCAATGCAAACA | |
| | 6 | Af_rightarm6F | GCAGGCGAATTTGGTTTT | 300 |
| | | Af_rightarm6R | GTGGTTGTCAATCACGGTA | |
| | 7 | Af_rightarm7F | CTGAAACGGCTTTTTAGGTG | 300 |
| | | Af_rightarm7R | CAGCCGACTTAACCCAAG | |
| | 8 | Af_rightarm8F | GAATAAACGCAGGGTACG | 283 |
| | | Af_rightarm8R | ATGAACACAATTTGCCCG | |
| | 9 | Af_rightarm9F | TTTGCCAAAATCGGGCG | 280 |
| | | Af_rightarm9R | CAAACAAAATCACGCCAAAT | |
| | 10 | Af_rightarm10F | ACGGCTTTGAATGCTTTG | 286 |
| | | Af_rightarm10R | GCGTATTGCCAAAATCCAC | |

| | | | | |
|---|---|---|---|---|
| ***S. muelleri ori*** (8,158 nt) | 1 | Sm_ori_1F | CAATGTAGATGCCGCTGAA | 300 |
| | | Sm_ori_1R | CAACGATAGGTCACATTTGC | |
| | 2 | Sm_ori_2F | GATTTAGCAGCAGAACAACC | 300 |
| | | Sm_ori_2R | GCCACAAATATCCACTACGT | |
| | 3 | Sm_ori_3F | CTATATATTCATTCGCTGGACG | 300 |
| | | Sm_ori_3R | TTACCGCTTTTACGTGGG | |
| | 4 | Sm_ori_4F | ATTGATTACGTTGGCTCATG | 300 |
| | | Sm_ori_4R | TTTTACGAGCAAGCTGTCT | |
| | 5 | Sm_ori_5F | GTGCTGATTTACGACGAAG | 300 |
| | | Sm_ori_5R | CACCAATTGAGAAGTACGC | |
| | 6 | Sm_ori_6F | CCAAACCCACTACACATAAT | 300 |
| | | Sm_ori_6R | GTTTTGGTGATCCTGAAAC | |
| | 7 | Sm_ori_7F | CAAAATTAGCCAGCGGAATC | 300 |
| | | Sm_ori_7R | TTTTGGGGCAGATTGATTC | |
| | 8 | Sm_ori_8F | TATTACCTAACTTTCGCTCCG | 300 |
| | | Sm_ori_8R | AAAAAGGAGTGTGTACCGA | |
| | 9 | Sm_ori_9F | TACTATCCTGCATAC CTGCA | 325 |
| | | Sm_ori_9R | AATTTGGTGGCTACGAATG | |
| | 10 | Sm_ori_10F | CATTTGTGCAGTTTGGTTAG | 300 |
| | | Sm_ori_10R | GTGAAAATGTCCATGAATCC | |

| | | | | |
|---|---|---|---|---|
| ***S. muelleri ter*** (8,187 nt) | 1 | Sm_ter_1F | GATTTGCGGTCAATCGTT | 300 |
| | | Sm_ter_1R | GGGCAAATTCAATTCAGACC | |
| | 2 | Sm_ter_2F | ATGAAACCATCTGAACCGT | 300 |
| | | Sm_ter_2R | CATTGATGCACTGGTTATTG | |
| | 3 | Sm_ter_3F | GATGATTTCACGGATTTTAGCC | 300 |
| | | Sm_ter_3R | CGTTCTCAATTTGTAGACCAAG | |
| | 4 | Sm_ter_4F | TCGTTAATTCAGGCGTGTC | 300 |
| | | Sm_ter_4R | GGACGGTTGGACTTATTCG | |
| | 5 | Sm_ter_5F | TTGTCCGTCTAAATAAGGTGTC | 300 |
| | | Sm_ter_5R | TTGGACAAACTCAAACATGG | |
| | 6 | Sm_ter_6F | CACCGAATGATCAGGAATG | 300 |
| | | Sm_ter_6R | GGTTTAAAAGCGAGAGACG | |
| | 7 | Sm_ter_7F | TTCCTTGTGGGTCTAATACTTG | 300 |
| | | Sm_ter_7R | CAGAGCCATTAAGTAAATCACG | |
| | 8 | Sm_ter_8F | AGCATCAACACGTACGAAA | 300 |
| | | Sm_ter_8R | GATTATCGGCATGAGACGG | |
| | 9 | Sm_ter_9F | GCGTAGGATTATTCTCATTAGC | 314 |
| | | Sm_ori_10R | GTGAAAATGTCCATGAATCC | |

| | | | | |
|---|---|---|---|---|
| ***S. muelleri 5x parS_c*** (9,335 nt) | 1 | Sm_parS1F | AATACCCATCCTACGTCTGA | 300 |
| | | Sm_parS1R | CTCTTGTTGGCGTTCTAAAATC | |
| | 2 | Sm_parS2F | TTATGCACGTGAGATTGAC | 300 |
| | | Sm_parS2R | GCATTACGCAACAAGTCAT | |
| | 3 | Sm_parS3F | CAACATCACAAAGTACCAGTAC | 300 |
| | | Sm_parS3R | CTACGGCTGTAGGTGATAAT | |
| | 4 | Sm_parS4F | TGGATATTCCTGTGGTACCA | 300 |
| | | Sm_parS4R | CTGGGTCATAAGTGTAATTCAC | |
| | 5 | Sm_parS5F | GTAAAGCTAAAGACCACACAG | 300 |
| | | Sm_parS5R | GTACACCTGTAACAGTTGT | |
| | 6 | Sm_parS6F | AGGCAATGTAACGACAGAAG | 300 |
| | | Sm_parS6R | GTAACTGTGTAAACCGCCTTA | |
| | 7 | Sm_parS7F | GGTAATACCAAAGGCACAAATC | 300 |
| | | Sm_parS7R | CGGTTCTAATTGCAACGTC | |
| | 8 | Sm_parS8F | AGTTCTGGCGTGACTTTATT | 300 |
| | | Sm_parS8R | AACTTTACCAGTCGTGTAAC | |
| | 9 | Sm_parS9F | TACTGATAGTTTACCAGTAGCG | 300 |
| | | Sm_parS9R | AATTTTGCTGTTGAGGTATCG | |

| | | | | |
|---|---|---|---|---|
| ***Ca. T. hypermnestrae ori*** (2,858 nt) | 1 | Th_dnaA_start | TCCGTGGAAATCTGGAATCAATGT | 306 |
| | | Th_ori_1R | TGTGGATTTTTCTTGAGGTGTGG | |
| | 2 | Th_ori_2F | CCGCGAGCAACTTAATTACCG | 344 |
| | | Th_ori_2R | GCCTGCGAAGAACTGAATATCA | |
| | 3 | Th_ori_3F | TCGAATCCAAACAGCAAATCGT | 315 |
| | | Th_ori_3R | GTACGTTTCTGGCAAATTCGAG | |
| | 4 | Th_ori_4F | GGACATGCTTGCTGCTCAG | 315 |
| | | Th_ori_4R | CATTATGCTGTCAGGGTTCTGAAC | |
| | 5 | Th_ori_5F | TTCAGTGCCAGCAATTTCGCC | 310 |
| | | Th_ori_5R | CCGGATAAAGTCGTCCTTTGCC | |
| | 6 | Th_ori_6F | CGGAACCGATCTAGAAATAGAGG | 304 |
| | | Th_ori_6R | CGCCATCGAAAATGCAGTCTT | |
| | 7 | Th_ori_7F | CAGGACGTTCGCTATTATCT | 334 |
| | | Th_ori_7R | CATAGGCAACTTGCTCCGGA | |
| | 8 | Th_ori_8F | AGATACACTGAAGCAGGCATTG | 317 |
| | | Th_ori_8R | GTCTCATCGGCATTACTACAAAG | |
| | 9 | Th_ori_9F | GACACATGATTTTGAAGGTAGAG | 216 |
| | | Th_dnaA_stop | CAGCCTCTCAGTGTCACCGT | |

| | | | | |
|---|---|---|---|---|
| ***Ca. T. hypermnestrae ter*** (3,239 nt) | 1 | Th_dif_F1 | GTAAGGGGATTGCCAGGCTTG | 311 |
| | | Th_dif_R1 | GTGGACGACACGGTGCATTT | |
| | 2 | Th_dif_F2 | CACGATGCCCAGCGTCATG | 333 |
| | | Th_dif_R2 | CTCTCGACCAAGGAGGTACTTGC | |
| | 3 | Th_dif_F3 | GTCGCCCGCTTGTCGATATCG | 314 |
| | | Th_dif_R3 | GAGAACCTGACCGGCGTTTATT | |
| | 4 | Th_dif_F4 | ATGAAATCGGTATCCGTACGG | 318 |
| | | Th_dif_R4 | GAATTTCTCCGAGGCGCCG | |
| | 5 | Th_dif_F5 | AGCGTCAGAAACCCGATGG | 323 |
| | | Th_dif_R5 | CTGCTGATCATTGTGGTCGCC | |
| | 6 | Th_dif_F6 | GCCAGCGGCAGGAGATGC | 324 |
| | | Th_dif_R6 | GGTTGTCGATCTGATTGAGGACCC | |
| | 7 | Th_dif_F7 | GTCAGCGCATCCCCCTCG | 319 |
| | | Th_dif_R7 | GGCTGGTGATTGCAGGGACC | |
| | 8 | Th_dif_F8 | AGCGATGGCCCAAAATCCAAG | 305 |
| | | Th_dif_R8 | ATGATGTCCCCACGGGATAATTCG | |
| | 9 | Th_dif_F9 | CATGCAGACCTTTGTCGCGC | 330 |
| | | Th_dif_R9 | GGCTTCCTCCTTGATGTGCTCG | |
| | 10 | Th_dif_F10 | GTGCTGGCGGCGATGGGA | 326 |
| | | Th_dif_R10 | GCCTGTGCTTAAGTGGCTGGT | |
| | 11 | Th_dif_F11 | ACCGCGATCACGCGCTGA | 325 |
| | | Th_dif_R11 | AAACCGGAACCTGCTGACCA | |
| | 12 | Th_dif_F1.2 | CGGATTACGCGGATGGGGTAA | |
| | | Th_dif_R11.2 | TATTAGCAGATTGCGACGCGCG | |

| | | | | |
|---|---|---|---|---|
| ***Ca. T. hypermnestrae parS2*** (3,524 nt) | 1 | Th_parS2_1f | CTCTACCTGTTTGATCGCGTTCG | 319 |
| | | Th_parS2_1r | CAAGGTCTTCAGCGAGATGTACG | |
| | 2 | Th_parS2_2f | TCGGCGGATTCGAGCCACA | 290 |
| | | Th_parS2_2r | CTGAGTCGTCTCCTCTCCATGG | |
| | 3 | Th_parS2_3f | GAGCCTCCCAGCATCATTTCC | 313 |
| | | Th_parS2_3r | ATGCCGGCCATCAGGTCTTC | |
| | 4 | Th_parS2_4f | CCATCGGGTCCGTCCAGGA | 321 |
| | | Th_parS2_4r | CGCTTGTTTCGAGCCTCTATTCC | |
| | 5 | Th_parS2_5f | GTTATTTATACTACAGCGCCGAGCG | 330 |
| | | Th_parS2_5r | GCCAGGTTTTTCGCCACGG | |
| | 6 | Th_parS2_6f | CGAGGACAAAGCCAAGGTCG | 316 |
| | | Th_parS2_6r | GGTGAGGAAGGTGCCGATG | |
| | 7 | Th_parS2_7f | GGATGGCCGACAGCGGTAAG | 305 |
| | | Th_parS2_7r | AATGACGTGCTTGATGGTGCC | |
| | 8 | Th_parS2_8f | AACGACTCGATCGACGGCA | 313 |
| | | Th_parS2_8r | GATCTTAATCATGCGCGATTCTCC | |
| | 9 | Th_parS2_9f | GACGACCACAAGCTGGTACG | 320 |
| | | Th_parS2_9r | GCCTCGATCATCTCCTCGG | |
| | 10 | Th_parS2_10f | GATCCGCTCGGTCGTCAGTG | 311 |
| | | Th_parS2_10r | CCGCTCAGGCATCCTCCTC | |
| | 11 | Th_parS2_11f | GGATGTCCCGGATTGTGCGG | 333 |
| | | Th_parS2_11r | CTGAGCAGGACGTTATACCGG | |

| | | | | |
|---|---|---|---|---|
| **Ca. T. hypermnestrae *ftsQAZ*** (3,690 nt) | 1 | Th_ftsQAZ_F1.2 | CTGCCCCTGATCCGCCTGGA | 335 |
| | | Th_ftsQAZ_R1.1 | AATGGTTCAAGGCTGGTCTCC | |
| | 2 | Th_ftsQAZ_F2.1 | CCGGACAAGCGATGAACAGG | 329 |
| | | Th_ftsQAZ_R2.1 | CAGGCGATCCGGCCAGTGA | |
| | 3 | Th_ftsQAZ_F3.1 | GAGCTCCGGGTGGTGGAAC | 323 |
| | | Th_ftsQAZ_R3.1 | GGGTAGACCCGGATAAACCTGGC | |
| | 4 | Th_ftsQAZ_F4.1 | CCGTCGGTGATCGATATGCG | 319 |
| | | Th_ftsQAZ_R4.1 | TCCTTCACCTCGCCCACCAG | |
| | 5 | Th_ftsQAZ_F5.1 | CGACGAGGGGATCGAGATTATCG | 312 |
| | | Th_ftsQAZ_R5.1 | TGGGGCAGGATGTGCAGGAT | |
| | 6 | Th_ftsQAZ_F6.1 | AATTCATCATCGACAATCAGGAAGGG | 308 |
| | | Th_ftsQAZ_R6.1 | GCCGGCGATGGGAATCACG | |
| | 7 | Th_ftsQAZ_F7.1 | GATCAGGTGACCAACGATATCGCC | 286 |
| | | Th_ftsQAZ_R7.1 | AGCTGCCGCCGGTCATGAC | |
| | 8 | Th_ftsQAZ_F8.1 | CCAAGATGGCGGGGCTCAT | 361 |
| | | Th_ftsQAZ_R8.1 | CCGTTTTGCTCGATGGTTATCTGATT | |
| | 9 | Th_ftsQAZ_F9.1 | ATGTTTGAACTCATGGATACCAACGG | 397 |
| | | Th_ftsQAZ_R9.1 | TCACTACCGCCACGGTCAGG | |
| | 10 | Th_ftsQAZ_F10.1 | CCAAGCCCTTTGCGTTCGAG | 349 |
| | | Th_ftsQAZ_R10.1 | AAGGGGCTGTTGATGGCCG | |
| | 11 | Th_ftsQAZ_F11.1 | GCTGGAGGATATCGACCTGGC | 336 |
| | | Th_ftsQAZ_R11.1 | CCCTTGTTGCGATAGACCGCC | |

| | | | | |
|---|---|---|---|---|
| ***C. steedae ori*** (7,353 nt) | 1 | C_Ori1F | TCAATTTCAAAAACCGGCTC | 300 |
| | | C_Ori1R | GATAGAACACATTATGTCTGCG | |
| | 2 | C_Ori2F | GCATGAGTGTGAAAGTAGC | 300 |
| | | C_Ori2R | AAAAACCTGTGCATTCCAG | |
| | 3 | C_Ori3F | AACTGTGGTGTTCAACGTTA | 300 |
| | | C_Ori3R | AGTTTGCCTTTCAGTTCAG | |
| | 4 | C_Ori4F | GTAGCCACGGGTTCATTGA | 331 |
| | | C_Ori4R | ACCACTTAGCCTAATTATCGG | |
| | 5 | C_Ori5F | AGGTTAAACAGTCAGTGGTAG | 300 |
| | | C_Ori5R | TTTTTCGATCAAAGACGGG | |
| | 6 | C_Ori6F | TTTACCGAGCAATATGTCG | 300 |
| | | C_Ori6R | GCGTACAAAGCGGTTAATC | |
| | 7 | C_Ori7F | GTGGTGCTGATTTACGATGA | 300 |
| | | C_Ori7R | CAGACGTTGACTCAATGTG | |
| | 8 | C_Ori8F | GCGTCCCACGGATAATTAAT | 300 |
| | | C_Ori8R | CGAATATTACGCCAACGAAAG | |
| | 9 | C_Ori9F | GCTTACCATACCTTCAAGC | 300 |
| | | C_Ori9R | CGGCAAAAAGAAAGTTTGAG | |
| | 10 | C_Ori10F | GTTAGACATAAACCCGACATGA | 300 |
| | | C_Ori10R | CACCATGCTACTTGGTCTG | |
| | 11 | C_Ori11F | AGAAAGGAAACCCCATGTC | 300 |
| | | C_Ori11R | GATTAAAGGTGGTAAATTCGCG | |

| | | | | |
|---|---|---|---|---|
| ***C. steedae ter*** (4,498 nt) | 1 | C_Ter1F | TTGCCCGACAACATTACTAC | 300 |
| | | C_ Ter1R | CTGTTTGCGGTTCATATAGG | |
| | 2 | C_Ter2F | GAAGGTTTTGAACGCATTCA | 300 |
| | | C_Ter2R | CGAACTTTACACCTTTCCTTC | |
| | 3 | C_Ter3F | CGTAGATTTAAACGCTGACCT | 300 |
| | | C_Ter3R | TAAGCCATTTCCACACTCG | |
| | 4 | C_Ter4F | GCCGACAGTATTTGCTTAC | 300 |
| | | C_Ter4R | CTTTGTGTTGCCCGATATAATC | |
| | 5 | C_Ter5F | CCAAAGAGCAACTGGGTAT | 300 |
| | | C_Ter5R | TGTCCAGTTGAAAATCTTCC | |
| | 6 | C_Ter6F | GCTTACCTTATTAGATGTGG | 300 |
| | | C_Ter6R | GTAGGTTTTGTAGGGCTTT | |
| | 7 | C_Ter7F | GGTGGCGTATTGCTTAACT | 300 |
| | | C_Ter7R | GTGTAAAGGTTCTAAATGGGA C | |
| | 8 | C_Ter8F | ACACCTTTTTATCGGTAGC | 300 |
| | | C_Ter8R | TCGTCAAGGGGAATATCAG | |
| | 9 | C_Ter9F | AACAATGTGGACAAGCACT | 300 |
| | | C_Ter9R | TACTGTGGGGGTCTAGTTG | |
| | 10 | C_Ter10F | CGGAATTACAGGTATCACAAC | 300 |
| | | C_Ter10R | CAAGATGTTCCTTTAACCAGTG | |

**Table S9.** PCR cycling conditions for DNA FISH probe production.

| Target | Primer | Product (nt) | Cycling conditions |
|---|---|---|---|
| *A. filiformis ori* | see Table S8 | Initial 3,622 nt-long fragment | 98°C for 5 min, followed by 31 cycles at 98°C for 20 s, 64°C (-0.3°C every cycle) for 20 s, 72°C for 120 s, followed by a final elongation step at 72°C for 5 min. |
| | see Table S8 | ~300 nt-long probes (P1-P11) | 98°C for 5 min, followed by 31 cycles at 98°C for 20 s, 64°C (-0.3°C every cycle) for 20 s, 72°C for 15 s, followed by a final elongation step at 72°C for 5 min. |
| *A. filiformis ter* | see Table S8 | Initial 3,583 nt-long fragment | 98°C for 5 min, followed by 31 cycles at 98°C for 20 s, 63°C (-0.3°C every cycle) for 20 s, 72°C for 120 s, followed by a final elongation step at 72°C for 5 min. |
| | see Table S8 | ~300 nt-long probes (P1-P11) | 98°C for 5 min, followed by 31 cycles at 98°C for 20 s, 62°C (-0.3°C every cycle) for 20 s, 72°C for 15 s, followed by a final elongation step at 72°C for 5 min. |
| *A. filiformis* left arm | see Table S8 | Initial 6,718 nt-long fragment | 98°C for 5 min, followed by 31 cycles at 98°C for 20 s, 63°C (-0.3°C every cycle) for 20 s, 72°C for 120 s, followed by a final elongation step at 72°C for 5 min. |

| | | | |
|---|---|---|---|
| | see Table S8 | ~300 nt-long probes (P1-P10) | 98°C for 5 min, followed by 31 cycles at 98°C for 20 s, 63°C (-0.3°C every cycle) for 20 s, 72°C for 30 s, followed by a final elongation step at 72°C for 5 min. |
| *A. filiformis* right arm | see Table S8 | Initial 7,846 nt-long fragment | 98°C for 5 min, followed by 31 cycles at 98°C for 20 s, 63°C (-0.3°C every cycle) for 20 s, 72°C for 120 s, followed by a final elongation step at 72°C for 5 min. |
| | see Table S8 | ~300 nt-long probes (P1-P10) | 98°C for 5 min, followed by 31 cycles at 98°C for 20 s, 63°C (-0.3°C every cycle) for 20 s, 72°C for 30 s, followed by a final elongation step at 72°C for 5 min. |
| *S. muelleri ori* | see Table S8 | Initial 8,158 nt-long fragment | 98°C for 5 min, followed by 31 cycles at 98°C for 20 s, 58°C (-0.3°C every cycle) for 20 s, 72°C for 240s, followed by a final elongation step at 72°C for 5 min. |
| | see Table S8 | ~300 nt-long probes (P1-P10) | 98°C for 5 min, followed by 31 cycles at 98°C for 20 s, 58°C (-0.3°C every cycle) for 20 s, 72°C for 15s, followed by a final elongation step at 72°C for 5 min. |
| *S. muelleri ter* | see Table S8 | Initial 8,187 nt-long fragment | 98°C for 5 min, followed by 31 cycles at 98°C for 20 s, 58°C (-0.3°C every cycle) for 20 s, 72°C for 240s, followed by a final elongation step at 72°C for 5 min. |

| | | | |
|---|---|---|---|
| | see Table S8 | ~300 nt-long probes (P1-P9) | 98°C for 5 min, followed by 31 cycles at 98°C for 20 s, 58°C (-0.3°C every cycle) for 20 s, 72°C for 15s, followed by a final elongation step at 72°C for 5 min. |
| *S. muelleri 5xparS_c* | see Table S8 | Initial 9,335 nt-long fragment | 98°C for 5 min, followed by 31 cycles at 98°C for 20 s, 63°C (-0.3°C every cycle) for 20 s, 72°C for 120 s, followed by a final elongation step at 72°C for 5 min. |
| | see Table S8 | ~300 nt-long probes (P1-P9) | 98°C for 5 min, followed by 31 cycles at 98°C for 20 s, 63°C (-0.3°C every cycle) for 20 s, 72°C for 30 s, followed by a final elongation step at 72°C for 5 min. |
| *C. steedae ori* | see Table S8 | Initial 7,353 nt-long fragment | 95°C for 5 min, followed by 31 cycles at 95°C for 20 s, 58°C (-0.3°C every cycle) for 20 s, 72°C for 270s, followed by a final elongation step at 72°C for 5 min. |
| | see Table S8 | 300 nt-long probes (P1-P11) | 95°C for 5 min, followed by 31 cycles at 95°C for 20 s, 58°C (-0.3°C every cycle) for 20 s, 72°C for 30s, followed by a final elongation step at 72°C for 5 min. |
| *C. steedae ter* | see Table S8 | Initial 4, 498 nt-long fragment | 95°C for 5 min, followed by 31 cycles at 95°C for 20 s, 58°C (-0.3°C every cycle) for 20 s, 72°C for 270s, followed by a final elongation step at 72°C for 5 min. |

| | | | |
|---|---|---|---|
| | see Table S8 | 300 nt-long probes (P1-P10) | 95°C for 5 min, followed by 31 cycles at 95°C for 20 s, 58°C (-0.3°C every cycle) for 20 s, 72°C for 30s, followed by a final elongation step at 72°C for 5 min. |
| *Ca.* T. hypermnestrae *ori* | see Table S8 | Initial 2,858 nt-long fragment | 95°C for 5 min, followed by 31 cycles at 98°C for 45 s, 57°C for 45 s, 72°C for 180 s, followed by a final elongation step at 72°C for 5 min. |
| | see Table S8 | ~300 nt-long probes (P1-P9) | 95°C for 5 min, followed by 31 cycles at 98°C for 45 s, 57°C for 45 s, 72°C for 30 s, followed by a final elongation step at 72°C for 5 min. |
| *Ca.* T. hypermnestrae *ter* | see Table S8 | Initial 3,239 nt-long fragment | 98°C for 5 min, followed by 31 cycles at 98°C for 20 s, 63°C (-0.3°C every cycle) for 20 s, 72°C for 125 s, followed by a final elongation step at 72°C for 5 min. |
| | see Table S8 | ~300 nt-long probes (P1-P10) | 98°C for 5 min, followed by 31 cycles at 98°C for 20 s, 58°C for 20 s, 72°C for 15 s, followed by a final elongation step at 72°C for 5 min. |
| *Ca.* T. hypermnestrae *ftsQAZ* | see Table S8 | Initial 4,010 nt-long fragment | 98°C for 5 min, followed by 31 cycles at 98°C for 20 s, 65°C (-0.3°C every cycle) for 20 s, 72°C for 120 s, followed by a final elongation step at 72°C for 5 min. |

| | | | |
|---|---|---|---|
| | see Table S8 | ~300 nt-long probes (P1-P11) | 95°C for 5 min, followed by 31 cycles at 94°C for 60 s, 57°C for 30 s, 72°C for 30 s, followed by a final elongation step at 72°C for 5 min. |
| *Ca.* T. hypermnestrae *parS2* | see Table S8 | Initial 3,632 nt-long fragment | 95°C for 5 min, followed by 31 cycles at 98°C for 20 s, 64°C (-0.3°C every cycle) for 20 s, 72°C for 120 s, followed by a final elongation step at 72°C for 5 min. |
| | see Table S8 | ~300 nt-long probes (P1-P10) | 98°C for 5 min, followed by 31 cycles at 98°C for 20 s, 64°C (-0.2°C every cycle) for 20 s, 72°C for 20 s, followed by a final elongation step at 72°C for 5 min. |

**Table S10.** SRA accession numbers

| | Sample name | NCBI accession number |
|---|---|---|
| **Genome sequencing** | *A. filiformis* long reads | fast5: SRR13289602, fastq: SRR16893507 |
| | *A. filiformis* short reads | SRR15998661 |
| | *C. steedae*, long reads | fast5: SRR16896741, fastq: SRR16947409 |
| | *C. steedae,* short reads | SRR19164917 |
| | *A. filiformis* assembly | GCF_020162295.1 |
| | *C. steedae,* assembly | GCF_023547005.1 |
| **Marker frequency analysis** | *A. filiformis*, late exponential phase | SRR17498973 |
| | *A. filiformis*, stationary phase | SRR17498972 |
| | *C. steedae*, late exponential phase | SRR17498976 |
| | *C. steedae*, stationary phase | SRR17498975 |
| | *E. coli,* exponential phase* | ERR902995 |
| | *E. coli,* stationary phase* | ERR902996 |
| | *A. filiformis* 1kb-binned raw counts | GSE194131 |
| | *C. steedae* 1kb-binned raw counts | GSE205139 |
| **RNA-Seq** | *A. filiformis*, exponential phase (3 biological replicates) | SRR17498971, SRR17498970, SRR17498961 |
| | *A. filiformis*, stationary phase (3 biological replicates) | SRR17498969, SRR17498968, SRR17498954 |
| | *A. filiformis*, plate (3 biological replicates) | SRR17498958, SRR17498959, SRR17498960 |
| | *A. filiformis* raw count table | GSE194132 |
| | *C. steedae* raw count table | GSE250250 |
| | *C. steedae,* plate (3 biological replicates) | SRR17498948, SRR17498950, SRR17498939 |
| | *C. steedae,* exponential phase (3 biological replicates) | SRR17498951, SRR17498952, SRR17498953 |
| | *C. steedae,* stationary phase (3 biological replicates) | SRR17498945, SRR17498946, SRR17498947 |
| **3C-Seq** | *A. filiformis*, exponential phase (replicate 1) | SRR19164944 |
| | *A. filiformis*, exponential phase (replicate 2) | SRR19164943 |
| | *A. filiformis*, stationary phase | SRR19164942 |
| | *A. filiformis*, plate (replicate 1) | SRR27205587 |
| | *A. filiformis*, plate (replicate 2) | SRR27206238 |
| | *A. filiformis*, exponential phase with 25 µg/ml rifampicin for 30 min | SRR27206238 |

| | | |
|---|---|---|
| | *A. filiformis*, exponential phase with 25 µg/ml Chloramphenicol for 30 min | SRR27206240 |
| | *C. steedae*, plate | SRR27206419 |
| | *C. steedae*, exponential phase | SRR27206418 |
| | *C. steedae*, stationary phase | SRR27206420 |
| | *A. filiformis* 3C-seq matrices | GSE250400 |
| | *C. steedae* 3C-seq matrices | GSE250401 |
| **ChIP-seq** | *A. filiformis* ChIP-seq reads | SRX32354318, SRX32354319 |
| | *A. filiformis* ChIP-seq counts | GSE322725 |
| | *S. muelleri* ChIP-seq reads | SRX32354384, SRX32354385 |
| | *S. muelleri* ChIP-seq counts | GSE322727 |

* From Ivanova et al., 2015
